# A microenvironment-driven, HLA-II-associated insulin neoantigen elicits persistent memory T cell activation in diabetes

**DOI:** 10.1101/2024.11.07.622538

**Authors:** Neetu Srivastava, Anthony N. Vomund, Rongzhen Yu, Orion J. Peterson, Yuqing Yang, David P. Turicek, Omar Abousaway, Tiandao Li, Lisa Kain, Pamela Stone, Aisha Ansar, Cristina C. Clement, Siddhartha Sharma, Rima Melhem, Bo Zhang, Chang Liu, Alok V. Joglekar, Hao Hu, Chyi-Song Hsieh, Laura Campisi, Laura Santambrogio, Luc Teyton, Emil R. Unanue, Ana Maria Arbelaez, Cheryl F. Lichti, Xiaoxiao Wan

## Abstract

The evolving antigenic landscape of autoimmune diabetes reflects a dynamic failure to preserve self-tolerance. Yet, how novel neoantigens emerge in humans remains incompletely understood. Here, we designed an immunopeptidomics-based approach to probe HLA-II-bound, islet-derived neoepitopes in patients with type 1 diabetes (T1D). We uncovered a microenvironment-driven Cys→Ser transformation, conserved between mice and humans, that reshapes autoreactivity to insulin, the core β-cell antigen, at the single-residue level. This transformation, which we call “C19S,” arises from oxidative remodeling of insulin in stressed pancreatic islets and can also occur in inflammatory antigen-presenting cells, contributing to a feed-forward loop of neoepitope formation and presentation as diabetes progresses. Despite involving just one amino acid, C19S is specifically recognized by HLA-DQ8-restricted, register-specific CD4^+^ T cells that expand in individuals with T1D. These C19S-specific CD4^+^ T cells lack regulatory potential but acquire a poised central memory phenotype that persists at different disease stages. These findings reveal a distinct, microenvironment-driven route of neoantigen formation that fuels sustained autoreactivity in diabetes.

## INTRODUCTION

Type 1 diabetes (T1D) results from the autoimmune destruction of insulin-producing β cells in the pancreatic islets of Langerhans. Despite extensive study of native β-cell antigens, the mechanisms underlying the collapse of immune tolerance remain incompletely understood. Neoantigens are novel forms of self-peptides that escape immune tolerance and become targets of autoreactive T cells^1^. Several well-characterized mechanisms contribute to neoantigen formation in T1D. Post-translational modifications (PTMs) encompass a wide range of biochemical changes to β-cell proteins, such as citrullination, deamination, and carbonylation, which alter amino acid side chains and generate modified epitopes capable of breaking immune tolerance^2–11^. By contrast, hybrid insulin peptides (HIPs) represent a specific mechanism wherein insulin peptides fuse with fragments of other β-cell proteins, forming novel junctional sequences that are absent in the native proteome^12^. These studies have both advanced our understanding of T1D pathogenesis and established β-cell neoantigens as valuable biomarkers and therapeutic targets.

The diabetic autoimmune process is characterized by continuous epitope spreading, facilitating a dynamic antigenic landscape. Whether other mechanisms lead to the formation of neoantigens and their presentation to autoreactive T cells remains unanswered. In a broader context, cancer immunology studies have revealed that neoantigens can be generated through previously unrecognized mechanisms during chronic inflammation. While single nucleotide polymorphisms have long been recognized as a major source of tumor-specific neoepitopes, recent studies have described a new class of neoantigens, termed “substitutants,” which arise when amino acid depletion leads to tRNA misincorporation, replacing one residue with another^13,14^. These findings suggest that non-mutational mechanisms can introduce single amino acid substitutions in self-proteins and create immunogenic neoepitopes. This concept is relevant to autoimmune diseases, where self-tissues typically harbor few somatic mutations. Moreover, since extra-genomic neoantigens cannot be detected by transcriptomic approaches like RNA sequencing, their discovery relies on mass spectrometry (MS)-based immunopeptidomics, with specialized searches required to confirm their often rare presence in the self-proteome or peptidome^15^.

Pancreatic β cells are intrinsically prone to oxidative stress and inflammation and are further exposed to several pathological signals throughout diabetes development. These conditions are thought to create a conducive tissue microenvironment that fosters the generation of diverse neoantigens^16–18^. Although studies using non-obese diabetic (NOD) mice have provided valuable insights, identifying HLA-II-presented neoepitopes derived from human islets in vivo remains a challenge.

In this study, we applied a β-cell degranulation strategy coupled with HLA-II immunopeptidome profiling to systematically identify islet-derived peptides presented in human peripheral blood. Unexpectedly, we detected a group of HLA-II-bound insulin B-chain peptides harboring a cysteine-to-serine (Cys→Ser) transformation at the 19^th^ position (C19S). Unlike conventional PTMs and HIPs that modify pre-existing peptides or join two canonical peptide segments, respectively, C19S directly alters the sequence of post-synthetic insulin at a single residue. We developed functional assays that localized C19S to post-synthetic insulin products within β-cell granules, a process distinct from formation of substitutants in cancer, which arises at the tRNA level during translation^13,14^. Mechanistically, C19S transformation is driven by oxidative stress in the inflammatory islet microenvironment. These distinctive features of C19S prompted us to further investigate its presentation by diabetes-associated MHC-II molecules and the phenotype of its neoepitope-specific CD4⁺ T cells in both NOD mice and humans with T1D.

## RESULTS

### Identification of insulin peptides with a Cys→Ser transformation in human and mouse MHC-II peptidomes

To probe potential HLA-II-bound neoepitopes derived from human islets in vivo, we used a β-cell degranulation strategy combined with immunopeptidome analysis in human PBMCs. This approach stems from our previous mouse studies, which showed that in vivo glucose stimulation triggers exocytosis of β-cell-derived peptides from pancreatic islets into circulation^19,20^. These released peptides are transiently presented by MHC-II-expressing antigen-presenting cells (APCs) in the blood within 30 to 120 minutes following glucose injection, providing a window for detecting rare islet-derived peptides in the self MHC-II peptidome of PBMCs^20^. We reasoned that this approach might circumvent the practical challenge of obtaining islet samples from T1D patients and enable the identification of neoepitopes potentially formed in inflamed human islets.

To apply this strategy in humans, we recruited T1D subjects with either 3- or 18-month onset (3mos, n=10; 18mos, n=10), as well as non-diabetic healthy controls (ND, n=10). Most participants carried at least one copy of the risk-conferring DR3-DQ2 or DR4-DQ8 haplotype^21,22^ (9 out of 10 in 3mos and 18mos; 6 out of 10 in ND) (Supplementary Table 1). Each participant fasted, had an initial blood draw, and then underwent a mixed meal tolerance test (MMTT) to stimulate β-cell degranulation (Fig. 1a). Under the MMTT condition, we collected blood samples at 90 and 120 minutes post-MMTT and monitored C-peptide levels at various time points throughout MMTT to evaluate β-cell function (Fig. 1a). To account for variability between individuals, we grouped PBMC samples by disease state (ND, 3mos, 18mos) and time point (0, 90, 120 min), combining individual samples within each cohort. We then performed sequential isolation of pan-HLA-DQ and pan-HLA-DR peptidomes, analyzing a total of 18 peptidome samples.

**Figure 1.**
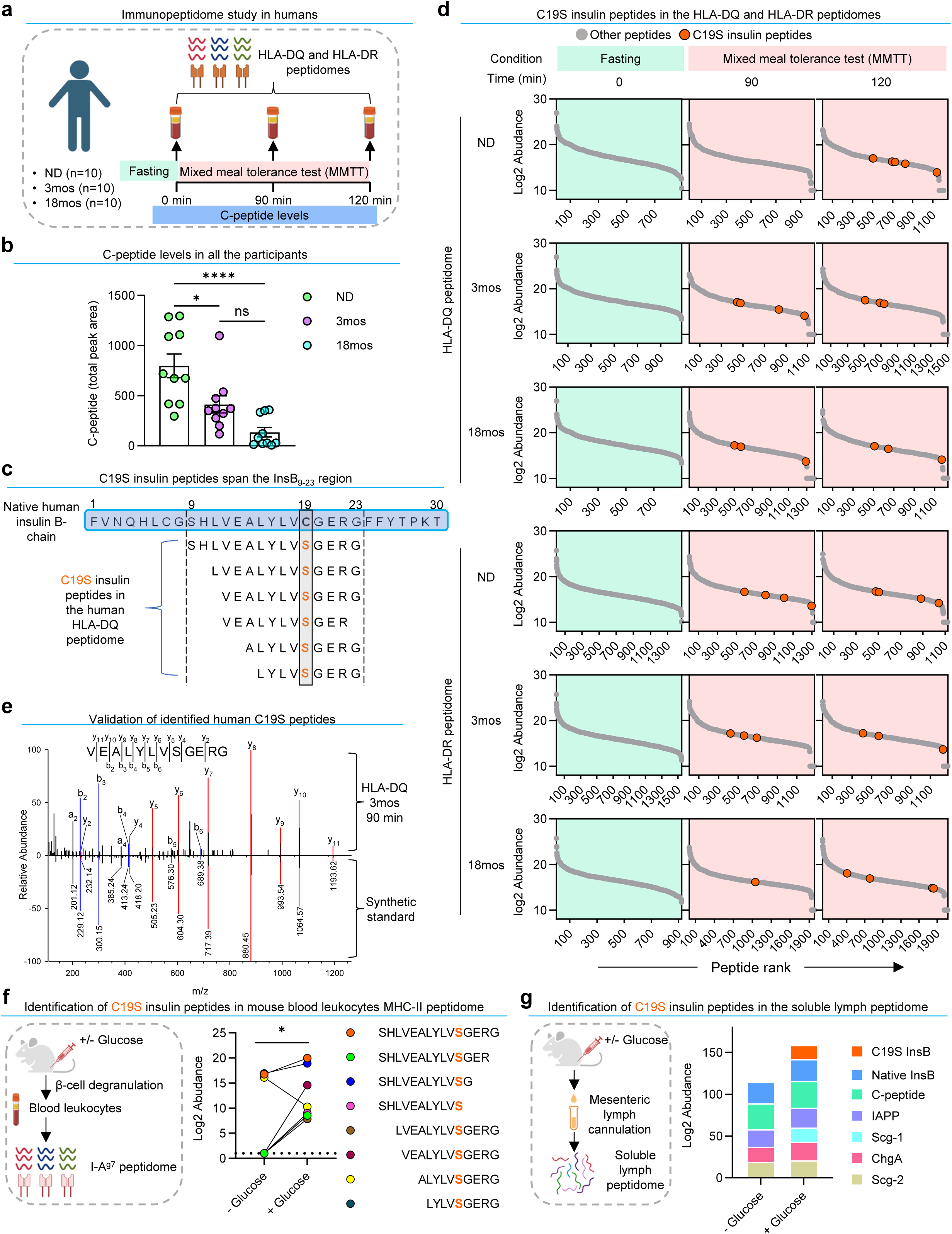
Identification of β-cell-derived insulin peptides with a Cys→Ser transformation in human and mouse MHC-II peptidomes. **a.** Schematic for analyzing human HLA-DQ and HLA-DR peptidomes of PBMCs from three cohorts of participants before and after a mixed meal tolerance test (MMTT). ND, non-diabetic; 3mos, 3-month onset; 18mos, 18-month onset. **b.** C-peptide levels of the three human cohorts. Each point represents the total peak area of an individual participant, calculated by summing measurements from all time points as shown in Extended Data Fig. 1b. **c.** Individual C19S insulin peptides identified in the HLA-DQ peptidome. **d.** Rank plots showing each C19S insulin peptide identified in the HLA-DQ (upper) and HLA-DR (lower) peptidomes across the three human cohorts at indicated time points of MMTT. Grey dots represent all other peptides in the peptidomes, and orange dots denote individual C19S insulin peptides. **e.** A mirror plot showing the complete match of a representative C19S insulin peptide identified in the human HLA-DQ peptidome to its synthetic standard. **f.** Schematic (left) and quantification (right) of insulin B-chain peptides with C19S (each point) identified in the blood leukocyte MHC-II peptidome of NOD mice, showing increased abundance after glucose challenge. *P < 0.05; Wilcoxon signed-rank test. **g.** Schematic (left) for analyzing soluble lymph peptidome and composition (right) of indicated β-cell-derived peptides identified in the soluble lymph peptidomes. The data are based on the cumulative abundance of all the peptides belonging to a given β-cell protein.

We previously detected native insulin peptides in the PBMC peptidome of prediabetic NOD mice given glucose stimulation^20^. Consistent with these results, we identified native insulin B-chain peptides in the HLA-DQ and HLA-DR peptidomes of the non-diabetic human samples (Extended Data Fig. 1a). Notably, all of these insulin peptides were only detected post-MMTT, not during fasting (Extended Data Fig. 1a), confirming that MMTT facilitated the detection of β-cell-derived peptides in the human PBMC HLA-II peptidome. Although the T1D cohorts showed diminished C-peptide levels (Fig. 1b; Extended Data Fig. 1b), indicating β-cell loss or dysfunction, several native insulin peptides remained detectable in their HLA-II peptidomes following MMTT. These peptides may derive from residual islets known to remain in T1D patients, with their identification also facilitated by MMTT.

Upon further analysis, we identified a previously unrecognized Cys→Ser transformation at position 19 of the insulin B-chain (e.g., SHLVEALYLV**<u>C</u>**GERG → SHLVEALYLV**<u>S</u>**GERG), hereafter referred to as C19S. To confirm these results, we repeated the database search with C19S insulin sequences appended to the canonical Uniprot-Human database, which yielded additional insulin B-chain peptides with C19S. All identified peptides exhibited characteristics typical of MHC-II-bound peptides: they ranged in length from 9-15 residues and spanned the B-chain 9-23 segment (InsB_9-23_) (Fig. 1c), a well-defined MHC-II-binding region in both NOD mice and humans with T1D^19,23–28^.

In both the HLA-DQ and HLA-DR peptidomes, C19S insulin peptides were only detected after MMTT (Fig. 1d), reinforcing the role of MMTT in enabling their detection and suggesting that they may be released along with native insulin. For further verification, we synthesized peptides based on 11 unique C19S insulin peptide sequences identified in the HLA-II peptidomes and confirmed that each peptide from the PBMC peptidome samples matched the synthetic standard controls (Fig. 1e; Extended Data Fig. 1c). Collectively, these data suggest that, in addition to native insulin peptides, the C19S variant can be presented by T1D-predisposing HLA-II molecules.

The identification of C19S insulin peptides in human HLA-II peptidomes prompted us to assess their presence in the mouse MHC-II peptidome. First, we reanalyzed the MHC-II (I-A^g7^) peptidomes from blood leukocytes of prediabetic NOD mice^20^ and identified a group of peptides with the same C19S transformation, which also increased in abundance following glucose challenge (Fig. 1f). Moreover, in our previous analysis of the I-A^g7^ peptidomes of pancreatic islets from NOD mice^26^, we identified and verified the exact InsB_9-23_(C19S) peptide (SHLVEALYLV**<u>S</u>**GERG) (Extended Data Fig. 1d).

In addition to MHC-II-bound peptides, we assessed soluble peptidomes of mesenteric lymph cannulated from 6-week-old female NOD mice with or without glucose challenge. At baseline (no glucose injection), the lymph peptidome already contained peptides from several secretory protein antigens in β cells, such as native insulin (B-chain and C-peptides), islet amyloid polypeptides (IAPP), and chromogranin A (ChgA), which further increased after glucose challenge (Fig. 1g). Notably, the InsB_9-23_(C19S) peptide was identified and verified in the lymph peptidome of glucose-injected mice (Extended Data Fig. 1e). Thus, C19S insulin peptides are identified in both soluble and MHC-II-bound forms and can be presented at disease-relevant sites in NOD mice.

### Disease-relevant signals amplify C19S transformation in pancreatic β cells

C19S insulin peptides were identified in the HLA-II peptidomes of both non-diabetic and diabetic individuals. This observation suggests that C19S transformation may occur in native insulin under non-autoimmune conditions, raising the question of whether this process is associated with disease progression. Addressing this question would require a direct comparison of C19S transformation between autoimmune and non-autoimmune settings. While the immunopeptidome assay enabled us to identify C19S insulin peptides, the statistical comparison between the T1D and non-diabetic cohort might be skewed, since the T1D patients have reduced overall levels of source insulin peptides due to islet loss.

To address this limitation, we developed quantitative antigen presentation assays utilizing CD4^+^ T cell hybridomas specifically recognizing the C19S insulin peptide or its native counterpart. From NOD mice immunized with the synthetic InsB_9-23_(C19S) peptide (SHLVEALYLV**<u>S</u>**GERG), we generated a panel of CD4^+^ T cell hybridomas, including the S5 clone, which showed robust reactivity to the InsB_9-23_(C19S) peptide, while remaining largely unresponsive to the native InsB_9-_

_23_ peptide (SHLVEALYLV**<u>C</u>**GERG) (Fig. 2a). This reactivity was distinct from the previously characterized 9B9 CD4^+^ T cell clone^19,20,29,30^, which specifically recognized native InsB_9-23_ but exhibited minimal reactivity to InsB_9-23_(C19S) (Fig. 2b). Notably, both T cells exhibited comparable reactivities to their respective peptides (Extended Data Fig. 2a), enabling quantitative comparisons.

**Figure 2.**
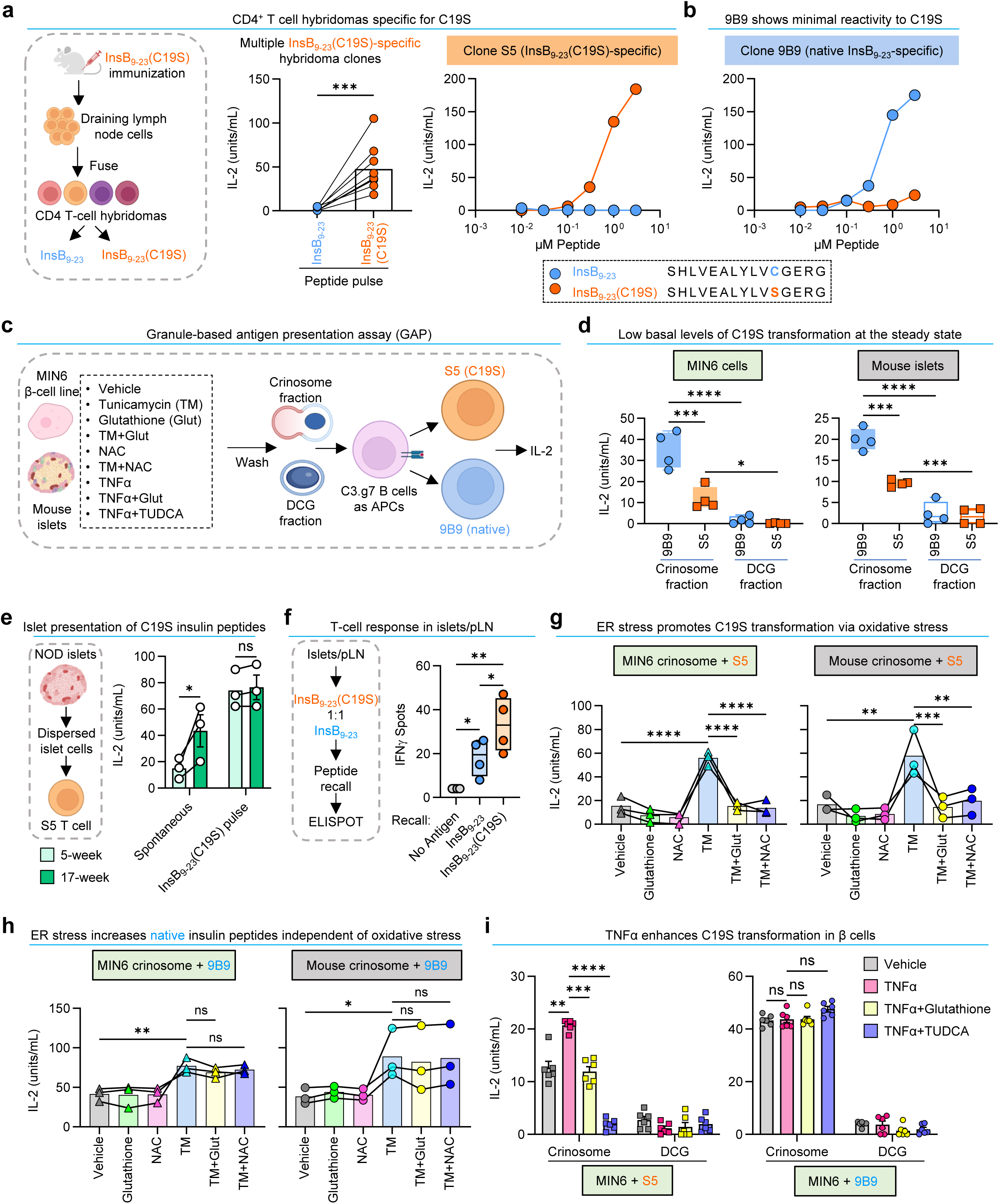
Disease-relevant signals drive C19S transformation via oxidative stress in β-cell granules. **a.** Reactivity of CD4^+^ T cell hybridomas specifically recognizing InsB_9-23_(C19S) but not native InsB_9-23_. Data show quantification (right) of multiple individual T cell hybridomas (each point) responding to 0.3 µM peptide and responses of the S5 T cell (left) to serially diluted peptide concentrations. **b.** Reactivity of the 9B9 T cell to the native InsB_9-23_ but not the InsB_9-23_(C19S) peptide. **c.** Schematic of the GAP assay. **d.** Responses of the 9B9 or the S5 T cell to crinosome or DCG fractions isolated from unmanipulated MIN6 cells or B6 mouse islets. **e.** In vivo presentation of C19S insulin peptides in pancreatic islets from 5- and 17-week-old female NOD mice. **f.** ELISPOT assay for T cell responses (IFNγ production) in islets and pLNs during antigen recall with the InsB_9-23_ or InsB_9-23_(C19S) peptide. **g.** Responses of the S5 T cell to the crinosome fraction isolated from MIN6 cells and B6 mouse islets following indicated treatments. **h.** Responses of the 9B9 T cell to the crinosome fraction isolated from MIN6 cells and B6 mouse islets following indicated treatments. **i.** Responses of the S5 or 9B9 T cell to crinosome and DCG fractions isolated from MIN6 cells stimulated with TNFα, with or without glutathione or TUDCA. Data (mean ± SEM) show results from multiple independent experiments (each point). All experiments using the GAP assay (**a, b, d, g, h, i**) used C3.g7 cells as APCs. ns, not significant; *P < 0.05; **P < 0.01; ***P < 0.001; ****P < 0.0001; two-tailed paired t test (**a**); repeated measures one-way ANOVA (**f, g, h**); repeated measures two-way ANOVA (**d, e, i**).

Utilizing these epitope-specific T cell hybridomas, we developed a granule-based antigen presentation (GAP) assay to assess changes in the generation of native and C19S insulin in β cells under steady-state and inflammatory conditions (Fig. 2c). We specifically assessed β-cell granules, which have been recognized as major sites for regular insulin biosynthesis and autoantigen generation in T1D^31,32^. Previous studies have found distinct insulin products in two sets of β-cell granules^19,29^: insulin dense-core granules (DCGs), which primarily store intact insulin, and crinosomes, a minor set of degradative vesicles generated through the fusion of excessive DCGs with lysosomes, primarily containing shorter peptide fragments. Based on these studies, we tested both crinosomes and DCGs in the GAP assay. Subcellular granule fractions representing DCGs and crinosomes were isolated from different sources of β cells, including a β-cell line and primary islets. Upon isolation, the crinosome and DCG fractions were validated by measuring their granule content and lysosomal activity (acid phosphatase levels) (Extended Data Fig. 2b-d). Furthermore, C3.g7 APCs (B cell lymphoma expressing I-A^g7^) were treated with these granules and examined for presentation of insulin peptides derived from the granules. The levels of C19S insulin peptide versus the native form were then probed using S5 and 9B9 T cells (Fig. 2c).

We first used the GAP assay to assess if C19S can occur in β cells in the absence of autoimmunity, given the identification of C19S insulin peptides in the MHC-II peptidomes from prediabetic NOD mice and non-diabetic humans. To this end, we tested granules isolated from the MIN6 β-cell line (derived from diabetes-resistant B6 mice) and primary islets from B6 mice. Both sources of β cells showed similar results. As reported previously^19,29^, the native InsB_9-23_ peptide (assayed by 9B9) was mostly found in crinosomes, with minimal presence in the DCG fraction (Fig. 2d). The crinosome fraction also activated the S5 T cell, indicating the presence of C19S peptides, but the responses were 30-50% lower than those observed with 9B9 (Fig. 2d). In contrast, S5 T cell responses to the DCG fraction were minimal (Fig. 2d). These data indicate that, under non-autoimmune conditions, C19S insulin peptides are primarily localized in crinosomes at low basal levels.

Next, we assessed whether C19S insulin peptide generation was associated with diabetes development. As an initial approach, we used the InsB_9-23_(C19S)-specific S5 T cell to examine the in vivo presentation in pancreatic islets of NOD mice. The S5 T cell responded to dispersed islet cells in the absence of exogenous antigen pulse (Fig. 2e), indicating natural presentation of C19S insulin peptides in islets. Although such presentation was detectable in islets from 5-week-old mice, we observed a significant increase in islets from 17-week-old prediabetic NOD mice (Fig. 2e). These results led us to functionally test responses to C19S insulin peptides in diabetes-relevant locations. We stimulated cells pooled from islets and pancreatic lymph nodes (pLNs) with equal amounts of InsB_9-23_ and InsB_9-23_(C19S) peptides and assessed T cell responses (IFNγ production) via ELISPOT during antigen recall. As reported previously^26,29^, recall with native InsB_9-23_ induced IFNγ production (Fig. 2f). However, recall with the InsB_9-23_(C19S) peptide triggered even higher IFNγ responses (Fig. 2f). Thus, although C19S insulin peptides are produced at low levels at steady state, their presentation increases with disease progression, suggesting that C19S transformation is not a static event but can be amplified during diabetes development.

To further investigate the mechanisms underlying C19S transformation in autoimmune conditions, we assessed the roles of disease-relevant signals, including ER stress, oxidative stress, and inflammatory cytokines, in this process. The cysteine 19 residue of the insulin B-chain (B(C19)) is the first residue to form a disulfide bond with cysteine 20 of the A-chain, a step critical for proper insulin folding in the ER^33,34^. We reasoned that C19S might reflect insulin misfolding, a process closely linked to ER stress^35,36^. Additionally, ER stress can induce the production of reactive oxygen species (ROS), resulting in oxidative stress in β cells^37^. These findings, along with evidence that cysteine can be converted to serine under oxidative conditions^38^, prompted us to test the roles of ER and oxidative stress in C19S transformation in β cells.

To assess the role of ER stress using the GAP assay, we exposed MIN6 cells and B6 mouse islets to the ER stress inducer tunicamycin for 2 hours, followed by extensive washing. This short treatment did not cause β-cell death or alter total protein levels in the granule samples (Extended Data Fig. 2e). However, the S5 T cell exhibited markedly enhanced responses to crinosome fractions isolated from MIN6 cells and B6 mouse islets treated with tunicamycin (Fig. 2g), suggesting that ER stress in β cells can directly promote C19S transformation.

To assess the role of oxidative stress, we included the antioxidants glutathione or N-acetylcysteine (NAC) in the GAP assay. Notably, the tunicamycin-induced increase in C19S transformation was significantly inhibited by glutathione or NAC in both MIN6 cells and mouse islets (Fig. 2g). In contrast, although the tunicamycin treatment also elevated native InsB_9-23_ levels in crinosomes, this effect was not influenced by NAC or glutathione (Fig. 2h). Additionally, at baseline, DCGs from MIN6 cells and B6 mouse islets contained minimal native or C19S insulin peptides, while tunicamycin increased both forms (Extended Data Fig. 2f,g). Glutathione and NAC also selectively inhibited the increase of C19S (Extended Data Fig. 2f) but not native insulin peptides (Extended Data Fig. 2g) in DCGs. These results collectively demonstrate that C19S transformation in insulin is specifically driven by oxidative stress in the islet microenvironment, a feature that distinguishes them from native insulin peptides.

Next, we assessed the role of inflammatory cytokines in C19S transformation. T1D development is associated with inflammatory cytokine signaling in pancreatic islets, with IL-1β, IFNγ, and TNFα known to induce ER and oxidative stress in β cells^39,40^. Since inflamed islets from NOD mice may already have active cytokine signaling, which potentially interferes with the assessment of the effects of individual cytokines, we tested the influence of IL-1β, IFNγ, and TNFα on C19S transformation in MIN6 cells. Initially, we conducted a pilot screen using a glucose-induced antigen transfer assay, which suggested that TNFα or IFNγ exposure increased production of C19S insulin peptides, with TNFα showing the strongest effect (Extended Data Fig. 2h). Confirming these results, the GAP assay also revealed a significant increase in S5 T cell responses to crinosome fractions from TNFα-treated MIN6 cells (Fig. 2i). This increase was inhibited by glutathione or by tauroursodeoxycholic acid (TUDCA) (Fig. 2i), an ER stress inhibitor known to suppress T1D development^41^. In contrast, TNFα did not affect native InsB_9-23_ levels and had minimal impact on DCGs (Fig. 2i). These results demonstrate that inflammatory cytokines, particularly TNFα, promote C19S transformation by exacerbating ER and redox stress. This interplay between oxidative stress, ER stress, and cytokine signaling may contribute to the observed increase in C19S epitope presentation as disease progresses.

### C19S represents a context-dependent single amino acid transformation

The generation of C19S insulin peptides in stressed and inflamed islets suggests that C19S transformation is microenvironment-driven and context-dependent, potentially representing a rare event in the proteome. Since antigen presentation is a highly selective process, the identification of C19S in MHC-II-bound insulin peptides in both mice and humans is somewhat unexpected. We reasoned that the high abundance of insulin in β-cell granules, combined with oxidative sensitivity of the B(C19) residue, might particularly facilitate the generation of C19S insulin peptides, enabling their presentation. Since targeted analyses, such as the GAP assay, examine only predefined peptides, we decided to use a systematic approach to examine whether C19S could be unbiasedly identified among the broad spectrum of PTMs and single amino acid variants (SAVs) and how β-cell ER stress may shape these processes.

Considering that C19S was mostly detected in degraded post-synthetic insulin peptides in crinosomes at steady state and that the proteome profile of MIN6 cells closely resembles primary β cells^42^, we isolated crinosome peptidomes from equal numbers of MIN6 cells treated with vehicle or tunicamycin for 2 hours (Fig. 3a). The samples were prepared in triplicate, fractionated to increase depth of coverage, and block randomized for analysis by MS to minimize confounding effects during the analytical process. After characterizing the protein composition through the initial PEAKS DB search, we performed sequential PEAKS PTM and SPIDER searches to identify initially unspecified PTM and all possible SAVs, respectively. For quantification, the identified PTMs and SAVs were collated at the peptide-spectrum match (PSM) level to perform an unbiased survey of all changes between the two conditions.

**Figure 3.**
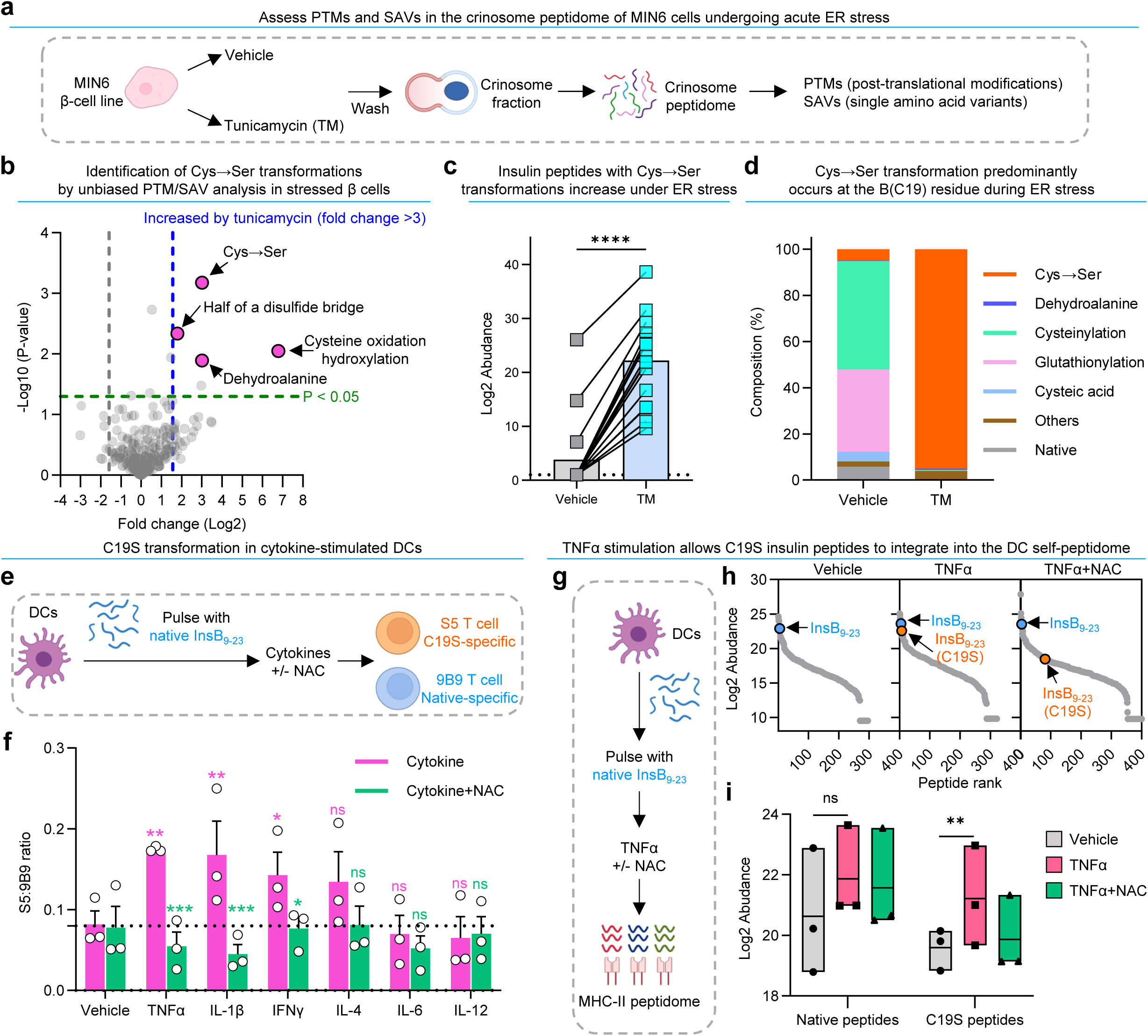
C19S represents a context-dependent single amino acid transformation emerging in β cells and dendritic cells. **a.** Schematic of the PTM/SAV search in crinosome peptidomes isolated from MIN6 β cells treated with vehicle or tunicamycin for 2 hours. **b.** A volcano plot showing changes in PTMs and SAVs identified in the crinosome peptidome of MIN6 cells given tunicamycin relative to vehicle treatment. Several cysteine modifications with significant increase (P < 0.05, fold change > 3) after treatment with tunicamycin were indicated. See a complete list in Supplementary Table 2. **c.** Abundance of individual insulin peptides with Cys→Ser transformations (each point) identified in the crinosome peptidome of MIN6 cells given vehicle or tunicamycin treatment. See a complete list in Supplementary Table 3. ****P < 0.0001; Wilcoxon signed-rank test **d.** Composition of indicated PTMs and Cys→Ser transformation on the B(C19) residue of insulin peptides identified in the crinosome peptidome of MIN6 cells given vehicle or tunicamycin treatment. **e.** Schematic of the antigen presentation assay to assess the presentation of native or C19S insulin peptides by DCs pulsed with native InsB_9-23_ and stimulated with different cytokines. **f.** C19S transformation from the native InsB_9-23_ peptide in DCs upon cytokine stimulation, shown as the ratio of S5 to 9B9 T cell responses with or without NAC treatment. Data (mean ± SEM) summarize results from three independent experiments (each point). ns, not significant; *P < 0.05; **P < 0.01; ***P < 0.001; repeated measures two-way ANOVA. Pink stars: cytokine vs. vehicle; green stars: cytokine vs. cytokine ^+^ NAC. **g.** Schematic (left) of MHC-II immunopeptidome analysis of InsB_9-23_-pulsed DCs stimulated with TNFα with or without NAC. **h.** Rank plots showing all identified peptides in the DC MHC-II peptidomes (grey dots), ranked by abundance. Native InsB_9-23_ and modified InsB_9-23_(C19S) peptides are denoted. **i.** Abundance of native and C19S insulin peptides identified in the DC MHC-II immunopeptidome analyses. Each point represents cumulative abundance of all identified native or C19S insulin peptides from one of the three independent experiments. ns, not significant; **P < 0.01; repeated measures two-way ANOVA.

Among the 348 identified PTMs and SAVs (Table S2), Cys→Ser transformations exhibited the most significant increase following tunicamycin treatment (Fig. 3b). Notably, Cys→Ser was identified as a SAV in the SPIDER search rather than through the PTM search. However, the ER stress condition also increased several PTMs, especially cysteine modifications, such as oxidation and hydroxylation, though to a lesser extent than Cys→Ser (Fig. 3b). We did not observe significant changes in oxidative modifications on other residues (e.g., histidine, tryptophan, proline, methionine) (Supplementary Table 2), suggesting that the observed cysteine oxidations reflect biological changes rather than sample preparation artifacts. Collectively, these results confirm the known role of ER stress in promoting the generation of conventional PTMs and reveal that the Cys→Ser transformation emerges rapidly and prominently in β cells responding to acute ER stress.

Tracing Cys→Ser transformations back to their source peptides revealed that most were localized to the insulin B chain. To verify these results, we first created a database in which all cysteine residues were replaced by serine and found no additional peptides with Cys→Ser transformations. Next, we appended Cys→Ser variants of abundant β-cell proteins (insulin-1, insulin-2, IAPP, ChgA, and Scg-1) to the canonical database and repeated the search. This confirmed the identification of Cys→Ser transformations in various insulin B-chain peptides (total 16) (Supplementary Table 3), most of which occurred on the B(C19) residue (Supplementary Table 3), consistent with results seen in the MHC-II peptidome analyses. Some peptides had dual transformations at both the 7^th^ and 19^th^ positions (e.g., **<u>C</u>**GSHLVEALYLV**<u>C</u>**GERG→**<u>S</u>**GSHLVEALYLV**<u>S</u>**GERG). All of these insulin peptides exhibited a significant increase in their abundance following tunicamycin treatment (Fig. 3c; Supplementary Table 3).

We then specifically analyzed the B(C19) residue and assessed how the Cys→Ser transformation compared to other PTMs. In the vehicle-treated control, B(C19) exhibited PTMs like cysteinylation, glutathionylation, and formation of cysteic acid and dehydroalanine, whereas Cys→Ser was only a minor component (Fig. 3d). However, after the 2-hour tunicamycin treatment, the Cys→Ser transformation emerged as the dominant event (Fig. 3d), supporting the evidence that this process is particularly driven by the stressed islet microenvironment compared to conventional PTMs on the same residue. Overall, these results demonstrate that Cys→Ser is context dependent and preferentially occurs at the redox-sensitive B(C19) residue of insulin in β cells.

### C19S transformation in cytokine-activated DCs

Our analysis suggests that MHC-II presentation of C19S insulin peptides is amplified as diabetes develops. Although β cells can increase C19S transformation under stress and inflammation, effective MHC-II presentation may also depend on an amplification process of presentation within APCs. During T1D development, dendritic cells (DCs) migrate to inflamed islets, where they actively present β-cell antigens, including native insulin peptides^29,43^, while simultaneously being exposed to inflammatory signals such as cytokines. Although cytokine stimulation enhances licensing and antigen presentation capacity of DCs, whether these signals also contribute to neoantigen formation remains less clear. These findings prompted us to test whether C19S transformation could also occur within cytokine-activated DCs.

As an initial approach, we enriched DCs in vivo by treating NOD mice with recombinant FLT3 ligand, isolated DCs, and subsequently pulsed them with the native InsB_9-23_ peptide. Equal aliquots of these InsB_9-23_-pulsed DCs were then exposed to different cytokines, with or without the antioxidant NAC. The presentation levels of the native InsB_9-23_ and InsB_9-23_(C19S) peptide were probed by 9B9 and S5 T cells (Fig. 3e). We measured the ratio of S5 to 9B9 T cell responses (S5:9B9) to evaluate the C19S transformation. Stimulation with IL-1β, TNFα, and IFNγ significantly increased the S5:9B9 ratio, suggesting C19S transformation from the native peptide, while the influences of IL-4, IL-6, and IL-12 were less profound (Fig. 3f). Furthermore, NAC largely blocked the increase in C19S peptide levels induced by IL-1β, TNFα, and IFNγ (Fig. 3f), indicating the dependence on oxidative stress.

The antigen presentation assay suggested that cytokine stimulation allowed C19S insulin peptides to integrate into the broad and competitive DC self-peptidome. To investigate this, we pulsed DCs with InsB_9-23_, treated them with TNFα or TNFα plus NAC, and isolated I-A^g7^ peptidomes for analysis by MS (Fig. 3g). While the native InsB_9-23_ peptide was identified in the DC peptidome under all conditions, the InsB_9-23_(C19S) peptide appeared in TNFα-stimulated DCs (Fig. 3h). We observed a significant increase in all identified C19S insulin peptides in three independent experiments following TNFα stimulation, which was inhibited by NAC (Fig. 3i), demonstrating that C19S transformation can occur de novo in DCs under cytokine stimulation.

### C19S alters CD4^+^ T cell recognition in a register-dependent manner

The generation of CD4^+^ T cell hybridomas responding to the InsB_9-23_(C19S) peptide but not native InsB_9-23_ (Fig. 2a) suggests the existence of CD4^+^ T cells specifically recognizing C19S in NOD mice. This observation was unexpected, given that the binding properties of the InsB_9-23_ segment to I-A^g7^ and its corresponding CD4^+^ T cell specificities have been extensively characterized. To address this, we examined how C19S might alter I-A^g7^ binding and whether it could be recognized by register-specific CD4^+^ T cells in relation to classic insulin-reactive T cells.

C19S is positioned within two 9-mer I-A^g7^-binding cores, InsB_12-20_ (VEALYLV**<u>C</u>**G) and InsB_13-21_ (EALYLV**<u>C</u>**GE), which use G20 and E21, respectively, as their P9 anchor residues^44^. This register shift allows native InsB_12-20_ and InsB_13-21_ to be recognized by two distinct CD4^+^ T cell populations through a mechanism called P9 switch^27^. To evaluate how C19S affects register recognition, we synthesized peptides with InsB_12-20_(C19S) (VEALYLV**<u>S</u>**G) or InsB_13-21_(C19S) (EALYLV**<u>S</u>**GE) as putative registers, each nested within surrogate flanking residues^44^ (Extended Data Fig. 3a). In the InsB_13-21_ register, C19S resides in P7, an I-A^g7^-binding pocket. For the InsB_12-20_ register, however, C19S introduces a new TCR contact residue at P8. As a promiscuous peptide binder, I-A^g7^ has an unusual binding groove with a widened entrance to the P9 pocket, suggesting that changes in P8 could also impact binding^45,46^. Based on these findings, we performed both cell-based (Extended Data Fig. 3b) and biochemical (Extended Data Fig. 3c) binding assays to assess how C19S influences peptide binding.

All peptides with C19S exhibited a marked reduction in binding compared to their native counterparts (Extended Data Fig. 3b,c). Native InsB_12-20_ is already a weak-binding epitope, forming unstable peptide-MHC-II complexes^26,27,29,44^; yet the nested InsB_12-20_(C19S) peptide showed an additional reduction in binding, making it the weakest-binding peptide among those tested (Extended Data Fig. 3b,c). To verify this effect, we tested a surrogate InsB_12-20_(C19A) peptide (Extended Data Fig. 3a), which also showed reduced binding to I-A^g7^ (Extended Data Fig. 3d). Thus, consistent with recent evidence that P8 residues can modulate MHC-II binding^47,48^, the C19S transformation reduced the binding of the InsB_12-20_ epitope to I-A^g7^.

Despite this, most InsB_9-23_(C19S)-specific T cell hybridomas (Fig. 2a) preferentially recognized the InsB_12-20_(C19S) register, with only a few reacting to InsB_13-21_(C19S) (Extended Data Fig. 3e). Furthermore, substituting G20 or E21 with the inhibitory residue K abolished all T cell responses (Extended Data Fig. 3e), confirming that G20 and E21 remain as P9 anchor residues in C19S-containing registers.

Since C19S introduces a change at the P8 TCR contact residue in the InsB_12-20_ register, we reasoned that this may explain the observed T cell specificity to the InsB_12-20_(C19S) register. To test this in the endogenous T cell pool, we produced I-A^g7^-based tetramers incorporating the InsB_12-20_(C19S) and InsB_13-21_(C19S) epitopes. In our previous studies^27–29^, we generated tetramers containing the native InsB_12-20_ and InsB_13-21_ epitopes, which allowed for tracking the cognate register-specific CD4^+^ T cells in vivo. All tetramers used the exact 9-mer epitopes (native or C19S) to ensure specificity (Extended Data Fig. 3f). Using the magnetic enrichment protocol^49^ (Extended Data Fig. 3g), neither tetramer stained CD8^+^ T cells in NOD mice nor CD4^+^ T cells from a NOD strain expressing I-A^b^ instead of I-A^g7^ (Extended Data Fig. 3h), confirming MHC restriction.

The InsB_12-20_(C19S):I-A^g7^ tetramer identified a distinct CD4^+^ T cell population in secondary lymphoid organs (SLOs; pooled spleens and various lymph nodes) of 8-week-old female NOD mice, with minimal overlap with CD4^+^ T cells labeled by the native InsB_12-20_:I-A^g7^ tetramer (Fig. 4a). Moreover, the frequencies of these InsB_12-20_(C19S)-specific CD4^+^ T cells were consistently higher than those specific for the native InsB_12-20_ epitope (Fig. 4a). As an additional control, an InsB_12-20_(C19A):I-A^g7^ tetramer likewise identified a discrete T cell population distinct from native InsB_12-20_-specific cells (Extended Data Fig. 3i). In contrast, co-staining with the InsB_13-21_:I-A^g7^ and InsB_13-21_(C19S):I-A^g7^ tetramers revealed a largely overlapping population, indicating cross-reactivity (Extended Data Fig. 3j). These results highlight that T cell recognition of C19S depends on its positioning, as substitution at P7 in InsB_13-21_ does not significantly alter recognition, whereas its placement at the P8 TCR contact position in the InsB_12-20_ register corresponds to a distinct CD4^+^ T cell population.

**Figure 4.**
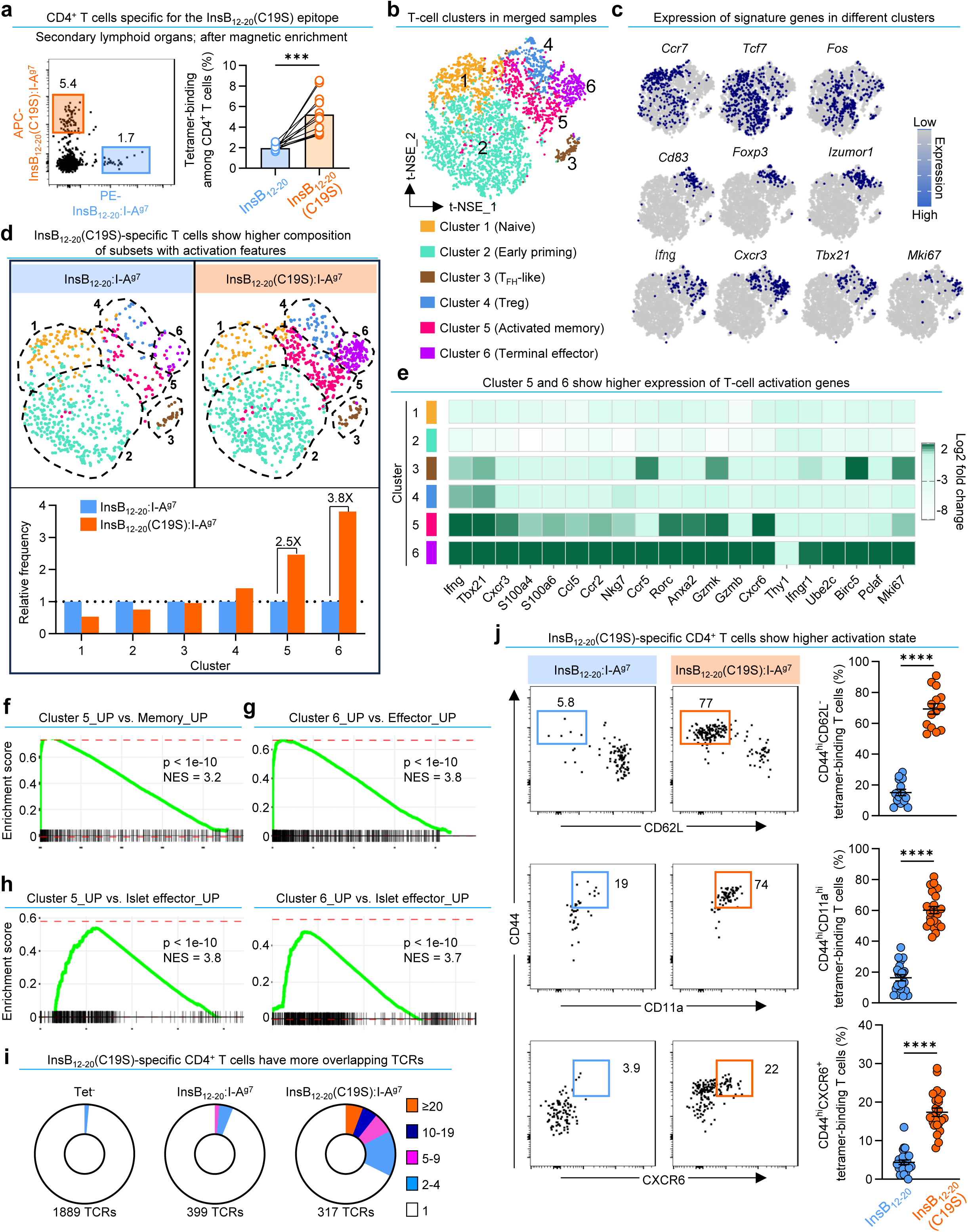
C19S-specific CD4⁺ T cells exhibit distinct transcriptional states marked by peripheral activation and clonal expansion. **a.** Flow cytometry analysis (left) showing co-staining with InsB_12-20_:I-A^g7^ and InsB_12-20_(C19S):I-A^g7^ tetramers revealing two distinct register-specific T cell populations. The quantification (right) summarizes the frequencies of the tetramer-binding populations in individual mice (each point) from six independent experiments. ***P < 0.001; two-tailed paired t test. **b.** t-distributed stochastic neighbor embedding (t-SNE) plots depicting T cell clusters merged from InsB_12-20_-specific, InsB_12-20_(C19S)-specific, and Tet^⁻^ CD4^+^ T cells. **c.** Feature plots showing the expression of functional genes classifying CD4^+^ T cell clusters. **d.** t-SNE plots (upper) showing cluster distribution between InsB_12-20_- and InsB_12-20_(C19S)-specific CD4^+^ T cells. The bar graph (lower) depicts the relative frequency of each cluster. **e.** A heat map showing the expression of indicated T cell activation and cycling genes among all clusters. **f.** GSEA showing enrichment of cluster 5 for memory T cell signatures from dataset GSE9650. **g.** GSEA showing enrichment of cluster 6 for effector T cell signatures from dataset GSE9650. **h.** GSEA showing enrichment of both cluster 5 and 6 for intra-islet, terminally activated CD4^+^ T cells depicted in dataset GSE262101. **i.** Pie charts showing clonal overlap of TCRs identified in tetramer-negative, InsB_12-20_-specific, and InsB_12-20_(C19S)-specific CD4^+^ T cells. Clonal overlap was defined as TCRs with identical amino acid sequences in the full CDR3 regions of both α and β chains. **j.** Flow cytometry analysis of indicated T cell activation markers in InsB_12-20_- and InsB_12-20_(C19S)-specific CD4^+^ T cells in SLOs of 8-week-old female NOD mice. The quantification (mean ± SEM) summarizes cell frequencies in individual mice (each point) from 7-10 independent experiments. ****P < 0.0001; one-way ANOVA with Dunnett’s multiple comparisons test.

### Distinct transcriptional heterogeneity of C19S-specific CD4^+^ T cells

Previous studies have characterized transcriptional and functional signatures of CD4⁺ T cells recognizing native insulin epitopes in NOD mice^19,26,27,29,50,51^. The fact that the InsB_12-20_(C19S) epitope only differs from its native counterpart by a single residue prompted us to determine whether these C19S-specific CD4⁺ T cells exhibit distinctive transcriptional features. To this end, we performed single-cell RNA sequencing (scRNA-seq) on InsB_12-20_(C19S):I-A^g7^ and InsB_12-20_:I-A^g7^ tetramer-binding CD4⁺ T cells, with tetramer-negative (Tet⁻) CD4⁺ T cells as controls. We obtained and analyzed 719 InsB_12-20_:I-A^g7^ and 1,256 InsB_12-20_(C19S) tetramer-binding cells sorted from SLOs of 8-week-old female NOD mice and performed scRNA-seq in replicates. Unsupervised clustering of the combined datasets revealed six clusters (Fig. 4b) characterized by differential expression of phenotypic T cell genes (Fig. 4c).

InsB_12-20_(C19S):I-A^g7^ tetramer-binding cells shared several transcriptional signatures previously reported in InsB_12-20_-reactive T cells. We previously found that antigen encounter by InsB_12-20_-reactive CD4^+^ T cells is not restricted to the pancreatic lymph node but disseminates across various SLOs^19^. During this weak but successive antigen exposure, these T cells acquired an early priming state characterized by upregulation of several TCR and NFκB activation genes before the expression of typical effector genes^19,29^. This signature was reflected in cluster 2, which was significantly enriched for the gene profile of the InsB_12-20_-specific 8F10 TCR transgenic CD4^+^ T cells upon antigen encounter in the distal inguinal lymph node^19^ (Extended Data Fig. 4a), as determined by gene set enrichment analysis (GSEA). Confirming this early priming signature, most InsB_12-20_:I-A^g7^ tetramer-binding cells were found in cluster 2. About half of InsB_12-20_(C19S):I-A^g7^ tetramer-binding cells were also classified as part of cluster 2 (Extended Data Fig. 4b), suggesting presentation and recognition of the InsB_12-20_(C19S) epitope in the periphery.

InsB_12-20_-reactive CD4^+^ T cells were shown to differentiate into T follicular helper cells (T_FH_) in SLOs of NOD mice, assisting production of class-switched insulin autoantibodies^51^. Consistent with this, we identified cluster 3, which exhibited T_FH_-like signatures, significantly enriched for a dataset describing transcriptional profiles of T_FH_ cells^52^ (Extended Data Fig. 4c). Cluster 3 was detected in both InsB_12-20_(C19S):I-A^g7^ and InsB_12-20_:I-A^g7^ tetramer-binding CD4^+^ T cells, with small and comparable proportions (Extended Data Fig. 4b).

Despite these similarities, InsB_12-20_(C19S):I-A^g7^ tetramer-binding cells contained fewer naïve-like T cells (cluster 1) than their native counterparts, suggesting a more antigen-experienced profile. This distinction aligns with their increased representation in clusters 5 and 6 (Fig. 4d; Extended Data Fig. 4b), which are associated with T cell activation, memory formation, and terminal differentiation. Both cluster 5 and 6 significantly upregulated typical T cell activation genes, such as *Ifng*, *Cxcr6*, *Tbx21*, compared to other clusters (Fig. 4e; Extended Data Fig. 4d). By GSEA, cluster 5 was also enriched for memory T cell signatures^53^ (Fig. 4f), consistent with previous observations of early memory responses in diabetes development in NOD mice^54^. Cluster 5 may therefore represent activated memory T cells. Cluster 6 had the strongest expression of effector T cell genes with further upregulation of cell cycling genes such as *Ube2c*, *Birc5*, *Pclaf*, and *Mki67* (Fig. 4e), and was significantly enriched for defined effector T cell signatures^53^ (Fig. 4g), indicating terminal differentiation. Furthermore, both cluster 5 and cluster 6 were enriched for the gene profile of a previously identified intra-islet effector CD4^+^ T cell subset with terminal activation^55^ (Fig. 4h).

The different transcriptional heterogeneity of InsB_12-20_(C19S):I-A^g7^ tetramer-binding cells suggests that they can progress into a heightened activation state in the periphery. Supporting this, single-cell TCR sequencing revealed a higher degree of clonal overlap (TCRs with identical CDR3α and CDR3β amino acid sequences) in InsB_12-20_(C19S):I-A^g7^ tetramer-binding cells (Fig. 4i; Extended Data Fig. 4e), indicating clonal expansion. To further validate the transcriptional data, we used flow cytometry to assess several activation markers highly expressed by cluster 5 and 6, including CD44, CD11a, and CXCR6, in SLOs of individual 8-week-old female NOD mice. InsB_12-20_(C19S):I-A^g7^ tetramer-binding cells consistently showed significantly higher proportions of CD44^hi^CD62L^lo^, CD44^hi^CD11a^hi^, and CD44^hi^CXCR6^+^ subsets, as compared to InsB_12-20_:I-A^g7^ tetramer-binding cells (Fig. 4j), demonstrating a more activated profile of InsB_12-20_(C19S)-specific CD4^+^ T cells at the protein level.

### Epitope availability and inflammatory cues regulate C19S-specific CD4⁺ T cell activation

InsB_12-20_(C19S)-specific CD4⁺ T cells exhibit a higher activation phenotype compared to their InsB_12-20_-specific counterparts. This raised the question of whether the observed activation profile truly depends on the presence of the C19S neoepitope in vivo, prompting us to search for a model that would genetically ablate reactivity to C19S insulin. Because the B(C19) residue is essential for proper insulin folding, directly manipulating this residue would likely disturb insulin biosynthesis and impair β-cell function^56^. While evaluating alternative models, we unexpectedly found that the mutant transgenic insulin expressed by the NOD.*B16A* mouse, which carries a tyrosine-to-alanine substitution at position 16 of the B-chain (Y16A)^23^, did not support C19S-specific T cell recognition. This allowed us to examine C19S-specific T cell activation in the absence of cognate epitope recognition.

Confirming previous findings showing that the Y16A mutation abolishes the immunogenicity of the native InsB_9-23_ peptide^19,23^, the 9B9 T cell (native InsB_9-23_-specific) showed no reactivity to the synthetic InsB_9-23_(Y16A) peptide (SHLVEAL**<u>A</u>**LVCGERG) (Fig. 5a). However, the InsB_9-23_(C19S)-specific S5 T cell also failed to respond to any concentration of the InsB_9-23_(Y16A) peptide during in vitro stimulation (Fig. 5a), indicating that the NOD.*B16A* mice lack the necessary epitope for C19S-specific T cell recognition.

**Figure 5.**
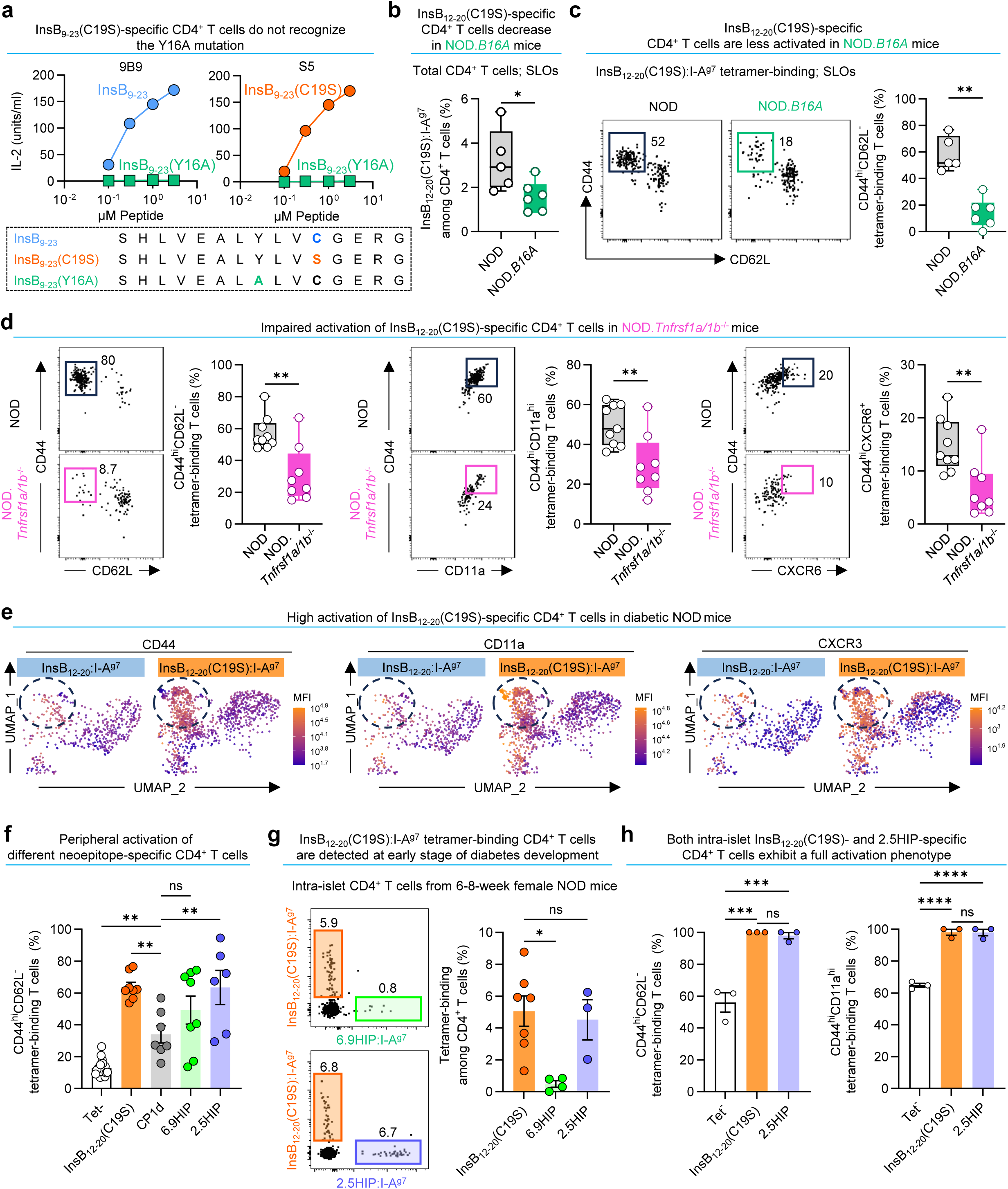
Epitope availability and inflammatory cues regulate peripheral activation, tissue infiltration, and diabetogenic potential of C19S-specific CD4⁺ T cells. **a.** IL-2 production by 9B9 and S5 T cells in response to InsB_9-23_, InsB_9-23_(C19S), and InsB_9-23_(Y16A) peptides. **b.** Frequencies of tetramer-binding InsB_12-20_(C19S)-specific CD4^+^ T cells in SLOs of 8-week-old female NOD and NOD.*B16A* mice. Data summarize results of individual mice (each point) examined in three independent experiments. *P < 0.05; Mann-Whitney test. **c.** Flow cytometry analysis (left) and quantification (right) showing the percentage of CD44^hi^CD62L⁻ activated T cells within tetramer-binding InsB_12-20_(C19S)-specific CD4^+^ T cells in SLOs of 8-week-old female NOD and NOD.*B16A* mice. Data summarize results of individual mice (each point) examined in three independent experiments. **P < 0.01; Mann-Whitney test. **d.** Flow cytometry analysis and quantification showing the expression of indicated T cell activation markers in tetramer-binding InsB_12-20_(C19S)-specific CD4^+^ T cells in SLOs of 8-week-old female WT NOD and NOD.*Tnfrsf1a/1b*^⁻/⁻^ mice. Data summarize results from individual mice (each point) examined in four independent experiments. **P < 0.01; Mann-Whitney test. **e.** Feature plots illustrating the expression of CD44, CD11a, and CXCR3 on InsB_12-20_:I-A^g7^ vs. InsB_12-20_(C19S):I-A^g7^ tetramer-binding CD4^+^ T cells from SLOs of diabetic NOD mice. Shown are equal numbers of tetramer-binding T cells for either epitope merged from eight mice. **f.** Percent of CD44^hi^CD62L^⁻^ activated T cells within indicated tetramer-binding CD4^+^ T cells in SLOs from 8-week-old female NOD mice. **g.** Flow cytometry analysis showing co-staining of intra-islet CD4^+^ T cells with the InsB_12-20_(C19S):I-A^g7^ tetramer along with the 6.9HIP:I-A^g7^ or 2.5HIP:I-A^g7^ tetramer. The quantification (right) summarizes the percentage of indicated tetramer-binding cells among intra-islet CD4^+^ T cells. **h.** Percentage of activated CD44^hi^CD62L⁻ and CD44^hi^CD11a^hi^ T cells among indicated intra-islet CD4^+^ T cell populations. Data (mean ± SEM) summarize results from at least three independent experiments. Each point represents individual mice (**b, d, f, g, h**). ns, not significant; *P < 0.05; **P < 0.01; ***P < 0.001; ****P < 0.0001; Mann-Whitney test (**b, c, d**); one-way ANOVA (**f, g, h**).

We then assessed InsB_12-20_(C19S):I-A^g7^ tetramer-binding CD4^+^ T cells in age/sex-matched NOD.*B16A* mice and wild-type (WT) NOD controls. Compared to WT mice, NOD.*B16A* mice exhibited a significant reduction in both InsB_12-20_(C19S)-specific CD4^+^ T cell frequency (Fig. 5b) and activation state (Fig. 5c), demonstrating that the presence of the C19S neoepitope is required for the maintenance and activation of InsB_12-20_(C19S)-specific CD4⁺ T cells in vivo.

Given the prominent role of TNFα signaling in promoting C19S transformation in both β cells and DCs, as well as in providing co-stimulatory signals to T cells, we assessed NOD mice deficient in both TNF receptor genes (NOD.*Tnfrsf1a/1b*^⁻/⁻^). These mice showed a significant reduction in the proportion of CD44^hi^CD62L^lo^, CD44^hi^CD11a^hi^, and CD44^hi^CXCR6^+^ InsB_12-20_(C19S):I-A^g7^ tetramer-binding CD4^+^ T cells, compared to age/sex-matched controls (Fig. 5d), indicating that TNFα signaling is important for optimal activation.

Because TNFα can influence both C19S insulin peptide generation and inflammatory co-stimulation, we next sought to distinguish these roles. To this end, we examined NOD mice with overt diabetes, when C19S generation is likely reduced due to β-cell loss, while inflammatory cues are prominent. T cell activation was evaluated by the expression of CD44, CD11a, and CXCR3. We observed an evident population of C19S-specific CD4⁺ T cells with high expression of all three activation markers, whereas native epitope-specific T cells showed substantially lower expression levels (Fig. 5e). These results indicate that the inflammatory milieu present at the diabetic stage can sufficiently sustain a high activation state in C19S-specific CD4⁺ T cells, even in the context of minimal antigen availability.

### C19S-specific CD4⁺ T cells migrate to pancreatic islets and acquire terminal activation

Consistent with previous studies^23,57^, both NOD.*B16A* and NOD.*Tnfrsf1a/1b*^⁻/⁻^ mice in our colony were resistant to diabetes development (Extended Data Fig. 5a). The observed inactivation of InsB_12-20_(C19S)-specific CD4^+^ T cells in both strains suggested their relevance to disease pathogenesis. To test this, we enriched polyclonal InsB_12-20_(C19S)-specific CD4^+^ T cells from InsB_9-23_(C19S)-immunized NOD mice, expanded them in vitro, and adoptively transferred them to NOD.*Rag1*^⁻/⁻^ recipients. Given that some InsB_9-23_(C19S)-reactive T cells responded to the InsB_13-21_(C19S) epitope (Extended Data Fig. 3d), we selectively expanded T cells using the weak-binding nested InsB_12-20_(C19S) peptide to maintain register specificity. Compared to studies using similar approaches^26,50,58^, this method resulted in only modest enrichment of InsB_12-20_(C19S)-specific CD4⁺ T cells and yielded a limited number of cells for transfer (Extended Data Fig. 5b). Despite this, diabetes development was significantly higher in NOD.*Rag1*^⁻/⁻^ recipients of enriched InsB_12-20_(C19S)-specific CD4^+^ T cells, although the incidence was incomplete (50%) (Extended Data Fig. 5c). Moreover, co-transfer of InsB_12-20_(C19S)-specific CD4⁺ T cells with polyclonal CD8⁺ T cells (purified from wildtype NOD mice) resulted in 100% diabetes incidence (Extended Data Fig. 5d). In contrast, CD8⁺ T cells alone, or CD8⁺ T cells co-transferred with CD4⁺ T cells enriched against a scrambled, non-cross-reactive, but immunogenic control peptide (Extended Data Fig. 5e,f), failed to induce diabetes (Extended Data Fig. 5d).

The adoptive transfer experiments suggest that InsB_12-20_(C19S)-specific CD4^+^ T cells can migrate to islets, mediate inflammation, and promote diabetes development by assisting cytotoxic CD8^+^ T cells. However, to further evaluate diabetogenic potential under physiological conditions, we decided to assess the peripheral activation and islet infiltration of InsB_12-20_(C19S)-specific CD4^+^ T cells along with several other CD4^+^ T cells with defined pathogenicity. First, we compared the peripheral activation of InsB_12-20_(C19S)-specific T cells to those recognizing three well-characterized islet neoepitopes: the deamidated insulin-1 C-peptide (CP1d; GDLQTLALEVARE), the 2.5HIP formed by C-peptide and ChgA (InsC-ChgA) (LQTLAL-WSRMD), and the InsC-IAPP 6.9HIP (LQTLAL-NAARD). I-A^g7^ tetramers for tracking CP1d-, 2.5HIP-, and 6.9HIP-reactive CD4⁺ T cells were produced using the same protocol as the InsB_12-20_(C19S):I-A^g7^ tetramer, as previously described^29^.

In SLOs of 8-week-old female NOD mice, all four tetramer-binding CD4⁺ T cell populations exhibited significantly higher proportions of CD44^hi^CD62L⁻ cells compared to tetramer-negative cells (Fig. 5f), confirming their peripheral activation. Notably, InsB_12-20_(C19S)-specific CD4⁺ T cells displayed an activation state even higher than that of CP1d-specific T cells (Fig. 5f), despite CP1d being a high-affinity ligand with ∼45-fold stronger I-A^g7^ binding than the native C-peptide^26^. Furthermore, InsB_12-20_(C19S)-specific T cell activation was similar to 2.5HIP- and 6.9HIP-reactive T cells (Fig. 5f), two contributors to T1D pathogenesis^12,58,59^.

Given the comparable activation of InsB_12-20_(C19S)-specific CD4⁺ T cells and pathogenic HIP-reactive T cells, we next examined whether InsB_12-20_(C19S)-specific T cells infiltrate pancreatic islets. We focused on an early disease stage to assess whether these cells participate in the initial waves of T cell infiltration. InsB_12-20_(C19S):I-A^g7^ tetramer-binding CD4⁺ T cells were readily detectable in the islets of 6-to 8-week-old female NOD mice, forming a population distinct from 2.5HIP- and 6.9HIP-reactive T cells (Fig. 5g). The frequencies of InsB_12-20_(C19S)-specific CD4⁺ T cells were consistently higher than those of 6.9HIP-reactive T cells (Fig. 5g), aligning with previous findings that 6.9HIP-reactive CD4⁺ T cells primarily accumulate in islets at later disease stages^59^. Moreover, the frequencies of InsB_12-20_(C19S)-specific T cells were similar to those of 2.5HIP-reactive T cells (Fig. 5g), which are known to drive early islet autoimmunity^12,58,59^. Nearly all intra-islet InsB_12-20_(C19S)- and 2.5HIP-specific CD4⁺ T cells were CD44^hi^CD62L^lo^ and CD44^hi^CD11a^hi^ (Fig. 5h), indicative of a terminally activated state in the islets.

### C19S transformation occurs in insulin in stressed and inflamed human islets

In our human PBMC HLA-II immunopeptidome analysis (Fig. 1), the detection of both native and C19S insulin peptides required MMTT stimulation. This observation suggests that C19S insulin is generated locally in human islets and can be released upon β-cell degranulation. To confirm whether C19S insulin is produced in human islets and assess how stress and inflammation influence this process, we sought to apply the GAP assay to human islets. Based on our previous isolation and validation of the crinosome and DCG subcellular fractions from human islets^19^, and given that the InsB_9-23_ segment is identical between mice and humans, we reasoned that offering human crinosomes and DCGs to C3.g7 APCs would allow for the detection of human insulin peptides (native or C19S) by the 9B9 and S5 T cell hybridomas.

We examined islet samples obtained from four non-diabetic donors (Supplementary Table 4). The islets were handpicked, treated with tunicamycin with or without glutathione for 2 hours, and then processed to crinosome and DCG isolation for testing by the GAP assay (Fig. 6a). In all four donors, tunicamycin treatment consistently increased the production of C19S insulin in both crinosome and DCG fractions compared to vehicle-treated controls (Fig. 6b). However, treatment with glutathione significantly inhibited tunicamycin-induced C19S generation, while leaving native insulin peptide levels unchanged (Fig. 6b). Thus, similar to results seen in MIN6 cells and mouse islets, ER and oxidative stress enhance C19S transformation in human islets.

**Figure 6.**
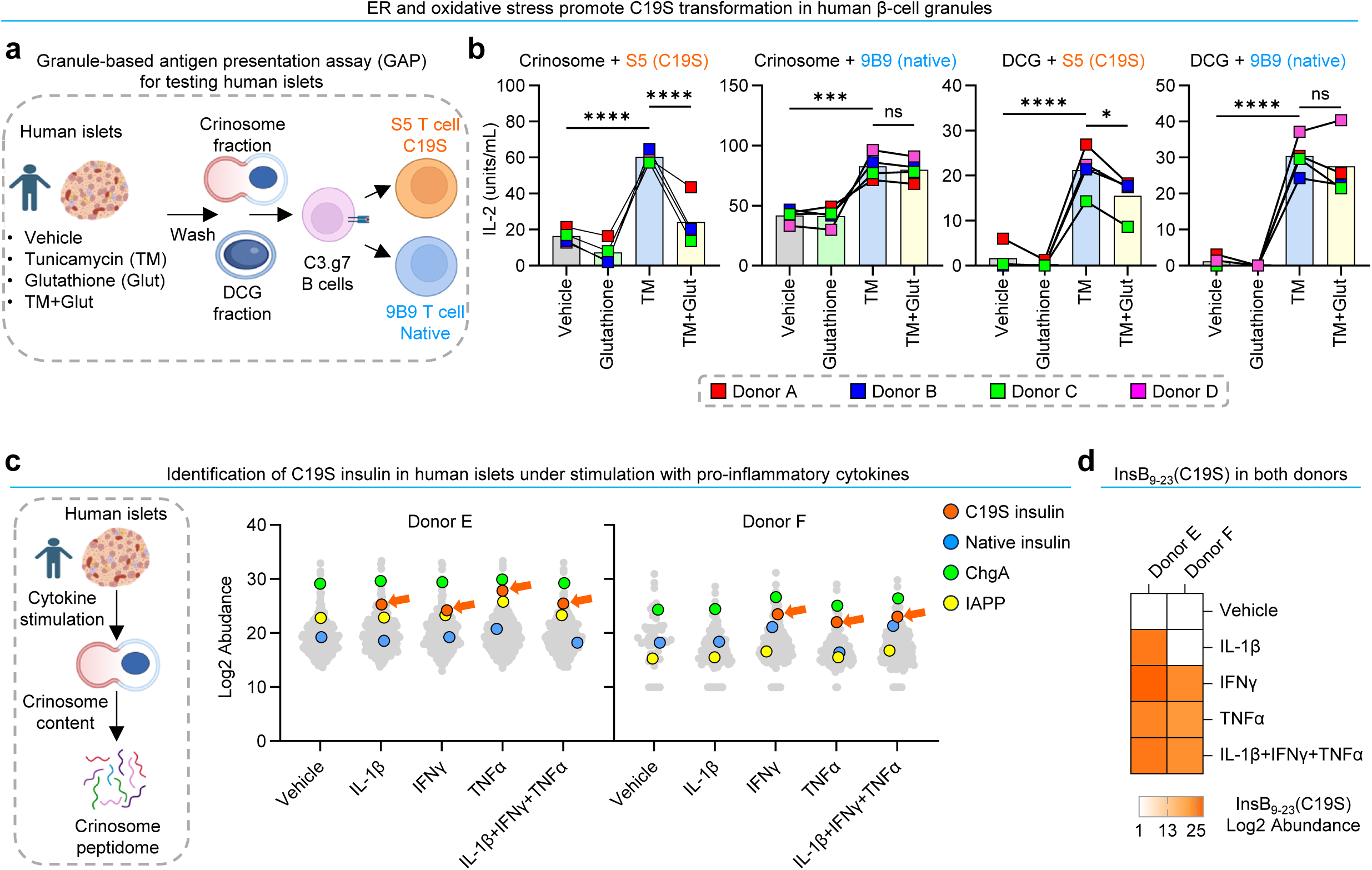
C19S transformation occurs in human islets during stress and inflammation. **a.** Schematic of the GAP assay using human islets as a source of β-cell granules. **b.** GAP assay showing the responses of the S5 and 9B9 T cells to crinosomes and DCGs isolated from human islets following indicated treatments. Data (mean ± SEM) summarize results from four donors (each point) examined in four independent experiments. ns, not significant; *P < 0.05; ***P < 0.001; ****P < 0.0001; repeated measures one-way ANOVA. **c.** Schematic (left) and identification (right) of C19S insulin peptides in crinosome peptidomes of human islets from two different donors exposed to indicated cytokines. Grey dots represent source proteins of all identified peptides, with native (blue) or C19S (orange) insulin, ChgA (green), and IAPP (yellow) indicated. **d.** A heatmap showing the appearance of the InsB_9-23_(C19S) peptide in the crinosome peptidomes of two human donors under different cytokine conditions.

To investigate whether inflammatory cytokines contribute to C19S transformation in human islets, we performed MS analysis of crinosome peptidomes isolated from human islets exposed to IL-1β, TNFα, IFNγ, and a combination of all three cytokines. Peptides from common β-cell proteins, including native insulin, ChgA, and IAPP, were identified in all conditions (Fig. 6c). With this unbiased approach, C19S insulin peptides were below the detection limit in vehicle-treated islets, possibly due to limited sample material. However, under cytokine stimulation conditions, including TNFα and IFNγ alone or TNFα, IFNγ, and IL-1β in combination, C19S insulin peptides were detected in crinosomes from both donors (Fig. 6c). Notably, we identified the InsB_9-23_(C19S) peptide in cytokine-stimulated human islets (Fig. 6d). These results collectively demonstrate that T1D-relevant stress and inflammatory signals act as conserved mechanisms to amplify C19S transformation in human islets.

### Expansion of HLA-DQ8-restricted, InsB_12-20_(C19S)-specific CD4^+^ T cells in T1D patients

Compared to NOD mice, human T1D involves a more diverse antigen repertoire, resulting in broader and more heterogeneous T cell responses. We previously showed in NOD mice that deamidation enhances the immunogenicity of C-peptide^26^; however, whether this PTM influences human C-peptide autoreactivity is currently under debate^60–62^, possibly due to notable sequence differences between mouse and human C-peptides. Recent advances have identified HIPs in human T1D, providing important insights into neoepitope-specific T cell responses^63–67^. Most human HIPs identified to date contain C-peptide segments and thus differ in composition, MHC-II presentation, and T cell specificity from HIPs identified in mouse studies. In contrast, the InsB_12-20_(C19S) epitope is identical between mice and humans and is generated through conserved, islet-intrinsic mechanisms. Given that I-A^g7^ shares similar structural and antigen presentation properties with human HLA-DQ8^68–70^, we examined whether HLA-DQ8-expressing individuals might harbor CD4⁺ T cells recognizing the InsB_12-20_(C19S) epitope during diabetes development.

We previously generated tetramer reagents to track HLA-DQ8-restricted CD4^+^ T cells specific for native InsB_12-20_ and InsB_13-21_ epitopes in humans at different stages of T1D^28^. Based on these studies, we produced HLA-DQ8-based tetramers incorporating the minimal 9-mer InsB_12-20_ or InsB_12-20_(C19S) register (Extended Data Fig. 6a) and used them to perform flow cytometry analysis on PBMC samples from three HLA-DQ8^+^ cohorts: non-diabetic controls (n = 7), individuals with recent-onset T1D (diagnosed within 12 months; n = 16), and individuals with established disease (diagnosed for > 5 years; n = 4) (Extended Data Fig. 6b). All participants in the three cohorts expressed at least one copy of the HLA-DQ8 haplotype (Supplementary Table 5).

Both the InsB_12-20_:DQ8 and InsB_12-20_(C19S):DQ8 tetramers exhibited minimal staining in CD4⁺ T cells from non-DQ8 healthy (DQ6/DQ7) or diabetic (DQ9/DQ5) individuals (Extended Data Fig. 6c), indicating their specificity for HLA-DQ8-restricted CD4^+^ T cells. As described previously^28^, the InsB_12-20_:DQ8 tetramer identified rare CD4^+^ T cells recognizing the native InsB_12-20_ epitope in DQ8^+^ individuals (Extended Data Fig. 6c). Some individuals occasionally showed a few diagonal events (Extended Data Fig. 6c). Notably, we identified a distinct population of CD4^+^ T cells labeled by the InsB_12-20_(C19S):DQ8 tetramer, which consistently appeared in all DQ8^+^ samples from the non-diabetic, recent-onset, and established cohorts, with little overlap with the InsB_12-20_-specific CD4^+^T cells (Extended Data Fig. 6c). To increase specificity, we performed dual-color staining using the InsB_12-20_(C19S):DQ8 tetramer conjugated with different fluorochromes, which also identified a discrete T cell population (Extended Data Fig. 6d). Collectively, these results demonstrate the presence of CD4^+^ T cells specific for the InsB_12-20_(C19S) epitope in DQ8^+^ individuals.

Next, we quantified InsB_12-20_:DQ8 and InsB_12-20_(C19S):DQ8 tetramer-binding CD4^+^ T cells in the three cohorts. As reported previously^28^, we observed a notable increase in InsB_12-20_-specific CD4^+^ T cells in patients with established T1D (Fig. 7a; Extended Data Fig. 6e). In contrast, InsB_12-20_(C19S)-specific CD4^+^ T cells were rare in nearly all non-diabetic individuals but showed significant expansion in patients with recent-onset or established T1D (Fig. 7a; Extended Data Fig. 6e). The numbers of these cells were notably variable among recent-onset patients, possibly reflecting heterogeneous activation at disease onset. To verify these results, we assessed recent-onset patients using dual-color tetramer staining, which confirmed the overall expansion trend of InsB_12-20_(C19S)-specific CD4^+^ T cells (Extended Data Fig. 6e). Furthermore, by the established stage, InsB_12-20_(C19S)-specific T cells were consistently more abundant and frequent than InsB_12-20_-specific T cells (Fig. 7a; Extended Data Fig. 6e). Collectively, these results demonstrate that InsB_12-20_(C19S)-specific CD4⁺ T cells expand significantly at diabetes onset and persist into the established stage, supporting their association with disease progression.

**Figure 7.**
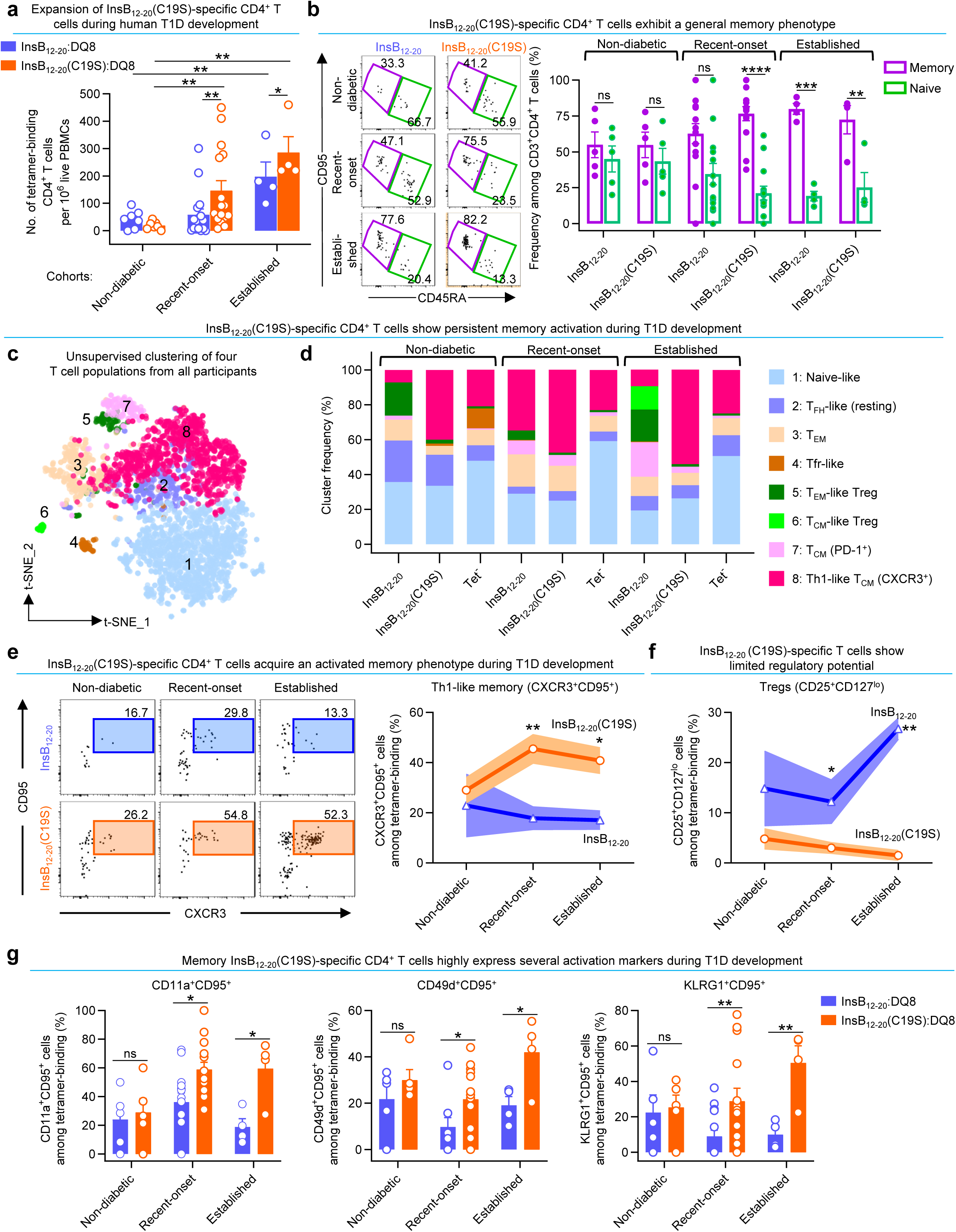
HLA-DQ8-restricted C19S-specific CD4^+^ T cells expand in T1D patients and acquire persistent memory activation during disease progression. **a.** Quantification of the numbers of indicated DQ8 tetramer-binding T cell populations from all 27 individuals across non-diabetic, recent-onset, and established cohorts. **b.** Representative FACS plots (left) and quantification (right) showing the distribution of naïve (CD45RA^+^CD95⁻) and memory (CD45RA⁻CD95^+^) cells within the indicated tetramer-binding CD4⁺ T cell populations in the three cohorts. **c.** A t-SNE plot showing eight CD4^+^ T cell clusters by unsupervised clustering of merged T cells from all 27 participants in the three cohorts. **d.** Distribution of each cluster in each of the four T cell populations across non-diabetic, recent-onset, and established cohorts. **e.** Representative FACS plots (left) and longitudinal quantification (right) showing CXCR3^+^CD95^+^ activated memory cells within CD127^hi^CD25⁻ conventional T cells among InsB_12-20_:DQ8 or InsB_12-20_(C19S):DQ8 tetramer-binding CD4⁺ T cells. **f.** Longitudinal quantification of CD25^+^CD127^lo^ Tregs within InsB_12-20_:DQ8 or InsB_12-20_(C19S):DQ8 tetramer-binding CD4⁺ T cells in the three cohorts. **g.** Frequencies of individual activation markers (CD11a, CD49d, and KLRG1) among CD95^+^ tetramer-binding CD4^+^ T cells. Data (**a, b, e, f, g**) summarize results from individual participants (each point) analyzed in 14 independent experiments. In **b, e, f**, **g,** samples without detectable InsB_12-20_:DQ8 or InsB_12-20_(C19S):DQ8 tetramer-binding CD4⁺ T cells were excluded from the analysis. ns, not significant; *P < 0.05; **P < 0.01; ***P < 0.001; ****P < 0.0001; Mann-Whitney test (**a**); two-tailed t test with Welch’s correction (**e, f, g**).

### Persistent memory activation of InsB_12-20_(C19S)-specific CD4⁺ T cells in human T1D

The observed expansion of InsB_12-20_(C19S)-specific CD4^+^ T cells during diabetes development led us to assess their phenotypic signatures. To this end, we designed a 27-marker spectral flow cytometry panel incorporating functional markers associated with activation, memory, and regulatory characteristics of human T cells. Given our previous detection of memory CD4⁺ T cells reactive to native insulin peptides^28^, we first asked whether InsB_12-20_(C19S)-specific CD4⁺ T cells exhibit a similar differentiation state. InsB_12-20_:DQ8 tetramer-binding CD4⁺ T cells in recent-onset patients showed a trend toward memory differentiation (CD95⁺CD45RA⁻), but this shift was not statistically significant and only became clearly evident in the patients with established T1D (Fig. 7b). In contrast, InsB_12-20_(C19S):DQ8 tetramer-binding CD4⁺ T cells exhibited a memory-dominant phenotype as early as the recent-onset stage, which persisted at similarly high levels in patients with established T1D (Fig. 7b).

Both InsB_12-20_- and InsB_12-20_(C19S)-specific CD4⁺ T cells exhibited a general memory phenotype, prompting us to explore whether they differed in their underlying activation or functional states. To this end, we combined InsB_12-20_:DQ8 and InsB_12-20_(C19S):DQ8 tetramer-binding cells, along with tetramer-negative controls, from all 27 individuals in the non-diabetic, recent-onset, and established cohorts, and performed unsupervised clustering of the merged samples. This analysis revealed 8 clusters (Fig. 7c) capturing the well-documented cellular diversity in autoreactive T cells involved in T1D^71^.

Cluster 1 was characterized by high expression of CD45RA and CCR7 (Extended Data Fig. 6f), consistent with a naïve-like T cell phenotype. Cluster 2 lacked memory and Treg markers but expressed CXCR5 and low levels of PD-1 (Extended Data Fig. 6f), a profile compatible with previously described circulating CXCR5^+^ T_FH_ cells in a resting state^72^.

Clusters 4, 5, and 6 exhibited a general Treg phenotype (CD25^+^CD127^lo^) but differed in their differentiation status. Cluster 4 expressed CXCR5, along with memory markers CD45RO and CD95 (Extended Data Fig. 6f), consistent with circulating memory-like T follicular regulatory (Tfr) cells described in humans^73^. Cluster 5 displayed an effector memory (T_EM_) profile (CD45RO^+^CD95^+^CCR4^+^CD45RA⁻CCR7⁻) (Extended Data Fig. 6f), resembling T_EM_-like Tregs^74^; its high expression of HLA-DR corresponded to previously reported DR^+^ Tregs capable of inducing potent and rapid suppression^75^. Cluster 6, by contrast, exhibited reduced CD25 expression, a hallmark of terminally differentiated CD25^low^ Tregs previously shown to expand in autoimmune patients^76^. This population (CD45RO^+^CD95^+^CCR4^+^CD45RA⁻CCR7^+^) also resembled central memory (T_CM_)-like Tregs and expressed high levels of PD-1 and CD226 (Extended Data Fig. 6f), two markers associated with Treg persistence and regulatory function^77,78^.

Clusters 3, 7, and 8 exhibited a general memory (CD45RO^+^CD95^+^CD45RA⁻) signature while lacking Treg markers (Extended Data Fig. 7c). Cluster 3 lacked CCR7 and preferentially expressed KLRG1 and CXCR3, consistent with an activated T_EM_ profile. Clusters 7 and 8 both retained CCR7, consistent with a T_CM_ phenotype and expressed activation markers CD11a, CD49d, KLRG1, CD28, CD38, and CXCR3 (Extended Data Fig. 6f). However, cluster 7 had higher expression of PD-1, suggesting restrained activation. In contrast, cluster 8 had higher levels of CXCR3, consistent with a Th1-like T_CM_ activation profile. This feature aligns with recent scRNA-seq analysis defining T_CM_ activation in islet-reactive CD4^+^ T cells from T1D patients^79^.

Analysis of cluster distributions revealed several distinct features of InsB_12-20_(C19S)-specific CD4⁺ T cells. Compared to the other two populations, these cells were more enriched in the Th1-like T_CM_ cluster 8 (Fig. 7d). In both recent-onset and established patients, cluster 8 accounted for more than half of the InsB_12-20_(C19S)-specific CD4^+^ T cells (Fig. 7d), indicating persistent skewing toward a Th1-like memory phenotype.

On the other hand, InsB_12-20_(C19S)-specific CD4^+^ T cells showed minimal representation in either of the two Treg-associated clusters, T_EM_-like cluster 5 and T_CM_-like cluster 6, across the non-diabetic, recent-onset, and established disease stages (Fig. 7d). By comparison, cluster 5 was already detectable in InsB_12-20_-specific CD4⁺ T cells at the non-diabetic stage and remained evident in established disease, while cluster 6 was only seen in these cells at the established T1D stage (Fig. 7d).

To validate the phenotypic features identified by unsupervised clustering, we analyzed CXCR3^+^CD95^+^ Th1-like memory cells in individual subjects. Frequencies of CXCR3⁺CD95⁺ cells were comparable between the two specificities in non-diabetic individuals. However, InsB_12-20_(C19S)-specific T cells showed a marked increase in this population by the recent-onset stage, which persisted at high levels in patients with established T1D (Fig. 7e). This trend was not observed in T cells reactive to the native epitope (Fig. 7e). In contrast, CD25⁺CD127^lo^ Tregs showed an opposite pattern: these cells were markedly fewer overall and further declined within the InsB_12-20_(C19S)-specific population across the stages (Fig. 7f). We also quantified additional activation markers associated with the Th1-like T_CM_ cluster, including CD11a, KLRG1, and CD49d, at the individual level. All three markers were significantly highly expressed in InsB_12-20_(C19S)-specific CD4^+^ T cells at both recent-onset and established disease stages (Fig. 7g). The activation levels appeared more heterogenous among recent-onset patients, again reflecting individual variation in autoimmune activities at diabetes onset. Collectively, these results highlight the distinct functional profile of InsB_12-20_(C19S)-specific T cells, characterized by progressive memory activation and limited regulatory potential, that emerges early and persists throughout the course of T1D development.

## DISCUSSION

Our study identifies a new mechanism of neoantigen formation in T1D, in which disease-relevant signals induce and amplify a Cys→Ser transformation (C19S) in insulin, leading to neoepitope presentation and CD4^+^ T cell autoreactivity. We detected C19S insulin peptides in both human and mouse MHC-II immunopeptidomes, demonstrating that this transformation occurs in vivo across species and integrates into the broad self-peptidome. Inflammation and oxidative stress enhance C19S transformation in β cells and APCs, underscoring the role of the islet microenvironment in shaping the autoimmune peptidome. We further show that C19S-specific CD4⁺ T cells expand in both NOD mice and HLA-DQ8⁺ T1D patients, displaying an activated memory phenotype linked to disease progression. In summary, C19S represents an extra-genomic, microenvironment-driven transformation that diversifies β-cell antigenicity at the single residue level.

A distinctive feature of the C19S transformation is its strong association with the physiological and pathological islet microenvironment. In human PBMC MHC-II immunopeptidomes, detection of C19S insulin peptides required prior MMTT stimulation, suggesting that C19S arises from β-cell-derived peptides released upon degranulation rather than being a passive byproduct of systemic insulin processing. This was further supported by GAP assay experiments, which confirmed the presence of C19S insulin peptides in β-cell granules in both mice and humans. Notably, at steady state, C19S peptides were predominantly localized within crinophagic granules (crinosomes) rather than DCGs. This suggests two possibilities: C19S transformation may occur directly within crinosomes where degraded insulin peptides are processed, or misfolded C19S insulin products packaged in DCGs may be selectively delivered to crinosomes for degradation. In the context of T1D, we previously detected low levels of native insulin peptides and HIPs in crinosomes under non-autoimmune conditions^19,26,29,80^, indicating that these peptides can be generated independently of immune activation. Similarly, C19S insulin peptides are also produced before autoimmunity occurs, supporting the emerging concept that intrinsic properties of pancreatic islets, particularly the highly oxidative islet microenvironment, may inadvertently contribute to the initiation of diabetic autoimmunity^55,81^.

Under pathological conditions, C19S transformation is markedly amplified, coinciding with increased intra-islet MHC-II presentation as diabetes progresses. We show that multiple T1D-relevant signals, including ER stress, oxidative stress, and inflammatory cytokines, drive this transformation. ER stress and inflammatory cytokines are well known to generate a range of PTMs in β cells^15^, and our recent work showed a role of ER stress in promoting 6.9HIP production in β-cell granules^80^. Although these pathological signals involve complex pathways, blocking oxidation largely inhibited C19S transformation under both ER stress and cytokine stimulation. This tight link between C19S transformation and oxidative stress supports existing evidence that inhibiting ER and redox stress can mitigate T1D progression^41,82–86^. Furthermore, C19S transformation is not confined to β cells but can also occur in cytokine-activated DCs. This feature distinguishes C19S from other neoantigen-generating mechanisms that are confined to tissue-resident cells, thereby contributing to insulin epitope spreading during T1D progression.

C19S as described here is best considered a single amino acid transformation, since it was detected through a SPIDER search rather than a conventional PTM search. Mechanistically, Cys →Ser also differs from PTMs that are enzyme-mediated and reversible, whereas prior work showed that oxidation can drive cysteine thiols through dehydroalanine intermediates to serine in an irreversible process^38^. A seminal study demonstrated that oxidative stress can also drive post-genetic recoding through methionine misacylation of non-cognate tRNAs, altering protein translation without involving classical PTMs^87^. Our study extends these concepts to β cells in vivo, showing that oxidative stress can generate C19S in the insulin B-chain, creating a disease-relevant neoepitope. Several features of insulin and the β-cell environment, its cysteine-rich structure, the redox sensitivity of B(C19) residue, and the intrinsically oxidative islet milieu converge to create a “perfect storm” for this transformation. Our recent immunopeptidome analysis identified native InsB_9–23_ as an MHC-II–bound peptide in the thymus^29^. After exhaustive searching, we did not detect the C19S variant, supporting the idea that C19S formation requires an oxidative tissue microenvironment such as pancreatic islets. Given the essential role of the B(C19) residue in insulin biosynthesis, C19S likely represents collateral damage rather than an adaptive modification. In a broader sense, our study suggests that Cys→Ser transformations may also occur in other disease contexts where ROS are elevated, such as viral infections and oxidative tumor microenvironments. Immunopeptidome analysis will be a powerful approach to indicate whether such peptides are presented by MHC-I or MHC-II molecules and whether they can act as neoantigens.

In the context of insulin autoreactivity, the previously described defective ribosomal insulin gene product (DRiP) in humans provides a remarkable example of how non-canonical translation events can generate highly immunogenic neoepitopes^88^. While the DRiP-derived epitope appears to be unique to humans, our data show that C19S transformation is a conserved process, generating identical neoepitopes in both mice and humans. In NOD mice, we found that the InsB_12-20_(C19S) epitope is recognized by a distinct CD4^+^ T cell population specific for the single amino acid C19S transformation. Although this differs from classical altered peptide ligand (APL) recognition^89^, which usually involves the same TCR responding differently to distinct peptide ligands, it illustrates the broader principle that a single-amino acid change in a self-peptide can profoundly alter T cell function. In this case, the largely non-overlapping repertoire responding to C19S is consistent with recognition of a neoantigen. An intriguing observation is the inactivation of the InsB_12-20_(C19S)-specific CD4^+^ T cells in the NOD.*B16A* mice, suggesting that the loss of reactivity to the C19S epitope may also contribute to the profound diabetes resistance phenotype in this mouse^23^. Along these lines, previous studies have indicated that C-terminal additions^90^ and peptide fusion^91^ of InsB_9-23_ can generate strong TCR agonists. The identification of C19S further highlights that autoreactivity to the InsB_9-23_ region manifests in multiple forms, and this diversity may explain why this segment remains a dominant target in autoimmune diabetes.

In humans, InsB_12-20_(C19S)-specific CD4⁺ T cells expand at diabetes onset, with numbers significantly increased compared to DQ8⁺ at-risk controls. Notably, these cells exhibit a T_CM_ activation phenotype, emerging early in at-risk individuals and persisting into the long-term established disease stage. Thus, independent of the NOD mouse data showing islet infiltration and terminal activation, the expansion and sustained activation of C19S-specific T_CM_ cells in humans suggest the pathological relevance of C19S transformation in T1D development. From a biomarker perspective, tracking C19S-specific T_CM_ cells could offer a dynamic measure of disease activity, progression risk, and therapeutic efficacy, aligning with the concept that antigen-specific CD4⁺ T cells can be used to monitor islet autoimmunity in real time^28^. Moreover, given the self-renewal capacity and tissue-homing potential of T_CM_ cells, C19S-specific T cells may readily target newly regenerated or transplanted β cells in patients with established T1D. Developing strategies to modulate these T_CM_ cells may have therapeutic implications in islet transplantation. Overall, our study suggests that C19S-specific CD4⁺ T_CM_ cells are primed for long-term persistence and may act as a sustained reservoir of anti-islet autoreactivity. Their persistence and reactivation capacity make them a potential target for immunotherapies aimed at disrupting pathogenic memory responses and restoring immune tolerance.

### Limitation of this study

Due to logistical constraints of MMTT and multiple blood draws required for juvenile participants in the human immunopeptidome study, we had limited sample materials to perform HLA-DQ8-or HLA-DQ2-specific peptidome isolation. The identification of C19S insulin peptides in the pan-HLA-DR peptidome suggests that these peptides may be presented by multiple HLA-II molecules, an area warranting further investigation. In all our peptidome analyses (MHC-II or soluble peptidomes), we applied stringent filtering criteria to enhance data fidelity. This approach may have inadvertently missed SAVs in other β-cell proteins that are less abundant than insulin. Also, the use of a short-term ER stress model may explain why certain redox-associated PTMs (i.e., deamidation, citrullination, carbonylation) did not show substantial changes in the MIN6 peptidome experiments. We unexpectedly found that the Y16A mutation also nullified the reactivity of the C19S epitope in the NOD.*B16A* mice. However, determining the contribution of C19S reactivity relative to the native insulin epitope remains challenging. Definitive evidence may require selectively ablating C19S transformation while preserving native insulin structure, but this is technically difficult since manipulating the B(C19) residue may disrupt normal insulin biosynthesis and β-cell function. In this regard, we are developing new reagents that may allow for specific tracking of C19S insulin peptides, based on our experience in producing peptide-specific monoclonal antibodies^19,20^. These tools may help define the localization of C19S insulin peptides in inflamed β cells and APCs and inform their roles in T1D.

## Acknowledgements

We extend our gratitude to Kodi S. Ravichandran and Jonathan Kipnis for their insightful guidance throughout this study and for their critical review of the manuscript. We thank Jennifer Ponce, Christelle Schatz, and Jinsheng Yu for their expertise in performing scRNA-seq analysis, as well as Dorjan Brinja and Erica Lantelme for their assistance with cell sorting. We acknowledge the dedicated efforts of the team at St. Louis Children’s Hospital in recruiting human subjects for this study and express our sincere appreciation to all the participants and their families. We thank Tina Turicek and Lucy Turicek for their assistance in preparing the manuscript. This work was supported by the Diabetes Research Center at Washington University (P30 DK020579 to X.W.), the Juvenile Diabetes Research Foundation (5-CDA-2022-1175-A-N to X.W.), and the National Institutes of Health (R01AI162591 and R01DK134437 to X.W.).

## Author Contributions

N.S. designed and conducted in vivo immunoassays and T cell biology experiments. A.N.V. carried out antigen presentation experiments and isolated immunopeptidomes. Y.R. performed T cell biology experiments and spectral flow cytometry analysis. O.J.P. prepared pancreatic islets and assisted in multiple experimental procedures. Y.Y., D.P.T., O.A., A.Z., and H.H. assisted various T cell biology experiments and manuscript writing. T.L. and B.Z. provided advice on the design of the scRNA-seq analysis and performed data analysis. L.K. and L.T. developed and produced I-A^g7^- and DQ8-based tetramers. C.C.C. and L.S. designed and supplied samples for the lymph peptidome experiments. A.V.J. provided critical advice on TCR analysis. S.S., R.M., and L.C. offered crucial advice on the human spectral flow cytometry analysis. P.S., and A.A. recruited human subjects. E.R.U. inspired us to search for non-canonical peptides in crinophagic granules and instructed on applying the GAP assay to human islets. A.M.A. supervised human subject recruitment and conducted clinical studies. C.F.L. performed mass spectrometry experiments and analyzed data. X.W. analyzed data and designed the study. A.V.J., L.C., C.-S.H., L.S., L.T., and A.M.A. critically evaluated data analyses and the manuscript. N.S., C.F.L., and X.W. wrote the manuscript.

## METHODS

### Mice

NOD/ShiLtJ (NOD), NOD.129S7(B6)-*Rag1*^tm1Mom^/J (NOD.*Rag1*^⁻/⁻^), NOD.Cg-Tg (Ins2*Y16A) 1EllIns1t^m1Jja^Ins2^tm1Jja^/GseJ (NOD.*B16A*), NOD.Tnfrsf1a/1b^⁻/⁻^, and C57BL/6 mice were originally obtained from The Jackson Laboratory. All mice were bred, maintained, and used in experiments in our specific pathogen-free (SPF) animal facility in accordance with the Division of Comparative Medicine at Washington University School of Medicine (Association for Assessment and Accreditation of Laboratory Animal Care accreditation no. A3381-01; Protocol No. 23-0429).

### Antibodies

The following antibodies were used for mouse studies: anti-CD45 (30-F11), anti-CD11c (N418), anti-CD11b (M1/70), anti-CD3ε (145-2C11), anti-CD4 (RM4-5), anti-CD8α (53-6.7), anti-CD44 (IM7), anti-CD90.2 (53-2.1), anti-B220 (RA3-6B2), anti-CXCR6 (SA051D1), anti-CD62L (MEL-14), anti-CD69 (H1.2F3), and anti-CD19 (6D5), were all purchased from BioLegend. Anti-CD11a (M17/4) was purchased from eBioscience (ThermoFisher Scientific). The anti-I-A^g7^ antibody (AG2.42.7) was generated in our laboratory.

The following antibodies were used for human studies: anti-CD19 (HIB19), anti-CD14 (HCD14), anti-CD3 (SK7), anti-CD8 (SK1), anti-CCR7 (G043H7), anti-CXCR3 (G025H7), anti-CD25 (BC96), anti-HLA-DR (L243), anti-CD28 (CD28.2), anti-CD95 (DX2), anti-PD-1 (EH12.2H7), anti-CD45RO (UCHL1), anti-CD38 (S17015A), and anti-CXCR5 (J252D4), were all purchased from BioLegend. Anti-CD4 (SK3), anti-CD45RA (HI100), anti-KLRG1 (13F12F2), and anti-CD226 (11A8.7.4) were purchased from eBioscience (ThermoFisher Scientific). Anti-CD127 (HIL-7R-M21), anti-CD27 (M-T271), anti-CCR6 (11A9), anti-CCR4 (1G1), anti-CD11a (G25.2), and anti-CD49d (9F10) antibodies were purchased from BD Biosciences.

### Human subject recruitment

All human participants were recruited through St. Louis Children’s Hospital at Washington University School of Medicine under protocols approved by the Institutional Review Board (IRB) at Washington University in St. Louis. Written informed consent was obtained from all participants or their legal guardians.

For the HLA-II immunopeptidome and MMTT study, 30 participants were enrolled over a four-year period, with 10 participants per cohort: (1) non-diabetic controls (n=10); (2) T1D patients at 3-month onset (n=10); and (3) T1D patients at 18-month onset (n=10). Some non-diabetic participants were siblings of individuals with T1D.

For the human T cell biology studies, a separate recruitment was conducted over a three-year period and included three cohorts: (1) non-diabetic controls, (2) recent-onset T1D patients (diagnosed within 12 months), and (3) individuals with established T1D (diagnosed for >5 years). More than 65 individuals were screened for HLA-DQ haplotypes, and PBMC samples from participants expressing at least one copy of the HLA-DQ8 allele were used for the study. Because only two DQ8⁺ individuals were identified among recruited non-diabetic controls, an additional five DQ8⁺ samples were obtained from Precision for Medicine, Inc. In total, 27 samples were included in the T cell biology experiments: non-diabetic (n=7), recent-onset (n=16), and established (n=4).

### Human HLA typing and sample size

High-resolution HLA class I and class II typing for participants in the immunopeptidome study was performed by Histogenetics (Ossining, NY) using ∼1 mL of whole blood. For the T cell biology study, HLA-DQ typing was conducted at the HLA Laboratory at Washington University School of Medicine using ∼0.5 mL of whole blood.

The immunopeptidomics component of this study was exploratory in nature, and sample size was guided by practical and technical considerations. Due to logistical constraints associated with the MMTT and repeated blood draws, approximately 3-5 mL of blood was collected per participant per time point. By pooling samples from 10 individuals per cohort, we estimated a yield of ∼3-12 × 10⁶ HLA-II⁺ cells per group. Based on our experience, this yield fell within a workable range enabling detection of representative β-cell-derived peptides in the HLA-II peptidome.

For the human T cell biology study, sample size was first guided by our prior publication^28^ and the mouse studies in the present study. Based on our experimental design comparing activation phenotypes between C19S- and native insulin-specific CD4⁺ T cells, we performed a power analysis using a two-sided t-test (α = 0.05, power = 0.80). Assuming a threefold difference in activation marker frequency and a standard deviation of 15-20%, the resulting effect size (Cohen’s d = 1.5-2.0) indicated that 5-8 subjects per group would be sufficient to detect this difference with 80% power. Our final cohort included 7 non-diabetic controls, 16 recent-onset patients, and 4 individuals with established T1D, and yielded statistically significant results.

### Human PBMC immunopeptidome analysis

After fasting for at least 8 hours, each participant had a baseline blood drawn (time 0) and was given an MMTT by drinking Boost High Protein Nutritional Energy Drink (Mead-Johnson) at 6 mL/kg (maximum 360 mL). Additional blood draws were performed at 90 and 120 minutes post-MMTT. C-peptide levels were measured 10 minutes before and immediately before MMTT, as well as 15, 30, 60, 90, and 120 minutes post-MMTT. All subjects had their glucose measured upon arrival at the study site; ND controls with glucose levels > 126 mg/dL and T1D patients with glucose levels > 250 mg/dL did not continue the study. T1D subjects also avoided doses of rapid-acting insulin 8 hours before testing unless needed to correct significant hyperglycemia.

For each participant (ND, 3mos, 18mos, n=10 per group), three blood samples (time 0, 90, 120 minutes) were collected in BD P800 Blood Collection System tubes (BD Biosciences, Cat #366421) containing a cocktail of protease inhibitors. An aliquot of whole blood (1 mL) was used for HLA typing by HistoGenetics. The rest was used to isolate PBMCs (buffy coats), which were then frozen as cell pellets at -80°C.

For sequential isolation of the HLA-DQ and HLA-DR peptidomes, the frozen cell pellets from each time point of each patient were thawed on the isolation day, pooled based on their conditions (ND, 3mos, 18mos), and suspended in fresh lysis buffer (40 mM MEGA 8 (MilliporeSigma, Cat #: O3129), 40 mM MEGA 9 (MilliporeSigma, Cat #: N1138), 1 mM phenylmethylsulfonyl fluoride (MilliporeSigma, Cat #: P7626), 0.2 mM iodoacetamide (MilliporeSigma, Cat #: I6125), 20 µg/mL leupeptin (MilliporeSigma, Cat #: L2884), and Roche Complete Mini Protease cocktail (Roche Diagnostics, Cat #: 11836153001) in PBS). There were a total of 18 samples, each containing 35 to 51 million PBMCs. The lysate suspensions were rocked for 2 hours at 4 °C and then centrifuged at 20,000xg for 30 minutes at 4 °C. To eliminate non-specific binding of peptides, the supernatant was first incubated with polyclonal mouse IgG (Leinco, Cat #: N229; 1.5 mg antibody per sample) bound to Sepharose 4B (MilliporeSigma, Cat #: C9142) at 4 °C for 30 minutes. The unbound fraction containing peptide-MHC-II complexes was collected and added to a tube containing PBS-washed Sepharose conjugated to the anti-pan-HLA-DQ antibody (Leinco, Cat # H262, 3.0 mg per sample). This mixture was incubated at 4 °C overnight. The HLA-DQ Sepharose was applied to a column and the flow-through portion was collected and incubated with the anti-pan-HLA-DR antibody (Leinco, Cat # H261, 3.0 mg per sample) conjugated with Sepharose at 4 °C overnight. Either HLA-DQ or HLA-DR Sepharose was washed four times as follows: 10 mL 150 mM NaCl and 20 mM Tris (pH 7.4), 10 mL 400 mM NaCl and 20 mM Tris (pH 7.4), 10 mL 150 mM NaCl and 20 mM Tris (pH 7.4), and 10 mL 20 mM Tris (pH 8.0). Peptides were eluted with 10% acetic acid (ThermoFisher, Cat #: A38SI-212) and dried using a SpeedVac. Eluted peptides were resuspended and passed through detergent removal spin columns (Pierce, Cat #: 87777) to remove traces of remaining detergent, and then cleaned using C18 Spin Columns (Pierce, Cat #: 89870).

### Generation of CD4^+^ T cell hybridomas

NOD mice (5-8-weeks old) were immunized with the InsB_9-23_(C19S) peptide (10 nmol) emulsified in Complete Freund’s Adjuvant (CFA; Difco) subcutaneously in the footpads of the hind legs. One week following immunization, the popliteal lymph nodes were collected and dispersed into single-cell suspensions. The cells were then boosted with 1 μM of the InsB_9-23_(C19S) peptide for 3 days and fused with the BW5147 fusion partner following standard protocols. The growth-positive T cell clones were screened against the InsB_9-23_(C19S) peptide using an antigen presentation assay. The antigen-responsive clones were further expanded and subcloned to generate monoclonal T cell hybridomas.

### Competition binding assay

For the cell-based binding assay, C3.g7 cells (10^4^) were mixed with competitor peptide and incubated for 30 minutes at 37°C. The HEL:11-25 peptide (1 µM) was added to the C3.g7/competitor peptide mixture and incubated for an additional 60 minutes at 37°C. Cells were centrifuged and washed. A CD4^+^ T cell hybridoma specific for the HEL:11-25 peptide (clone 10E11) was added to the washed and loaded C3.g7 cells and incubated overnight at 37°C. The supernatants were then collected, and the production of IL-2 was measured by ELISA. IC50 values for competitor peptides were generated by fitting the data using GraphPad Prism software as a four-parameter inhibitor vs. response curve. For the biochemical binding assay, peptides were diluted in PBS and then added to the reaction mix to provide a range between 5-1000 pmoles peptide/sample. For each sample, 0.1 U thrombin, 1 µg of soluble g7-clip, and 0.125 pmoles of 125I-labeled g7-MIME peptide were added to the diluted unlabeled peptides. The reaction was allowed to sit at room temperature overnight. BioGel P6 columns (BioRad, Cat # 732-6299) were used to separate unbound, labeled peptide, and the fraction of displaced labeled peptide was quantified with a gamma counter (Perkin Elmer, Wallac 1272 CliniGamma). Binding curves were determined by plotting the amount of displaced labeled peptide vs the concentration of unlabeled competitor. The binding data is reported as the concentration of unlabeled peptide required to reduce g7-MIME binding by 50%.

### Isolation of pancreatic islets

To isolate pancreatic islets, the peritoneal cavity was opened to expose the common bile duct. Using a dissection microscope, the bile duct leading to the duodenum was clamped. A solution of type XI collagenase (0.4 mg/mL; Sigma-Aldrich) in isolation buffer (composed of 1x HBSS, 10 mM HEPES, and 1 mM MgCl2; pH 7.4) was injected through the common bile duct to perfuse the pancreas. The inflated pancreas was then carefully removed and digested at 37°C for 12-14 minutes. Crude islets were collected and repeatedly washed with wash buffer (1x HBSS, 10 mM HEPES, 1 mM MgCl2, 1 mM CaCl2; pH 7.4). Under a microscope, pure islets were hand-picked, excluding any acinar tissue. The purified islets were then dispersed into a single-cell suspension using a non-enzymatic cell dissociation solution (Sigma-Aldrich) for 3 minutes at 37°C.

### ELISPOT

For evaluating T cell responses by immunization, mice were immunized with the indicated peptides (10 nmol) emulsified with CFA at the footpad. On day 7, the popliteal lymph node cells were restimulated with the indicated peptides (10 μM) for 24 hours, and IFNγ production was assessed by ELISPOT. To evaluate T cell responses in islets and pancreatic lymph nodes without immunization, total islet and pLN cells were harvested from 8-week-old female NOD mice, pooled, and cultured with a mixture of equal amounts of InsB_9-23_ and InsB_9-23_(C19S) peptide (1 µM), in the presence of IL-2 (20 U/mL) for 7 days. Total live cells were collected from the culture using Histopaque 1119. The live cells were added to a culture containing 20 U/mL IL-2 and 1 µM peptide as before, along with 2 × 10^6^ irradiated NOD splenocytes (3000 RAD) per mL of culture, and incubated for 3 days at 37°C. In all experiments, cells were harvested and assayed for reactivity by eliciting a recall response on IFNγ-coated 96-well multi-screen plates (Merck Millipore, Cat #: S2EM004M99) for ELISPOT. The reactive cells were counted using Immunospot Software (C.T.L.).

### T cell expansion and adoptive transfer

NOD mice were immunized with either InsB_9-23_(C19S) or scrambled peptide emulsified with CFA. Primary T cell lines were generated by culturing popliteal lymph node cells with 1 μM of either peptide in the presence of 20 U/mL of IL-2 and 2 × 10^6^/mL of irradiated splenocytes. After one week of culture, total live cells were collected using Histopaque-1119. The cells underwent four cycles of expansion with the nested InsB_12-20_(C19S) or scrambled control peptide for four weeks, each week supplemented with 0.1 μM peptide, 20 U/mL of IL-2 and 2 × 10^6^/mL of irradiated splenocytes. Due to the weak binding of the InsB_12-20_(C19S) peptide, only 5-8% percent of CD4^+^ T cells were tetramer-positive, roughly tenfold lower than what has been reported for strong peptide agonists^58^. We also found that post-expansion, these tetramer-binding CD4^+^ T cells are vulnerable to cell death during FACS sorting. A total of 1 × 10^6^ CD4^+^ T cells containing about 5 × 10^4^ InsB_12-20_(C19S)-specific CD4^+^ T cells were transferred intravenously per NOD.*Rag1*^⁻/⁻^ recipient. For CD8^+^ T cell co-transfer, polyclonal CD8^+^ T cells were purified using anti-mouse CD8 isolation magnetic beads (Miltenyi Biotec), and 1 × 10^6^ CD8^+^ T cells were transferred intravenously into NOD.*Rag1*^⁻/⁻^ recipients. Urine glucose levels in mice were checked weekly (AimStrip US-G; Germaine Laboratories) and those with a level ≥250 mg/dL for two consecutive readings were considered diabetic.

### I-A^g7^ tetramer staining using magnetic enrichment

The generation of I-A^g7^-based monomers has been previously described^27^. Biotinylated monomers were tetramerized by incubating them with APC- or PE-labeled streptavidin (Agilent Technologies, Cat #: PJ27S and PJRS301-1) at a 5:1 molar ratio of biotinylated molecules to labeled streptavidin for 1 hour at room temperature. Single-cell suspensions from the spleen and lymph nodes (inguinal, axillary, and pancreatic lymph nodes) were washed with PBS and incubated with the APC- or PE-labeled tetramers at a final concentration of 10 μg/mL for 1 hour at room temperature. Following incubation, the cells were washed twice with MACS buffer (2 mM EDTA, 0.5% BSA in PBS) and then incubated with 20 μl of anti-APC and anti-PE microbeads (Miltenyi Biotec, Cat #: 130-090-855 and 130-048-801) in 100 μl of MACS buffer at 4°C for 20 minutes. After washing, the cells were resuspended in 1 mL of MACS buffer and applied to LS columns (Miltenyi Biotec, Cat #: 130-042-401) for enriching tetramer-bound T cells. Both the flow-through (negative fraction) and column-bound cells (positive fraction) were collected for subsequent flow cytometry analysis.

### Flow cytometry analysis

Single-cell suspensions from spleen and lymph nodes (inguinal, axillary, and pancreatic lymph node) were prepared by mechanically disrupting the tissue using a syringe plunger through a 70-µm filter. Red blood cells were lysed, and the cell suspension was washed with PBS, and then stained with tetramers. Following tetramer staining, cells were washed with FACS buffer (1% BSA in 1xPBS) and incubated with FcR blocking antibody (2.4G2) for 10 minutes at 4°C. Surface staining was performed with fluorescently labeled antibodies (1:200 v/v) by incubating at 4°C for 30 minutes. Cells were washed twice with FACS buffer. Samples were examined using BD FACS Canto II Cell Analyzer or BD FACS Symphony Cell Analyzer (BD Biosciences). Data was analyzed using FlowJo 10.9.0 software (TreeStar).

### Cell sorting and library preparation for single-cell RNA sequencing analysis

For scRNA-seq experiments, spleen and lymph node cells were pooled from 8-week-old female NOD mice (n=6-10). After red blood cell lysis, cells were washed with MACS buffer, and the CD19^+^ B cells and CD11b^+^ myeloid cells were depleted using CD19 beads (Miltenyi Biotec, Cat #: 130-050-301) and CD11b beads (Miltenyi Biotec, Cat#: 130-097-142). Cells were then stained with fluorochrome-conjugated tetramers followed by magnetic enrichment using anti-PE and anti-APC beads. Surface staining was performed as described above and tetramer-positive and polyclonal cells were sorted using the BD FACS Aria II sorter (BD Biosciences).

cDNA was prepared after the GEM generation and barcoding, followed by the GEM-RT reaction and bead cleanup steps. Purified cDNA was amplified for 11-16 cycles before being cleaned up using SPRI Select beads. Samples were then run on a Bioanalyzer to determine the cDNA concentration. V(D)J target enrichment (TCR) was performed on the full-length cDNA. Gene Expression, Enriched TCR, and Feature libraries were prepared according to the recommended protocols from the 10× Genomics Chromium Single Cell 5′ Reagent Kits User Guide (v2 Chemistry Dual Index) with Feature Barcoding technology for Cell Surface Protein and Immune Receptor Mapping, with appropriate modifications to the PCR cycles based on the calculated cDNA concentration. For sample preparation on the 10× Genomics platform, the following kits were used: Chromium Next GEM Single Cell 5′ Kit v2, 16 rxns (Cat #: PN-1000263), Chromium Next GEMChip K Single Cell Kit, 48 rxns (Cat #: PN-1000286), Chromium Single Cell Human TCR Amplification Kit (Cat #: PN-1000252), Dual Index Kit TT Set A, 96 rxns(Cat #: PN-1000215), 5’ Feature Barcode Kit, 16 rxns (Cat #: PN-1000256), and Dual Index Kit TN Set A, 96 rxns (Cat #: PN-1000250).The concentration of each library was accurately determined through qPCR using the KAPA library Quantification Kit according to the manufacturer’s protocol (KAPA Biosystems/Roche) to achieve the desired cluster counts for the Illumina NovaSeq6000 instrument. Normalized libraries were sequenced on a NovaSeq6000 S4 Flow Cell using the XP workflow and a 151×10×10×151 sequencing recipe according to the manufacturer’s protocol. A median sequencing depth of 50,000 reads/cell was targeted for each Gene Expression library, and 5000 reads/cell for each V(D)J and Feature library.

### Single-cell RNA sequencing data analysis

Libraries were processed using Cell Ranger (v7.1.0). Low-quality barcodes and UMIs were filtered and mapped to the mouse genome (mm10). Both datasets were further aggregated using the Cell Ranger aggr pipeline (v7.1.0). The Cell Ranger aggr pipeline automatically equalizes the average read depth between samples. The gene expression from both datasets were filtered, normalized, and clustered, and the resulting Cloupe file was created and imported into the Loupe Browser for further analyses and visualization. To further define the unique molecular features of the cell population in the study, the feature barcode approach was employed to quantify each feature in each cell. The expression levels of cell surface proteins were measured via an antibody and antigen-multimer staining assay using TotalSeq-C. The Cell Ranger (v7.1.0) pipeline outputs the feature barcode counts for each cell barcode. Specific antibody detection and filtering were performed as follows: the log2 count of target ≥10, while the other log2 counts ≤6. PCA was performed to reduce the dimensionality of the dataset to its most important features, and the principal components were visualized by UMAP plots. The final analysis excluded cells with any expression of *Cd19*, *Lyz2*, *Adgre1*, *Flt3*, *Ncr1*, *Cd8b1*, *Cd79a*, *Cd79b*, *Tcrg-C1*, *Xcr1*, *Sirpa*, *Cd68*, *Fcgr1*, *Ly6g*, *Ly6c1*, and *H2-Ab1* transcripts. The differentially expressed genes between clusters or libraries were identified using the default algorithms. Bonferroni-adjusted p-values were used to determine significance at an FDR < 0.05. GSEA was performed using Phantasus^92^ (https://artyomovlab.wustl.edu/phantasus/).

### β-cell granule isolation

Human islets from de-identified, non-diabetic donors were purchased from Prodo Laboratory. Primary mouse islets were isolated as described above. Both human and mouse islets were hand-picked and dispersed using a non-enzymatic dispersion solution (Sigma-Aldrich, Cat #: C5914) for 3 min at 37°C. Islet cell suspensions (MIN6 cells, dispersed human and mouse islet cells) were then washed and resuspended in 1.0 mL PBS. Cell lysis was accomplished by passing islet cells through a Cell Homogenizer (Isobiotec) 5 times using the 10 μm clearance ball bearing to shear the cells. Cell lysate was spun for 10 min at 1,000 g, 4°C to pellet cell debris. The supernatant was transferred to a new tube followed by a repeat of the 1000 g spin. Pellets from both spins were discarded, and the supernatants were spun for 10 min at 5,000 g, 4°C. The supernatant was transferred to a new tube, and the pellet was retained. The 5,000 g spin was repeated, and the supernatant was transferred to a clean tube. Pellets from both 5,000 g spins were combined into one tube (as the crinosome fraction) and suspended in media. The supernatant was centrifuged for 20 min at 15,000 g, and the pellet was retained. The supernatant was transferred to a new tube and was centrifuged for 30 min at 25,000 g, 4°C. The pellet (as the DCG fraction) was suspended in medium. Acid phosphatase activity in the granule contents was assessed using an ELISA-based acid phosphatase assay kit (MilliporeSigma, Cat #: CS0740). Total protein levels in the granule contents were measured by a Micro BCA protein assay kit (Pierce, Cat #: 23225).

### Granule-based antigen presentation (GAP) assay

Subcellular fractions isolated by 5,000 g (5k), 15,000 (15k), or 25,000 g (25k) spins (from MIN6 cells, B6 mouse islets, and human islets) were offered to the C3.g7 APC line (B cell lymphoma expressing I-A^g7^; 5 × 10^4^/well) in a 96-well culture plate. After incubation for 2 hours, the CD4^+^ T cell hybridomas (5 × 10^4^/well) were added. After an overnight incubation, T cell responses were assessed by measuring IL-2 production in the culture supernatants using ELISA.

### β-cell and DC treatment

For experiments involving reducing agents, islet cell suspensions (MIN6 cells, dispersed human and mouse islet cells) were treated with 100 µM N-acetyl cysteine (NAC), glutathione (Sigma Cat# G4376), or TUDCA (Millipore, Cat# 580549) for 16 hours. For experiments involving ER stressors, islet cell suspensions were treated for 2 hours with either 1 µM thapsigargin (Sigma, Cat# T9033) or 1 µg/mL tunicamycin (Sigma, Cat# T7765). In experiments involving reducers and ER stressors, islet cell suspensions were treated with 100 µM NAC, glutathione, or TUDCA for 16 hours followed by addition of thapsigargin or tunicamycin for 2 hours. In all experiments, cells were washed extensively before granule isolation and presentation.

For treating human islets with cytokines, whole human islets received from Prodo Laboratories were cultured in CMRL 1066 media with 10% FBS, 1% Na Pyruvate, 1% Glutamine, 1% NEAA, and Pen/Strep. Islet cells were dispersed, collected by centrifugation, washed with media, and counted. Cells were split into 5 equal aliquots (∼10^6^ each) treated with vehicle, 50 ng/mL human TNFα (PeproTech, Cat #300-01A), 50 ng/mL human IL-1β (PeproTech, Cat #200-01B), 50 ng/mL human IFNγ (PeproTech, Cat #300-02), or all three cytokines combined. Cells were incubated for 16 hours at 37°C, washed twice with PBS, and resuspended for granule isolation.

For mouse DC antigen presentation, splenic CD11c^+^ cells were purified from NOD mice given recombinant Flt-3L-Ig (BioXcell, Cat #BE0342) using Miltenyi Beads (Miltenyi, Cat# 130-125-835). In microfuge tubes, CD11c^+^ DCs (∼1.5×10^6^ cells) were pulsed with 25 µM InsB_9-23_ peptide for 1 hour, and then were left untreated or treated with the following mouse cytokines (100 ng/mL): TNFα (PeproTech, Cat # 315-01A), IL-1β (PeproTech, Cat # 211-11B), IFNγ (PeproTech, Cat # 315-05), IL-4 (PeproTech, Cat # 214-14), IL-6 (PeproTech, Cat # 216-16), and IL-12 (PeproTech, Cat # 210-12) for 16 hours. DCs were collected, washed with PBS, and used for antigen presentation assay.

### Soluble crinosome and lymph peptidomes and DC MHC-II peptidome

Crinosome subcellular fractions were frozen at -80°C and thawed at 37°C for five cycles to release the contents of granules. Lymph samples were collected by cannulation from 6-week-old female NOD mice with or without glucose injection as described previously^93^. For both crinosome and lymph samples, a complete protease inhibitor cocktail was added, and they were concentrated by speed vac to a volume of ∼100 μl. C18 Spin Columns (Pierce, Cat #: 89870) were used to clean up the released proteins/peptides according to the manufacturer’s instructions. Samples were eluted in 0.1% formic acid (ThermoFisher, Cat #: A118-500)/95% acetonitrile (Burdick & Jackson, #015-4), and then dried with a Speed-Vac.

For mouse DC MHC-II immunopeptidome analysis, mouse splenic CD11c^+^ DCs were pulsed with 25 μM InsB_9-23_ peptide for 1 hour, followed by three different treatments: overnight culture without further additions, addition of 100 ng/mL TNFα for overnight culture, or 100 μM N-acetyl-cysteine (NAC) for 1 hour followed by TNFα for overnight culture in the presence of peptide/NAC. Cells were then collected, washed, and lysed for peptidome processing. The Mouse DC MHC-II peptidomes were isolated by a procedure similar to the human PBMC peptidomes, except that the anti-I-A^g7^ antibody conjugated to Sepharose was used.

### Mass Spectrometry

A Dionex UltiMate 3000 system (Thermo Scientific) was coupled to an Orbitrap Fusion Lumos (Thermo Scientific) through an EASY-Spray ion source (Thermo Scientific). Peptide samples were reconstituted in 2% acetonitrile (ACN)/0.1% formic acid and loaded (15 μL/minutes for 3 minutes) onto a trap column (100 μm × 2 cm, 5 μm Acclaim PepMap 100 C18, 50 °C), then eluted (200 nL/minutes) onto an EASY-Spray PepMap RSLC C18 column (2 μm, 50 cm × 75 μm ID, 50 °C, Thermo Scientific) and separated with the following gradient, all % Buffer B (0.1% formic acid in ACN): 0-110 minutes, 2-22%; 110-120 minutes, 22-35%; 120-130 minutes, 35-95%; 130-150 minutes, isocratic at 95%; 150-151 minutes, 95-2%, 151-171 minutes, isocratic at 2%. Spray voltage was 1,600-1,800 V, ion transfer tube temperature was 275 °C, and RF lens was 30%. Mass spectrometry scans were acquired in profile mode and MS/MS scans in centroid mode, for ions with charge states 2-5, with a cycle time of 1.5 s. For HCD, mass spectra were recorded from 375-1500 Da at 120K resolution (at m/z = 200), and MS/MS was triggered above a threshold of 2.0 × 10^4^, with quadrupole isolation (0.7 Da) at 15K resolution, and collision energy of 30%. Dynamic exclusion was used (60 s).

### Mass spectrometry data analysis

Processing and database searching were performed with PEAKS Studio Xpro (version 10.6, build 20201221, Bioinformatics Solutions Inc.). For crinosomes from MIN6 cells, files were searched against a Uniprot-Mouse database (Jan. 2022; 22,101 entries) with no enzyme specificity and carbamidomethylation (C), oxidation (M), and deamidation (NQ) as variable modifications. Parent mass tolerance was 10 ppm (20 ppm for MIN6 experiment), and fragment ion tolerance was 0.02 Da. The Common Repository for Adventitious Proteins database (www.thegpm.org/crap/) was used for contaminant identification, and FDR estimation was enabled. Subsequent PEAKS PTM and SPIDER searches were used to identify PTMs and SAVs, respectively. SPIDER search results, which contained both PTMs and SAVs, were filtered at a 1% FDR at the peptide level and exported for further analysis.

To further verify Cys→Ser transformations specificity, the MIN6 data files were searched against the Uniprot-Mouse database in which all cysteine residues were converted to serine using a custom Python script. Tolerances were set as above, with methionine oxidation and deamidation as variable modifications. For verification of Cys→Ser transformations in MIN6 cells and for crinosomes from mouse islets, database searching was performed with a Uniprot-Mouse database appended with Cys→Ser variants of β-cell proteins insulin 1, insulin 2, chromogranin-A, islet amyloid polypeptide, and secretogranin using the same parameters as above. For crinosomes from human islets, searches were performed against a Uniprot-Human database appended with Cys→Ser variants of β-cell proteins insulin, chromogranin-A, islet amyloid polypeptide, and secretogranins.

### Quantification of PTMs and SAVs

The PSM results from the initial SPIDER search were used for this analysis. A working list was obtained by compiling a list of all PTMs and SAVs by peptide sequence and source data file. Using the COUNTIFS function in Excel, totals were obtained for each PTM or SAV by data file. Results for fractions were then summed to obtain totals by sample, and these numbers were exported to GraphPad Prism for statistical analysis. For peptide-level quantification, peak areas from PEAKS were used. Each area was log2 transformed; a value of 2 was used in the event of missing values for peak area.

### HLA-DQ8 tetramer preparation and staining of Human PBMCs

Biotinylated monomers of HLA-DQ8 (HLA-DQA1*0301 and HLA-DQB1*0302), complexed with the insulin B chain peptide (InsB_12-20_ or InsB_12-20_(C19S)), were produced in-house^28^. The biotinylated monomers were purified through a two-step process: first by Ni-NTA affinity chromatography, followed by gel filtration using a Sephacryl-300 HR column (Cytiva). Aliquots were flash-frozen and stored at −80°C until use. The biotinylated monomers were tetramerized by incubating with streptavidin-PE or streptavidin-APC at a 5:1 molar ratio of biotinylated monomers to labeled streptavidin overnight at room temperature. PBMCs were washed twice with PBS and incubated with the APC- or PE-labeled tetramers at a final concentration of 2.5 µg/100 µL for 1 hour at room temperature. After incubation, the cells were washed twice with MACS buffer, then incubated with 20 µL of anti-APC and anti-PE microbeads in 100 µL of MACS buffer at 4°C for 20 minutes. The cells were then washed twice with FACS buffer (1% BSA in 1X PBS) and used for flow cytometry analysis.

### Spectral flow cytometry analysis

Cryopreserved PBMCs were thawed and washed twice with PBS; cell viability was above 90%. Following incubation of PBMC with PE and APC labeled tetramers, cells were stained with an anti-human FcR blocking antibody (human TruStain FcX blocking solution, Biolegend) for 10 minutes at 4°C. To maintain the resolution of positive and negative signals in a multi-color panel, we performed sequential staining of the following markers. PBMCs were first stained with anti-CXCR3 and anti-CCR4 antibodies for 5 minutes, then with anti-KLRG1 and anti-CD25 for 5 minutes at 37°C. The surface antibody cocktail was added directly and incubated for an additional 30 minutes at room temperature. Cells were stained with Live/Dead Blue viability dye (1:1000) for 25 minutes at 4°C. Cells were washed twice, and the flow cytometry data were acquired using Cytek Aurora cytometer (Cytek Biosciences) equipped with 5 lasers, followed by spectral unmixing using SpectroFlo software (Cytek Biosciences). Traditional flow cytometry of unmixed data was analyzed using FlowJo v10.9.

High-dimensional analysis of spectral flow cytometry data was performed using DownSample V3, FlowSOM, and ClusterExplorer software in FlowJo version 10.9^94–96^. InsB_12-20_- and InsB_12-20_(C19S)-specific tetramer-binding, diagonal, and tetramer-negative T cells were gated as live, singlet, CD19^⁻^CD11b^⁻^CD3^+^CD4^+^ cells. CD4^+^ tetramer-negative T cells were downsampled to match the number of tetramer-positive CD4^+^ T cells. The FCS files for the three CD4^+^ T cell populations from all 27 participants were concatenated. The combined file was uploaded into FlowJo, and t-SNE was performed. The following markers were used to generate t-SNE plots: CD45RO, CD45RA, CCR7, CD38, CD11a, CD95, CD27, CD25, CD49d, CCR4, KLRG1, CCR6, CXCR3, HLA-DR, PD1, CD127, CD28, CD226, and CXCR5. t-SNE data were subjected to unsupervised clustering using FlowSOM to generate hierarchical consensus clusters. These clusters were further analyzed, and heat maps were generated using ClusterExplorer software and Graphpad Prism. Data visualization and analysis were performed using R (R version 4.4.3). FCS files were imported and processed using the Bioconductor package flowCore (v2.18.0). All plots were generated using the ggplot2 package (v3.5.2).

## REFERENCES

1. Stern, L. J., Clement, C., Galluzzi, L. & Santambrogio, L. Non-mutational neoantigens in disease. Nat Immunol 25, 29–40 (2024).

2. Yang, M.-L. et al. Carbonyl Posttranslational Modification Associated With Early-Onset Type 1 Diabetes Autoimmunity. Diabetes 71, 1979–1993 (2022).

3. Peakman, M. et al. Naturally processed and presented epitopes of the islet cell autoantigen IA-2 eluted from HLA-DR4. J Clin Invest 104, 1449–1457 (1999).

4. McGinty, J. W. et al. Recognition of Posttranslationally Modified GAD65 Epitopes in Subjects With Type 1 Diabetes. Diabetes 63, 3033–3040 (2014).

5. Delong, T. et al. Diabetogenic T-Cell Clones Recognize an Altered Peptide of Chromogranin A. Diabetes 61, 3239–3246 (2012).

6. Delong, T. et al. Islet Amyloid Polypeptide Is a Target Antigen for Diabetogenic CD4+ T Cells. Diabetes 60, 2325–2330 (2011).

7. Mannering, S. I. et al. The insulin A-chain epitope recognized by human T cells is posttranslationally modified. Journal of Experimental Medicine 202, 1191–1197 (2005).

8. Babon, J. A. B. et al. Analysis of self-antigen specificity of islet-infiltrating T cells from human donors with type 1 diabetes. Nat. Med. 22, 1482–1487 (2016).

9. Scotto, M. et al. Zinc transporter (ZnT)8186–194 is an immunodominant CD8+ T cell epitope in HLA-A2+ type 1 diabetic patients. Diabetologia 55, 2026–2031 (2012).

10. Marre, M. L. et al. Modifying Enzymes Are Elicited by ER Stress, Generating Epitopes That Are Selectively Recognized by CD4+ T Cells in Patients With Type 1 Diabetes. Diabetes 67, 1356–1368 (2018).

11. Azoury, M. E. et al. CD8+ T Cells Variably Recognize Native Versus Citrullinated GRP78 Epitopes in Type 1 Diabetes. Diabetes 70, 2879–2891 (2021).

12. Delong, T. et al. Pathogenic CD4 T cells in type 1 diabetes recognize epitopes formed by peptide fusion. Science 351, 711–714 (2016).

13. Pataskar, A. et al. Tryptophan depletion results in tryptophan-to-phenylalanine substitutants. Nature 603, 721–727 (2022).

14. Yang, C. et al. Arginine deprivation enriches lung cancer proteomes with cysteine by inducing arginine-to-cysteine substitutants. Molecular Cell 84, 1904–1916.e7 (2024).

15. Lichti, C. F. & Wan, X. Using mass spectrometry to identify neoantigens in autoimmune diseases: The type 1 diabetes example. Seminars in Immunology 66, 101730 (2023).

16. Zhong, J., Rao, X., Xu, J.-F., Yang, P. & Wang, C.-Y. The Role of Endoplasmic Reticulum Stress in Autoimmune-Mediated Beta-Cell Destruction in Type 1 Diabetes. Journal of Diabetes Research 2012, 238980 (2012).

17. Sahin, G. S., Lee, H. & Engin, F. An accomplice more than a mere victim: The impact of β-cell ER stress on type 1 diabetes pathogenesis. Molecular Metabolism 54, 101365 (2021).

18. Marre, M. L., James, E. A. & Piganelli, J. D. β cell ER stress and the implications for immunogenicity in type 1 diabetes. Front. Cell Dev. Biol. 3, (2015).

19. Wan, X. et al. Pancreatic islets communicate with lymphoid tissues via exocytosis of insulin peptides. Nature 560, 107–111 (2018).

20. Vomund, A. N. et al. Blood leukocytes recapitulate diabetogenic peptide-MHC-II complexes displayed in the pancreatic islets. J Exp Med 218, (2021).

21. Todd, J. A., Bell, J. I. & McDevitt, H. O. HLA-DQ beta gene contributes to susceptibility and resistance to insulin-dependent diabetes mellitus. Nature 329, 599–604 (1987).

22. Acha-Orbea, H. & McDevitt, H. O. The first external domain of the nonobese diabetic mouse class II I-A beta chain is unique. Proc. Natl. Acad. Sci. U.S.A. 84, 2435–2439 (1987).

23. Nakayama, M. et al. Prime role for an insulin epitope in the development of type 1 diabetes in NOD mice. Nature 435, 220–223 (2005).

24. Daniel, D., Gill, R. G., Schloot, N. & Wegmann, D. Epitope specificity, cytokine production profile and diabetogenic activity of insulin-specific T cell clones isolated from NOD mice. European Journal of Immunology 25, 1056–1062 (1995).

25. Michels, A. W. et al. Islet-Derived CD4 T Cells Targeting Proinsulin in Human Autoimmune Diabetes. Diabetes 66, 722–734 (2016).

26. Wan, X. et al. The MHC-II peptidome of pancreatic islets identifies key features of autoimmune peptides. Nat Immunol 21, 455–463 (2020).

27. Gioia, L. et al. Position β57 of I-Ag7 controls early anti-insulin responses in NOD mice, linking an MHC susceptibility allele to type 1 diabetes onset. Science Immunology 4, eaaw6329 (2019).

28. Sharma, S. et al. Measuring anti-islet autoimmunity in mouse and human by profiling peripheral blood antigen-specific CD4 T cells. Science Translational Medicine 15, eade3614 (2023).

29. Hu, H. et al. Crinophagic granules in pancreatic β cells contribute to mouse autoimmune diabetes by diversifying pathogenic epitope repertoire. Nat Commun 15, 8318 (2024).

30. Srivastava, N. et al. Chromogranin A Deficiency Confers Protection From Autoimmune Diabetes via Multiple Mechanisms. Diabetes 70, 2860–2870 (2021).

31. Groegler, J., Callebaut, A., James, E. A. & Delong, T. The insulin secretory granule is a hotspot for autoantigen formation in type 1 diabetes. Diabetologia 67, 1507–1516 (2024).

32. Amdare, N., Purcell, A. W. & DiLorenzo, T. P. Noncontiguous T cell epitopes in autoimmune diabetes: From mice to men and back again. Journal of Biological Chemistry 297, 100827 (2021).

33. Haataja, L. et al. Disulfide Mispairing During Proinsulin Folding in the Endoplasmic Reticulum. Diabetes 65, 1050–1060 (2016).

34. Arunagiri, A. et al. Proinsulin misfolding is an early event in the progression to type 2 diabetes. eLife 8, e44532 (2019).

35. Sun, J. et al. Proinsulin misfolding and endoplasmic reticulum stress during the development and progression of diabetes⋆. Molecular Aspects of Medicine 42, 105–118 (2015).

36. Arunagiri, A. et al. Misfolded proinsulin in the endoplasmic reticulum during development of beta cell failure in diabetes. Annals of the New York Academy of Sciences 1418, 5–19 (2018).

37. Cao, S. S. & Kaufman, R. J. Endoplasmic Reticulum Stress and Oxidative Stress in Cell Fate Decision and Human Disease. Antioxidants & Redox Signaling 21, 396–413 (2014).

38. Jeong, J. et al. Novel Oxidative Modifications in Redox-Active Cysteine Residues*. Molecular & Cellular Proteomics 10, M110.000513 (2011).

39. Delmastro, M. M. & Piganelli, J. D. Oxidative Stress and Redox Modulation Potential in Type 1 Diabetes. Clinical and Developmental Immunology 2011, 1–15 (2011).

40. Oslowski, C. M. & Urano, F. Measuring ER stress and the unfolded protein response using mammalian tissue culture system. Methods Enzymol 490, 71–92 (2011).

41. Engin, F. et al. Restoration of the Unfolded Protein Response in Pancreatic β Cells Protects Mice Against Type 1 Diabetes. Sci. Transl. Med. 5, (2013).

42. Sacco, F. et al. Glucose-regulated and drug-perturbed phosphoproteome reveals molecular mechanisms controlling insulin secretion. Nat Commun 7, 13250 (2016).

43. Ferris, S. T. et al. A Minor Subset of *Batf3*-Dependent Antigen-Presenting Cells in Islets of Langerhans Is Essential for the Development of Autoimmune Diabetes. Immunity 41, 657– 669 (2014).

44. Mohan, J. F., Petzold, S. J. & Unanue, E. R. Register shifting of an insulin peptide–MHC complex allows diabetogenic T cells to escape thymic deletion. Journal of Experimental Medicine 208, 2375–2383 (2011).

45. Stratmann, T. et al. The I-Ag7 MHC Class II Molecule Linked to Murine Diabetes Is a Promiscuous Peptide Binder1. The Journal of Immunology 165, 3214–3225 (2000).

46. Latek, R. R. et al. Structural Basis of Peptide Binding and Presentation by the Type I Diabetes-Associated MHC Class II Molecule of NOD Mice. Immunity 12, 699–710 (2000).

47. Rappazzo, C. G., Huisman, B. D. & Birnbaum, M. E. Repertoire-scale determination of class II MHC peptide binding via yeast display improves antigen prediction. Nat Commun 11, 4414 (2020).

48. Collesano, L., Łuksza, M. & Lässig, M. Energy landscapes of peptide-MHC binding. PLOS Computational Biology 20, e1012380 (2024).

49. Moon, J. J. et al. Tracking epitope-specific T cells. Nat Protoc 4, 565–581 (2009).

50. Mohan, J. F. et al. Unique autoreactive T cells recognize insulin peptides generated within the islets of Langerhans in autoimmune diabetes. Nat. Immunol. 11, 350–354 (2010).

51. Wan, X., Thomas, J. W. & Unanue, E. R. Class-switched anti-insulin antibodies originate from unconventional antigen presentation in multiple lymphoid sites. Journal of Experimental Medicine 213, 967–978 (2016).

52. Podestà, M. A. et al. Stepwise differentiation of follicular helper T cells reveals distinct developmental and functional states. Nat Commun 14, 7712 (2023).

53. Wherry, E. J. et al. Molecular Signature of CD8+ T Cell Exhaustion during Chronic Viral Infection. Immunity 27, 670–684 (2007).

54. Zakharov, P. N., Hu, H., Wan, X. & Unanue, E. R. Single-cell RNA sequencing of murine islets shows high cellular complexity at all stages of autoimmune diabetes. J. Exp. Med. 217, (2020).

55. Srivastava, N. et al. CXCL16-dependent scavenging of oxidized lipids by islet macrophages promotes differentiation of pathogenic CD8+ T cells in diabetic autoimmunity. Immunity 0, (2024).

56. Støy, J. et al. In celebration of a century with insulin – Update of insulin gene mutations in diabetes. Molecular Metabolism 52, 101280 (2021).

57. Kägi, D. et al. TNF Receptor 1-Dependent β Cell Toxicity as an Effector Pathway in Autoimmune Diabetes1. The Journal of Immunology 162, 4598–4605 (1999).

58. Mitchell, J. S. et al. CD4+ T cells reactive to a hybrid peptide from insulin-chromogranin A adopt a distinct effector fate and are pathogenic in autoimmune diabetes. Immunity 57, 2399–2415.e8 (2024).

59. Baker, R. L. et al. CD4 T Cells Reactive to Hybrid Insulin Peptides Are Indicators of Disease Activity in the NOD Mouse. Diabetes 67, 1836–1846 (2018).

60. McLaughlin, R. J. et al. Human islets and dendritic cells generate post-translationally modified islet autoantigens. Clinical and Experimental Immunology 185, 133–140 (2016).

61. van Lummel, M. et al. Posttranslational Modification of HLA-DQ Binding Islet Autoantigens in Type 1 Diabetes. Diabetes 63, 237–247 (2013).

62. Foster, A. et al. Glutamine deamidation does not increase the immunogenicity of C-peptide in people with type 1 diabetes. Journal of Translational Autoimmunity 6, 100180 (2023).

63. Baker, R. L. et al. Hybrid Insulin Peptides Are Autoantigens in Type 1 Diabetes. Diabetes 68, 1830–1840 (2019).

64. Bhattacharjee, P., et al. Proinsulin C-peptide is a major source of HLA-DQ8 restricted hybrid insulin peptides recognized by human islet-infiltrating CD4+ T cells. PNAS Nexus 3, pgae491 (2024).

65. Callebaut, A. et al. An Insulin-Chromogranin A Hybrid Peptide Activates DR11-Restricted T Cells in Human Type 1 Diabetes. Diabetes 73, 743–750 (2024).

66. Hohenstein, A. C. et al. Novel T Cell reactivities to Hybrid Insulin Peptides in Islet Autoantibody-Positive At-Risk Subjects. Diabetes db240611 (2025) doi:10.2337/db24-0611.

67. Wiles, T. A. et al. Characterization of Human CD4 T Cells Specific for a C-Peptide/C-Peptide Hybrid Insulin Peptide. Front. Immunol. 12, (2021).

68. Corper, A. L. et al. A structural framework for deciphering the link between I-Ag7 and autoimmune diabetes. Science 288, 505–511 (2000).

69. Latek, R. R. et al. Structural basis of peptide binding and presentation by the type I diabetes-associated MHC class II molecule of NOD mice. Immunity 12, 699–710 (2000).

70. Lee, K. H., Wucherpfennig, K. W. & Wiley, D. C. Structure of a human insulin peptide-HLA-DQ8 complex and susceptibility to type 1 diabetes. Nat. Immunol. 2, 501–507 (2001).

71. Pugliese, A. Autoreactive T cells in type 1 diabetes. J Clin Invest 127, 2881–2891 (2017).

72. Morita, R. et al. Human Blood CXCR5+CD4+ T Cells Are Counterparts of T Follicular Cells and Contain Specific Subsets that Differentially Support Antibody Secretion. Immunity 34, 108–121 (2011).

73. Sage, P. T., Alvarez, D., Godec, J., Andrian, U. H. von & Sharpe, A. H. Circulating T follicular regulatory and helper cells have memory-like properties. J Clin Invest 124, 5191–5204 (2014).

74. Wendering, D. J. et al. Effector memory–type regulatory T cells display phenotypic and functional instability. Science Advances 10, eadn3470 (2024).

75. Ashley, C. W. & Baecher-Allan, C. Cutting Edge: Responder T Cells Regulate Human DR+ Effector Regulatory T Cell Activity via Granzyme B1. The Journal of Immunology 183, 4843– 4847 (2009).

76. Ferreira, R. C. et al. Cells with Treg-specific *FOXP3* demethylation but low CD25 are prevalent in autoimmunity. Journal of Autoimmunity 84, 75–86 (2017).

77. Thirawatananond, P. et al. Treg-Specific CD226 Deletion Reduces Diabetes Incidence in NOD Mice by Improving Regulatory T-Cell Stability. Diabetes 72, 1629–1640 (2023).

78. Fuhrman, C. A. et al. Divergent Phenotypes of Human Regulatory T Cells Expressing the Receptors TIGIT and CD226. J Immunol 195, 145–155 (2015).

79. Balmas, E., et al. Islet-autoreactive CD4^+^ T cells are linked with response to alefacept in type 1 diabetes. JCI Insight 8, (2023).

80. Wenzlau, J. M. et al. Mapping of a hybrid insulin peptide in the inflamed islet β-cells from NOD mice. Frontiers in Immunology 15, (2024).

81. Roep, B. O., Thomaidou, S., van Tienhoven, R. & Zaldumbide, A. Type 1 diabetes mellitus as a disease of the β-cell (do not blame the immune system?). Nat Rev Endocrinol 17, 150– 161 (2021).

82. Hayward, A. R., Shriber, M. & Sokol, R. Vitamin E supplementation reduces the incidence of diabetes but not insulitis in NOD mice. J Lab Clin Med 119, 503–507 (1992).

83. Tse, H. M. et al. NADPH oxidase deficiency regulates Th lineage commitment and modulates autoimmunity. J Immunol 185, 5247–5258 (2010).

84. Li, S. et al. Prevention of Autoimmune Diabetes in NOD Mice by Dimethyl Fumarate. Antioxidants (Basel) 10, 193 (2021).

85. Lee, H. et al. Beta Cell Dedifferentiation Induced by IRE1α Deletion Prevents Type 1 Diabetes. Cell Metab 31, 822–836.e5 (2020).

86. Argaev Frenkel, L., Rozenfeld, H., Rozenberg, K., Sampson, S. R. & Rosenzweig, T. N-Acetyl-l-Cysteine Supplement in Early Life or Adulthood Reduces Progression of Diabetes in Nonobese Diabetic Mice. Curr Dev Nutr 3, nzy097 (2019).

87. Netzer, N. et al. Innate immune and chemically triggered oxidative stress modifies translational fidelity. Nature 462, 522–526 (2009).

88. Kracht, M. J. L. et al. Autoimmunity against a defective ribosomal insulin gene product in type 1 diabetes. Nat Med 23, 501–507 (2017).

89. Evavold, B. D. & Allen, P. M. Separation of IL-4 Production from Th Cell Proliferation by an Altered T Cell Receptor Ligand. Science 252, 1308–1310 (1991).

90. Wang, Y. et al. How C-terminal additions to insulin B-chain fragments create superagonists for T cells in mouse and human type 1 diabetes. Science Immunology 4, eaav7517 (2019).

91. Wenzlau, J. M. et al. Insulin B-chain hybrid peptides are agonists for T cells reactive to insulin B:9-23 in autoimmune diabetes. Front Immunol 13, 926650 (2022).

92. Kleverov, M. et al. Phantasus, a web application for visual and interactive gene expression analysis. eLife 13, e85722 (2024).

93. Nanaware, P. P. et al. Role of the afferent lymph as an immunological conduit to analyze tissue antigenic and inflammatory load. Cell Reports 43, 114311 (2024).

94. Van Gassen, S. et al. FlowSOM: Using self-organizing maps for visualization and interpretation of cytometry data. Cytometry A 87, 636–645 (2015).

95. Quintelier, K. et al. Analyzing high-dimensional cytometry data using FlowSOM. Nat Protoc 16, 3775–3801 (2021).

96. Ujas, T. A., Obregon-Perko, V. & Stowe, A. M. A Guide on Analyzing Flow Cytometry Data Using Clustering Methods and Nonlinear Dimensionality Reduction (tSNE or UMAP). Methods Mol Biol 2616, 231–249 (2023).

