## Supplementary Figures 1-6; Supplementary Tables 1, 3, 4, 5 for "A microenvironment-driven, HLA-II-associated insulin neoantigen elicits persistent memory T cell activation in diabetes"

Extended Data Figure 1

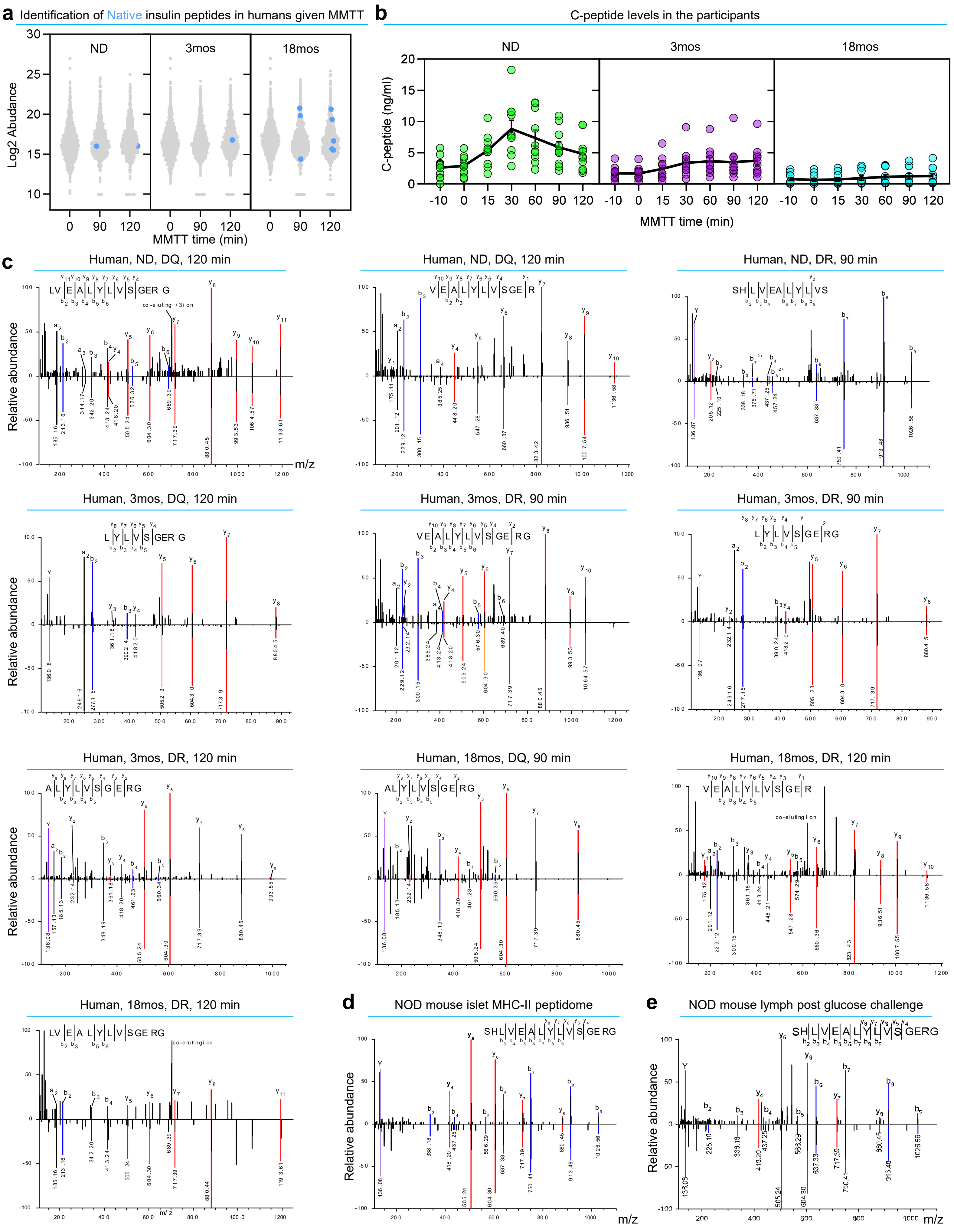

**Extended Data Figure 1. Identification and verification of C19S insulin peptides in humans and mice.**

- **a.** Quantification of native insulin B-chain peptides (each point) identified in the blood leukocyte MHC-II peptidome of NOD mice, showing increased abundance after glucose challenge. **$P < 0.01$; Wilcoxon signed-rank test.
- **b.** C-peptide levels of the three human cohorts at indicated time points before and after MMTT. Each point represents an individual participant.
- **c.** Mirror plots (total 10) showing the spectra of indicated C19S insulin B-chain peptides identified in the human HLA-DQ and HLA-DR PBMC peptidomes, all matching the synthetic standards.
- **d.** Mirror plots showing the spectra of the $InsB_{9\text{-}23}$(C19S) peptide identified in the MHC-II peptidome of pancreatic islets, which completely match the synthetic standard.
- **e.** A mirror plot showing the spectrum of the $InsB_{9\text{-}23}$(C19S) peptide identified in the mouse lymph peptidome (after glucose challenge), with a complete match to its synthetic standard.

In mirror plots (**c**, **d**, **e)**, the spectra of the peptides identified in biological samples are shown in the upper panel, with the corresponding synthetic standards in the lower panel.

Extended Data Figure 2

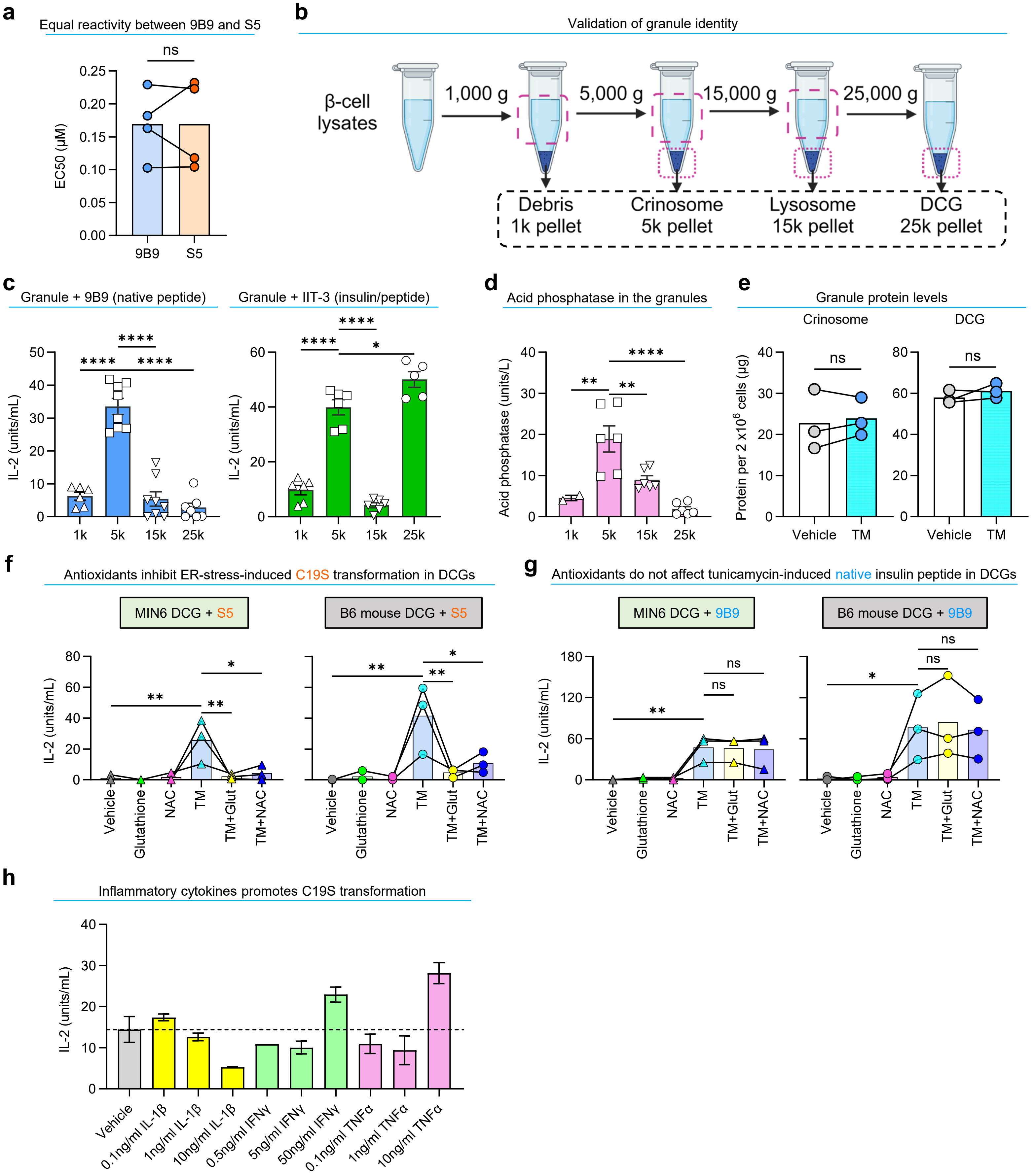

**Extended Data Figure 2. Analysis of C19S transformation in β cells under stress and inflammation.**

- **a.** Comparable reactivity of 9B9 and S5 T cells to the $InsB_{9\text{-}23}$ and $InsB_{9\text{-}23}$(C19S) peptides, respectively. Data show the peptide concentrations required to reach half-maximal responses (EC50) for each T cell.
- **b.** Schematic of the isolation and verification of individual subcellular fractions in β cells.
- **c.** GAP assay showing the responses of the 9B9 and IIT-3 T cell hybridomas to C3.g7 APCs offered with each indicated granule fraction. 9B9 specifically recognizes the native $InsB_{12\text{-}20}$ epitope, which is only presented when APCs are offered free $InsB_{9\text{-}23}$ peptide but not intact insulin, while IIT-3 recognizes the native $InsB_{13\text{-}21}$ epitope, which can be presented when APCs process both free $InsB_{9\text{-}23}$ and intact insulin. This distinction helps determine whether a given fraction contains degraded insulin peptide fragments (e.g., crinosomes) or intact insulin (e.g., DCGs).
- **d.** Acid phosphatase levels in each indicated granule fractions from MIN6 β cells.
- **e.** Total protein levels in the crinosome and DCG fractions from MIN6 treated with vehicle or tunicamycin.
- **f.** Responses of the S5 T cell to the DCG fraction isolated from MIN6 cells and B6 mouse islets following indicated treatments.
- **g.** Responses of the 9B9 T cell to the DCG fraction isolated from MIN6 cells and B6 mouse islets following indicated treatments.
- **h.** A pilot screen for C19S transformation in MIN6 cells exposed to different inflammatory cytokines using an antigen transfer assay. MIN6 cells were treated with indicated concentrations of cytokines for 16 hours. Following washing, equal numbers of cytokine-treated MIN6 cells were co-cultured with C3.g7 APCs in high glucose media (25 mM), a condition known to facilitate the exocytosis of insulin peptides from β cells, which can be acquired by C3.g7 cells for presentation. S5 T cells were then added to probe the formation of C19S insulin peptides.

Data (mean ± SEM) show results from multiple independent experiments (each point). ns, not significant; $^{*}P < 0.05$; $^{**}P < 0.01$; $^{***}P < 0.001$; $^{****}P < 0.0001$; two-tailed paired t test (**a, e**); one-way ANOVA (**c, d**); repeated measures one-way ANOVA (**f, g**).

Extended Data Figure 3

**a** Synthetic peptides used in this study

| Peptide | Sequence |
|---|---|
| InsB$_{9-23}$ | SHLVEALYLVCGERG |
| InsB$_{9-23}$(C19S) | SHLVEALYLVSGERG |
| Nested InsB$_{12-20}$ | TEGVEALYLVCGGGS |
| Nested InsB$_{12-20}$(C19S) | TEGVEALYLVSGGGS |
| Nested InsB$_{12-20}$(C19S) (K20) | TEGVEALYLVSKGGS |
| Nested InsB$_{12-20}$(C19A) | TEGVEALYLVAGGGS |
| Nested InsB$_{13-21}$ | TEGEALYLVCGEGGS |
| Nested InsB$_{13-21}$(C19S) | TEGEALYLVSGEGGS |
| Nested InsB$_{13-21}$(C19S) (K21) | TEGEALYLVSGKGGS |
| g7-MIME | GKKVATTVHAGYG |
| HEL$_{11-25}$ | AMKRHGLDNYRGYSL |

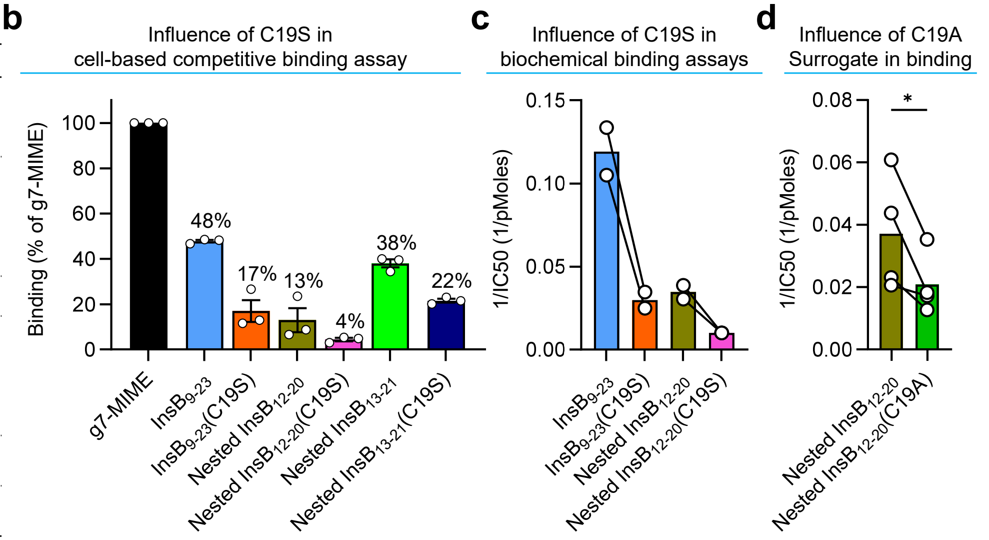


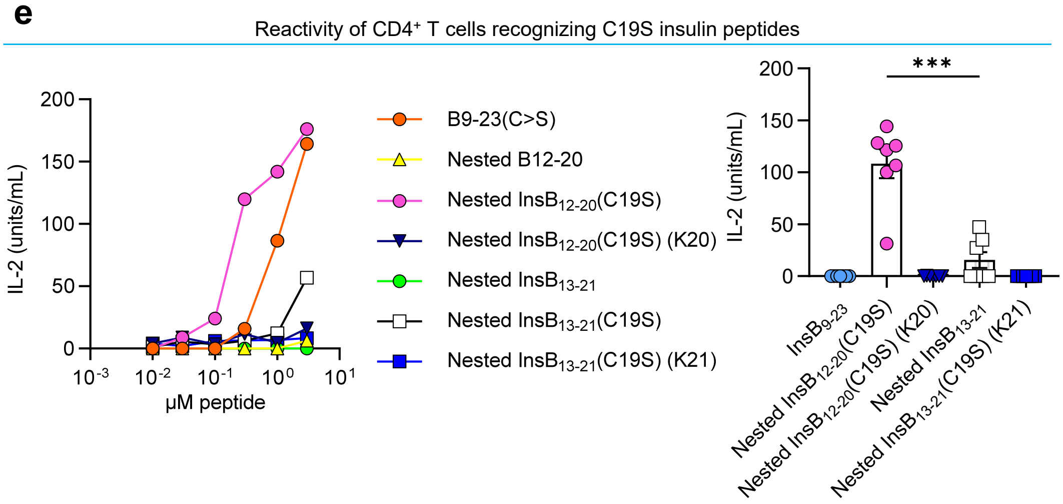


**f** I-A$^{g7}$-based tetramers used in NOD mice

| Tetramer | MHC-II | Epitope |
|---|---|---|
| InsB13-21:I-A$^{g7}$ | I-A$^{g7}$ | EALYLVCGE |
| InsB13-21(C19S):I-A$^{g7}$ | I-A$^{g7}$ | EALYLVSGE |
| InsB12-20:I-A$^{g7}$ | I-A$^{g7}$ | VEALYLVCG |
| InsB12-20(C19S):I-A$^{g7}$ | I-A$^{g7}$ | VEALYLVSG |
| InsB12-20(C19A):I-A$^{g7}$ | I-A$^{g7}$ | VEALYLVAG |

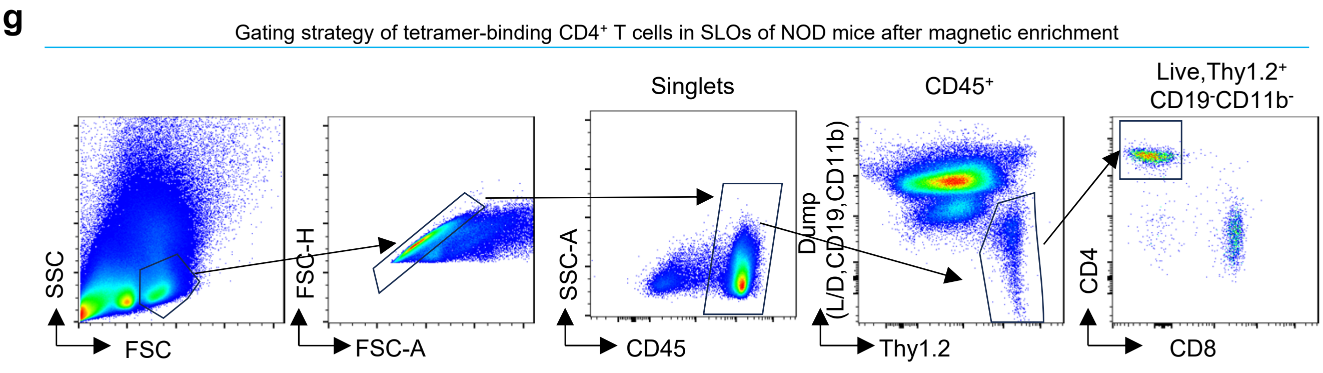


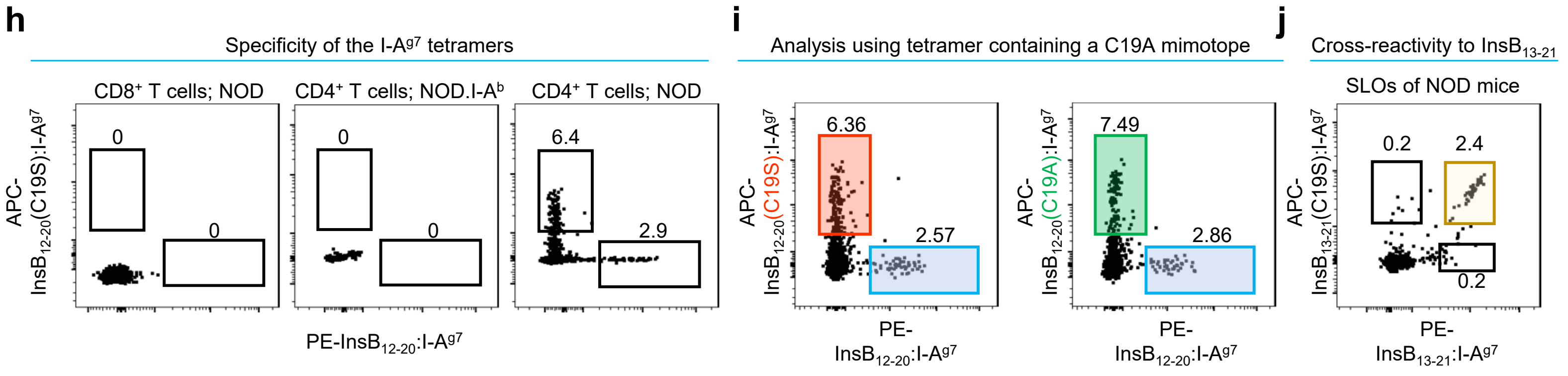

**Extended Data Figure 3. C19S diminishes MHC-II binding but is recognized by register-specific CD4$^+$ T cells.**

- **a.** A table showing sequences of synthetic peptides used in experiments in **b**, **c**, **d**.
- **b.** Relative binding affinity of indicated peptides to I-A$^{g7}$ measured by a cell-based binding assay, which evaluates the effects of indicated competitor peptides in inhibiting the binding of the hen egg lysozyme 11-25 (HEL11-25) peptide to C3.g7 cells (see Methods). Data (mean ± SEM) summarize results from three independent experiments (each point).
- **c.** Biochemical competitive binding assay showing relative binding strengths of indicated peptides to soluble I-A$^{g7}$ molecules. Data show results from two independent experiments (each point).
- **d.** Biochemical binding assay comparing I-A$^{g7}$ binding of InsB$_{12\text{-}20}$ and InsB$_{12\text{-}20}$(C19A) peptides across four independent experiments. Bars show mean ± SEM; *$P < 0.05$ by two-tailed paired t-test.
- **e.** Responses of a representative InsB$_{9\text{-}23}$(C19S)-reactive CD4$^+$ T cell hybridoma (left) to C3.g7 cells pulsed with indicated synthetic peptides. The quantification (right) summarizes results (mean ± SEM) from multiple hybridomas (each point) responding to 0.3 μM peptides. ***$P<0.001$; two-tailed paired t test.
- **f.** A table summarizing I-A$^{g7}$-based tetramers containing native and C19S insulin peptide registers used in mouse T cell studies.
- **g.** Gating strategy for tetramer-binding CD4$^+$ T cells in SLOs of NOD mice.
- **h.** Flow cytometry analysis showing co-staining with InsB$_{12\text{-}20}$:I-A$^{g7}$ and InsB$_{12\text{-}20}$(C19S):I-A$^{g7}$ tetramers in CD4$^+$ or CD8$^+$ T cells from regular NOD mice as well as in CD4$^+$ T cells from NOD.I-A$^b$ mice.
- **i.** Tetramer staining of CD4$^+$ T cells from secondary lymphoid organs of the same 10-week-old female NOD mice, comparing the InsB$_{12\text{-}20}$(C19S):I-A$^{g7}$ and InsB$_{12\text{-}20}$(C19A):I-A$^{g7}$ tetramers to the native InsB$_{12\text{-}20}$:I-A$^{g7}$ tetramer. Numbers indicate the percentage of tetramer-positive cells among CD4$^+$ T cells after magnetic enrichment. Data are representative of three independent experiments with reproducible results.
- **j.** Flow cytometry analysis showing co-staining with InsB$_{13\text{-}21}$:I-A$^{g7}$ and InsB$_{13\text{-}21}$(C19S):I-A$^{g7}$ tetramers identifying a largely overlapping T cell population.

Extended Data Figure 4

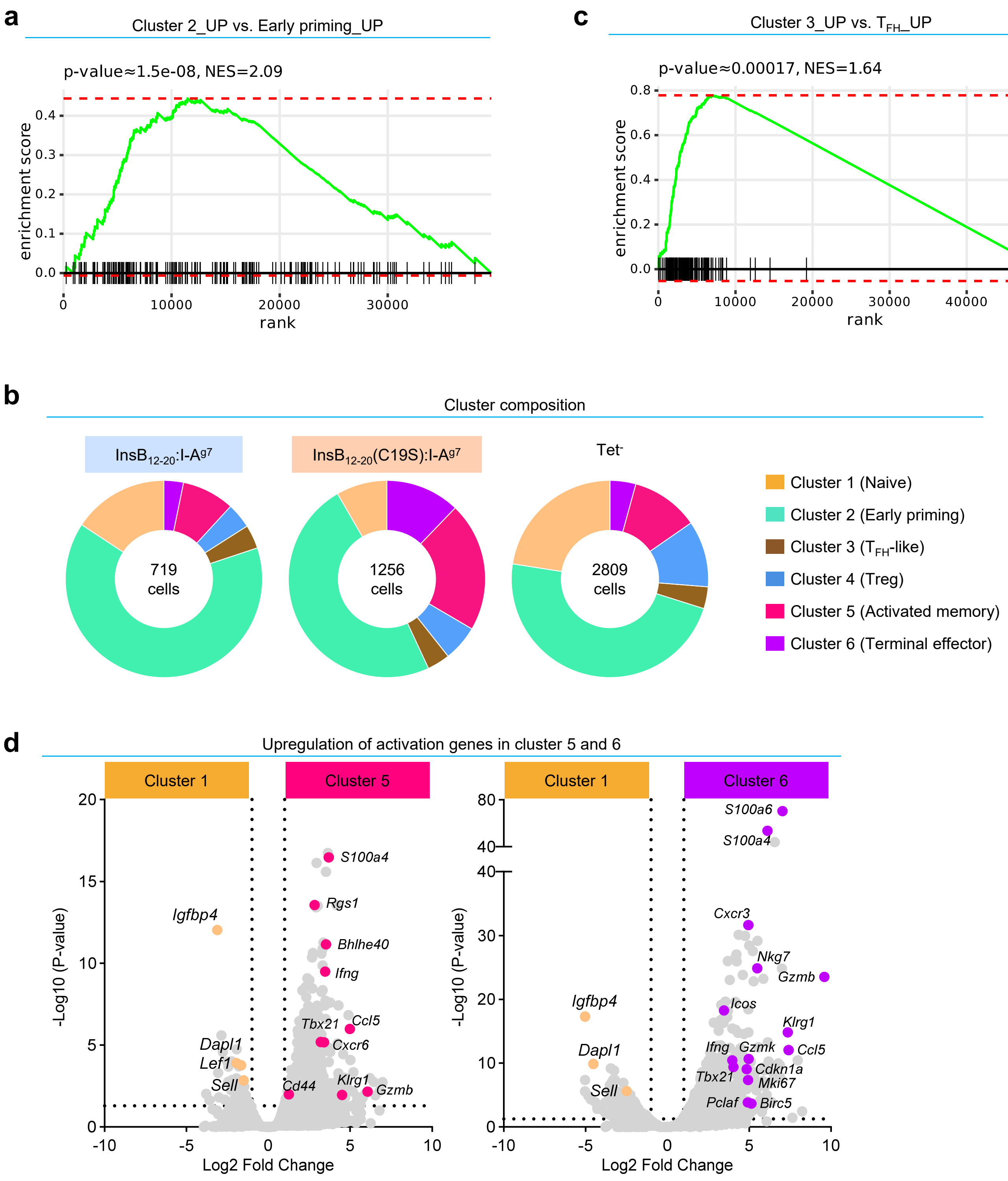


**e**

CDR3 sequences of three most expanded clones of C19S-specific $CD4^+$ T cells

| | CDR3α | CDR3β |
|---|---|---|
| Clonotype 1 | CAASGAGTGSNRLTF | CASSQETGGQAPLF |
| Clonotype 2 | CAASNTGKLTF | CASSQENTYTGQLYF |
| Clonotype 3 | CAASASGGSNYKLTF | CASSQDRGGGNTLYF |

**Extended Data Figure 4. scRNA-seq analysis of $InsB_{12\text{-}20}$- and $InsB_{12\text{-}20}$(C19S)-specific $CD4^{+}$ T cells in NOD mice.**

- **a.** GSEA showing enrichment of cluster 2 for early priming signatures from dataset GSE114824.
- **b.** Pie charts depicting the distribution of indicated clusters among $InsB_{12\text{-}20}$-specific, $InsB_{12\text{-}20}$(C19S)-specific, and $Tet^{-}$ $CD4^{+}$ T cells.
- **c.** GSEA showing enrichment of cluster 3 for $T_{FH}$ cell signatures from dataset GSE225724.
- **d.** Volcano plots showing the high expression of effector T cell genes in cluster 5 and 6 relative to the naïve T cell cluster 1. All indicated genes were significantly and differentially expressed (adjusted $P < 0.05$).
- **e.** CDR3α and CDR3β amino acid sequences of the three most expanded clonotypes among $InsB_{12\text{-}20}$(C19S):I-$A^{g7}$ tetramer-binding $CD4^{+}$ T cells.

Extended Data Figure 5

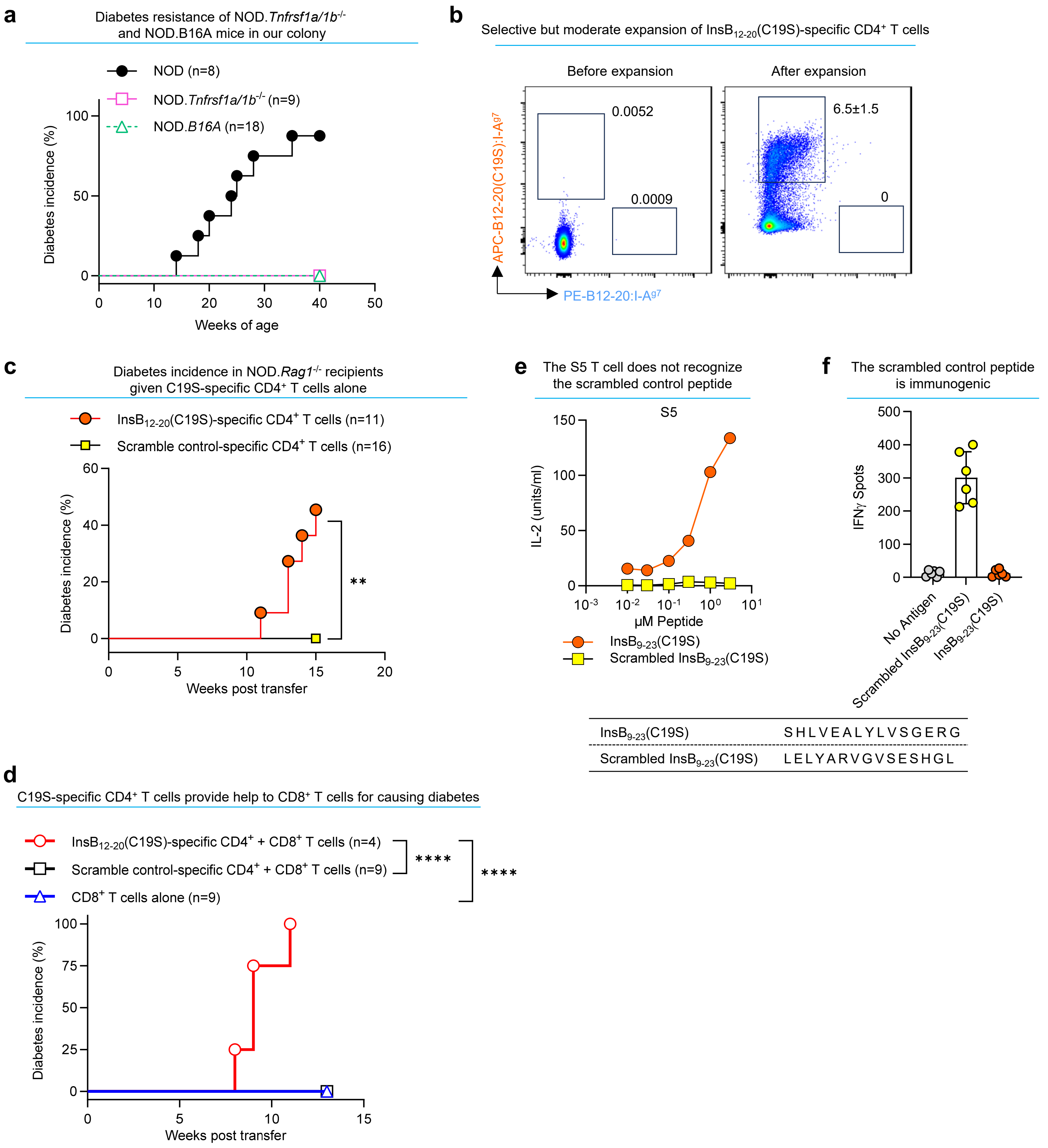

**Extended Data Figure 5. $InsB_{12-20}$(C19S)-specific $CD4^+$ T cells mediate islet inflammation and coordinate with $CD8^+$ T cells to drive diabetes development.**

- **a.** Diabetes incidence of female WT NOD, NOD.*Tnfrsf1a/1b*$^{-/-}$, and NOD.*B16A* mice in our colony.
- **b.** Flow cytometry analysis of $InsB_{12-20}$-specific and $InsB_{12-20}$(C19S)-specific tetramer-binding $CD4^+$ T cells before and after enrichment with the nested, weak-binding $InsB_{12-20}$(C19S) peptide. The data show a mild level of enrichment (see Methods) and are representative of four independent experiments.
- **c.** Diabetes incidence of NOD.*Rag1*$^{-/-}$ mice transferred with $CD4^+$ T cells enriched with the $InsB_{12-20}$(C19S) or the scrambled control peptide. $^{**}P < 0.01$; log-rank (Mantel-Cox) test.
- **d.** Diabetes incidence of NOD.*Rag1*$^{-/-}$ recipients of indicated combinations of $CD4^+$ and $CD8^+$ T cells. $^{****}P < 0.0001$; log-rank (Mantel-Cox) test.
- **e.** Reactivity of the S5 T cell to the $InsB_{9-23}$(C19S) peptide and its scrambled version.
- **f.** ELISPOT assay showing T cell responses (IFNγ production) in draining lymph node cells from NOD mice immunized with the scrambled control peptide upon antigen recall with the indicated peptides.

Extended Data Figure 6

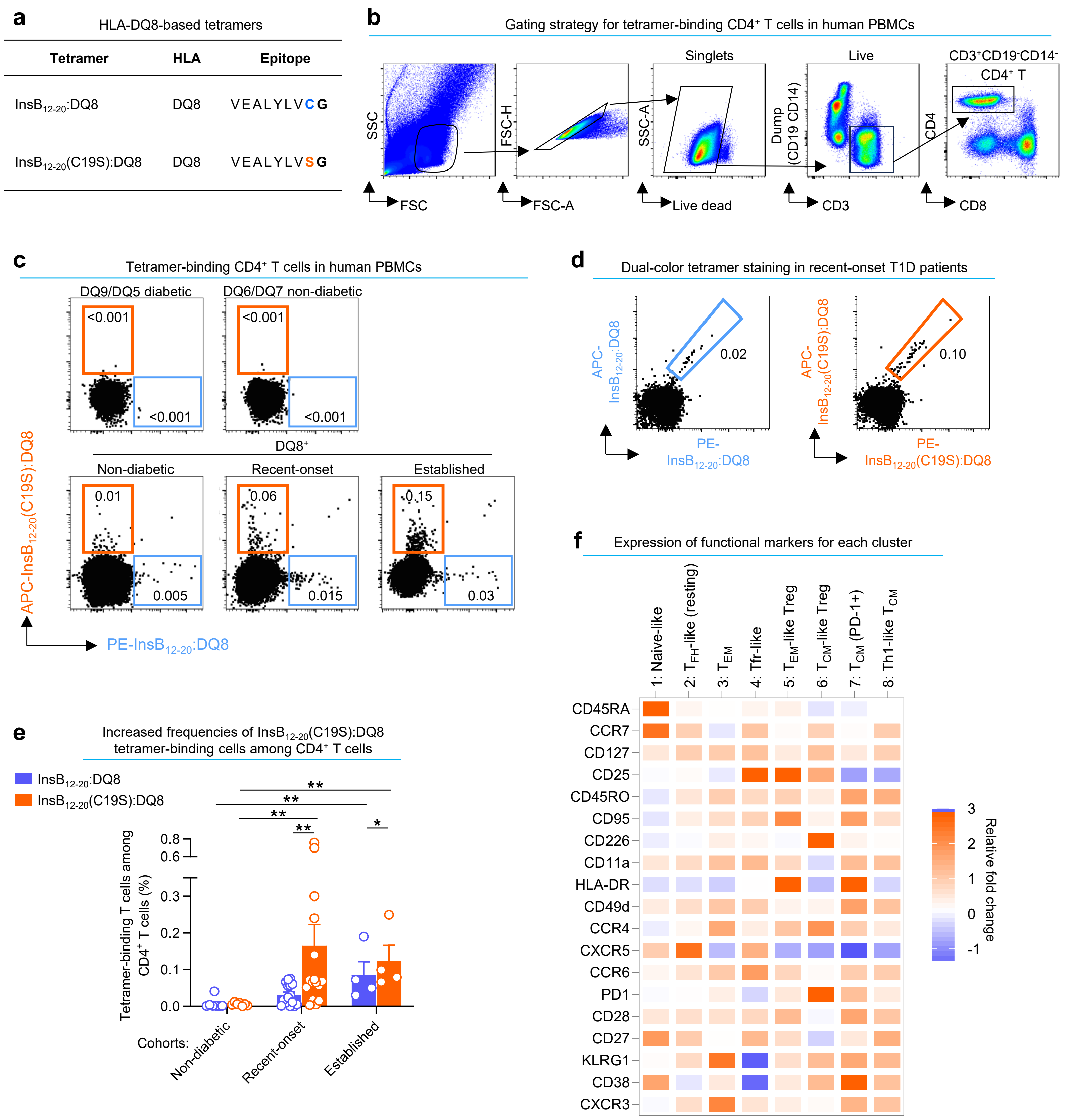

**Extended Data Figure 6. Analysis of DQ8-restricted, $InsB_{12-20}$- and $InsB_{12-20}$(C19S)-specific $CD4^+$ T cells in T1D patients.**

- **a.** A table summarizing the HLA-DQ8-based tetramers used in this study.
- **b.** Gating strategy for tetramer-binding $CD4^+$ T cells in human PBMCs without magnetic enrichment by spectral flow cytometry analysis.
- **c.** Flow cytometry analysis showing co-staining of indicated human PBMC samples with the PE-conjugated $InsB_{12-20}$:DQ8 and APC-conjugated $InsB_{12-20}$(C19S):DQ8 tetramers. FACS plots of three individuals of the non-diabetic, recent-onset, and established T1D cohorts were merged and shown.
- **d.** Dual-color tetramer staining of PBMCs from recent-onset T1D patients using $InsB_{12-20}$:DQ8 and $InsB_{12-20}$(C19S):DQ8 tetramers simultaneously conjugated with PE and APC. FACS plots from three individual recent-onset patients were merged and shown.
- **e.** Quantification of the frequencies of indicated DQ8 tetramer-binding T cells among $CD4^+$ T cells from all 27 individuals across non-diabetic, recent-onset, and established cohorts. Data summarize results from individual participants (each point) analyzed in 14 independent experiments. $^*P < 0.05$; $^{**}P < 0.01$; Mann-Whitney test.
- **f.** A heatmap showing the expression of indicated functional markers by each human T cell cluster.

**Supplementary Table 1. HLA-II haplotypes of human participants in the PBMC immunopeptidome analysis.**

| SampleID | visit | HLA-DQA1 allele1 | HLA-DQA1 allele2 | HLA-DQB1 allele1 | HLA-DQB1 allele2 | HLA-DRB1 allele1 | HLA-DRB1 allele2 | HLA-DPB1 allele1 | HLA-DPB1 allele2 | HLA-DPA1 allele1 | HLA-DPA1 allele2 | DRB1*04:01~DQB1*03:02 haplotype | DRB1*03:01~DQB1*02:01 haplotype |
|---|---|---|---|---|---|---|---|---|---|---|---|---|---|
| IPD-001 | 3mos | 01:01 | 05:01 | 02:01 | 05:01 | 01:01 | 03:01 | 01:01 | 03:01 | 01:03 | 02:01 | 0 | 1 |
| IPD-002 | 18mos | 01:01 | 05:01 | 02:01 | 05:01 | 01:03 | 03:01 | 02:01 | 04:01 | | | 0 | 1 |
| IPD-003 | 18mos | 05:01 | 05:01 | 02:01 | 02:01 | 03:01 | 03:01 | 03:01 | 04:02 | 01:03 | 01:03 | 0 | 2 |
| IPD-004 | 3mos | 02:01 | 05:01 | 02:01 | 02:02 | 03:01 | 07:01 | 04:01 | 06:01 | 01:03 | 01:03 | 0 | 1 |
| IPD-005 | 18mos | 05:01 | 05:01 | 02:01 | 02:01 | 03:01 | 03:01 | 01:01 | 04:01 | 01:03 | 02:01 | 0 | 2 |
| IPD-006 | control | 03:02 | 05:01 | 02:01 | 03:03 | 03:01 | 09:01 | 01:01 | 02:01 | 01:03 | 02:01 | 0 | 1 |
| IPD-007 | control | 01:01 | 03:03 | 03:01 | 05:01 | 01:01 | 04:01 | 04:01 | 04:02 | 01:03 | 01:03 | 0 | 0 |
| IPD-008 | 18mos | 03:03 | 05:01 | 02:01 | 03:01 | 03:01 | 04:01 | 04:01 | 04:01 | 01:03 | 01:03 | 0 | 1 |
| IPD-009 | control | 01:02 | 02:01 | 03:03 | 06:04 | 07:01 | 13:02 | 04:01 | 13:01 | 01:03 | 02:01 | 0 | 0 |
| IPD-011 | 3mos | 01:02 | 03:01 | 03:02 | 06:09 | 04:01 | 13:02 | 04:01 | 05:01 | 01:03 | 02:01 | 1 | 0 |
| IPD-012 | 3mos | 03:01 | 05:01 | 02:01 | 03:02 | 03:01 | 04:04 | 02:02 | 06:01 | 01:03 | 01:03 | 0 | 1 |
| IPD-013 | 18mos | 03:03 | 05:01 | 02:01 | 02:02 | 03:01 | 07:01 | 665:01 | 09:01 | 02:01 | 04:02 | 0 | 1 |
| IPD-014 | control | 01:02 | 01:02 | 06:02 | 06:02 | 15:03 | 15:03 | 105:01 | 11:01 | 02:01 | 03:01 | 0 | 0 |
| IPD-015 | 3mos | 01:02 | 05:01 | 02:01 | 06:04 | 03:01 | 13:02 | 04:01 | 14:01 | 01:03 | 02:01 | 0 | 1 |
| IPD-016 | 18mos | 03:01 | 05:01 | 02:01 | 03:02 | 03:01 | 04:04 | 02:01 | 04:02 | 01:03 | 01:03 | 0 | 1 |
| IPD-017 | 3mos | 03:03 | 05:01 | 02:01 | 03:02 | 03:01 | 04:05 | 02:01 | 04:01 | 01:03 | 01:03 | 0 | 1 |
| IPD-018 | control | 02:01 | 05:05 | 03:01 | 03:03 | 07:01 | 11:01 | 03:01 | 04:02 | 01:03 | 01:03 | 0 | 0 |
| IPD-019 | 18mos | 03:01 | 05:01 | 02:01 | 03:02 | 03:01 | 04:01 | 02:01 | 04:01 | 01:03 | 01:03 | 1 | 1 |
| IPD-020 | control | 03:01 | 03:03 | 03:01 | 03:02 | 04:01 | 04:01 | 02:01 | 04:01 | 01:03 | 01:03 | 1 | 0 |
| IPD-021 | control | 03:03 | 05:01 | 02:01 | 03:01 | 03:01 | 04:01 | 04:01 | 15:01 | 01:03 | 01:03 | 0 | 1 |
| IPD-022 | control | 03:01 | 05:01 | 02:01 | 03:02 | 03:01 | 04:01 | 02:01 | 04:01 | 01:03 | 01:03 | 1 | 1 |
| IPD-023 | 3mos | 01:03 | 05:01 | 02:01 | 06:03 | 03:01 | 13:01 | 04:01 | 09:01 | 01:03 | 02:01 | 0 | 1 |
| IPD-024 | 18mos | 03:01 | 03:01 | 03:02 | 03:02 | 04:01 | 04:04 | 11:01 | 11:01 | 02:01 | 02:01 | 1 | 0 |
| IPD-025 | 18mos | 02:01 | 05:01 | 02:01 | 02:02 | 03:01 | 07:01 | 04:01 | 04:01 | 01:03 | 01:03 | 0 | 1 |
| IPD-026 | control | 01:04 | 05:01 | 02:01 | 05:03 | 03:01 | 14:54 | 03:01 | 04:01 | 01:03 | 01:03 | 0 | 1 |
| IPD-027 | 3mos | 01:01 | 04:01 | 04:02 | 05:01 | 01:01 | 08:01 | 02:01 | 03:01 | 01:03 | 01:03 | 0 | 0 |
| IPD-028 | 3mos | 01:01 | 03:01 | 03:02 | 05:01 | 01:01 | 04:01 | 02:01 | 20:01 | 01:03 | 01:03 | 1 | 0 |
| IPD-029 | 3mos | 04:01 | 05:01 | 02:01 | 04:02 | 03:01 | 08:01 | 03:01 | 04:01 | 01:03 | 01:03 | 0 | 1 |
| IPD-030 | 18mos | 05:05 | 05:05 | 03:01 | 03:01 | 11:01 | 11:01 | 04:02 | 04:02 | 01:03 | 01:03 | 0 | 0 |
| IPD-031 | control | 02:01 | 05:01 | 02:01 | 03:03 | 03:01 | 07:01 | 01:01 | 04:01 | 01:03 | 02:01 | 0 | 1 |

### Supplementary Table 3. Insulin B-chain peptides with cysteine-to-serine transformations identified in crinosome peptidomes under ER stress

| Peptide | Start | End | Cysteine-to-seine | Log2 Abundance | | | | | |
|---|---|---|---|---|---|---|---|---|---|
| | | | | Vehicle_1 | Vehicle_2 | Vehicle_3 | TM_1 | TM_2 | TM_3 |
| SGPHLVEALYLVSGERG | 7 | 23 | C7, C19 | 1 | 1 | 1 | 21.94967 | 23.54911 | 23.14479 |
| SGSHLVEALYLVSGERG | 7 | 23 | C7, C19 | 1 | 1 | 1 | 1 | 17.79222 | 9.965784 |
| GSHLVEALYLVSGERG | 8 | 23 | C19 | 1 | 1 | 1 | 21.11785 | 22.56433 | 21.5644 |
| SHLVEALYLVSGERG | 9 | 23 | C19 | 24.29754 | 26.38111 | 27.41143 | 38.14328 | 38.75297 | 38.91163 |
| SHLVEALYLVSGER | 9 | 22 | C19 | 1 | 1 | 1 | 12.65212 | 9.965784 | 9.965784 |
| SHLVEALYLVSGE | 9 | 21 | C19 | 1 | 1 | 1 | 18.78143 | 19.11513 | 24.19184 |
| SHLVEALYLVSG | 9 | 20 | C19 | 1 | 1 | 1 | 24.42471 | 1 | 24.61613 |
| SHLVEALYLVS | 9 | 19 | C19 | 1 | 1 | 1 | 17.91865 | 21.44879 | 1 |
| HLVEALYLVSGERG | 10 | 23 | C19 | 1 | 1 | 19.35499 | 28.15238 | 29.81427 | 29.85325 |
| LVEALYLVSGERG | 11 | 23 | C19 | 18.19452 | 9.965784 | 16.16232 | 31.89634 | 31.15698 | 31.47518 |
| VEALYLVSGERG | 12 | 23 | C19 | 1 | 1 | 1 | 27.15518 | 27.18639 | 27.47522 |
| EALYLVSGERG | 13 | 23 | C19 | 1 | 1 | 1 | 26.24532 | 25.91332 | 26.14954 |
| ALYLVSGERG | 14 | 23 | C19 | 1 | 1 | 1 | 25.69336 | 25.13118 | 25.36182 |
| LYLVSGERG | 15 | 23 | C19 | 1 | 1 | 1 | 23.94566 | 25.76986 | 25.81617 |
| YLVSGERG | 16 | 23 | C19 | 1 | 1 | 1 | 17.18969 | 21.87725 | 26.86112 |
| LVSGERG | 17 | 23 | C19 | 1 | 1 | 1 | 1 | 29.61357 | 9.965784 |

**Supplementary Table 4. Information of human islet donors**

| | Age | Sex | BMI | HbA1C (%) | Islet viability (%) | Experiments |
|---|---|---|---|---|---|---|
| Donor A | 36 | Male | 23.2 | 5.3 | 95 | GAP assay |
| Donor B | 48 | Female | 36.39 | 5.6 | 95 | |
| Donor C | 46 | Male | 23.1 | 4.9 | 95 | |
| Donor D | 63 | Male | 21.3 | 5.5 | 95 | |
| Donor E | 40 | Male | 31.7 | 5.5 | 95 | Crinosome peptidome analysis |
| Donor F | 63 | Male | 34 | 5.1 | 95 | |

**Supplementary Table 5. HLA-II haplotypes of human participants for T cell analysis**

| Cohort | ID | DQ-SE-1 | DQ-SE-2 | DQA1-1 | DQA1-2 | DQB1-1 | DQB1-2 |
|---|---|---|---|---|---|---|---|
| Non-diabetic | **ND-001** | DQ2 | DQ8 | DQA1*03:01 | DQA1*03:03 | DQB1*02:02 | DQB1*03:02 |
| | **ND-002** | DQ8 | DQ8 | DQA1*03:01 | DQA1*03:01 | DQB1*03:02 | DQB1*03:02 |
| | **ND-003** | DQ2 | DQ8 | DQA1*03:01 | DQA1*05:01 | DQB1*03:02 | DQB1*02:01 |
| | **ND-004** | DQ2 | DQ8 | DQA1*02:01 | DQA1*03:01 | DQB1*02:02 | DQB1*03:02 |
| | **ND-005** | DQ7 | DQ8 | DQA1*03:03 | DQA1*03:03 | DQB1*03:01 | DQB1*03:02 |
| | **ND-015** | DQ8 | DQ8 | DQA1*03:01 | DQA1*03:01 | DQB1*03:02 | DQB1*03:02 |
| | **ND-017** | DQ7 | DQ8 | DQA1*03:01 | DQA1*05:01 | DQB1*03:01 | DQB1*03:02 |
| Recent-onset | **RO-002** | DQ2 | DQ8 | DQA1*03:01 | DQA1*05:01 | DQB1*02:01 | DQB1*03:02 |
| | **RO-006** | DQ8 | DQ8 | DQA1*03:01 | DQA1*03:01 | DQB1*03:02 | DQB1*03:02 |
| | **RO-007** | DQ8 | DQ6 | DQA1*01:03 | DQA1*03:01 | DQB1*03:02 | DQB1*06:03 |
| | **RO-009** | DQ2 | DQ8 | DQA1*03:01 | DQA1*05:01 | DQB1*02:01 | DQB1*03:02 |
| | **RO-012** | DQ7 | DQ8 | DQA1*03:01 | DQA1*03:01 | DQB1*03:01 | DQB1*03:02 |
| | **RO-013** | DQ2 | DQ8 | DQA1*03:02 | DQA1*05:01 | DQB1*02:01 | DQB1*03:02 |
| | **RO-020** | DQ2 | DQ8 | DQA1*03:01 | DQA1*05:01 | DQB1*02:01 | DQB1*03:02 |
| | **RO-021** | DQ8 | DQ8 | DQA1*03:01 | DQA1*03:01 | DQB1*03:02 | DQB1*03:02 |
| | **RO-027** | DQ8 | DQ8 | DQA1*03:01 | DQA1*03:01 | DQB1*03:02 | DQB1*03:02 |
| | **RO-029** | DQ8 | DQ5 | DQA1*03:01 | DQA1*01:01 | DQB1*03:02 | DQB1*05:01 |
| | **RO-030** | DQ2 | DQ8 | DQA1*03:01 | DQA1*05:01 | DQB1*02:01 | DQB1*03:02 |
| | **RO-031** | DQ8 | DQ5 | DQA1*03:01 | DQA1*01:01 | DQB1*03:02 | DQB1*05:01 |
| | **RO-036** | DQ2 | DQ8 | DQA1*03:01 | DQA1*05:01 | DQB1*02:01 | DQB1*03:02 |
| | **RO-038** | DQ8 | DQ8 | DQA1*03:01 | DQA1*03:01 | DQB1*03:02 | DQB1*03:02 |
| | **RO-040** | DQ8 | DQ6 | DQA1*01:03 | DQA1*03:01 | DQB1*03:02 | DQB1*06:03 |
| | **RO-042** | DQ8 | DQ6 | DQA1*01:03 | DQA1*03:01 | DQB1*03:02 | DQB1*06:03 |
| | **RO-048** | DQ2 | DQ8 | DQA1*03:01 | DQA1*05:01 | DQB1*02:01 | DQB1*03:02 |
| Establis hed | **ES-22** | DQ2 | DQ8 | DQA1*03:01 | DQA1*05:01 | DQB1*02:01 | DQB1*03:02 |
| | **ES-24** | DQ2 | DQ8 | DQA1*03:01 | DQA1*03:01 | DQB1*02:02 | DQB1*03:02 |
| | **ES-26** | DQ2 | DQ8 | DQA1*03:02 | DQA1*05:01 | DQB1*02:01 | DQB1*03:02 |
| | **ES-27** | DQ7 | DQ8 | DQA1*03:01 | DQA1*03:01 | DQB1*03:01 | DQB1*03:02 |
